# Beyond Consensus: Embracing Heterogeneity in Curated Neuroimaging Meta-Analysis

**DOI:** 10.1101/149567

**Authors:** Gia H. Ngo, Simon B. Eickhoff, Minh Nguyen, Gunes Sevinc, Peter T. Fox, R. Nathan Spreng, B.T. Thomas Yeo

## Abstract

Coordinate-based meta-analysis can provide important insights into mind-brain relationships. A popular approach for curated small-scale meta-analysis is activation likelihood estimation (ALE), which identifies brain regions consistently activated across a selected set of experiments, such as within a functional domain or mental disorder. ALE can also be utilized in meta-analytic co-activation modeling (MACM) to identify brain regions consistently co-activated with a seed region. Therefore, ALE aims to find consensus across experiments, treating heterogeneity across experiments as noise. However, heterogeneity within an ALE analysis of a functional domain might indicate the presence of functional sub-domains. Similarly, heterogeneity within a MACM analysis might indicate the involvement of a seed region in multiple co-activation patterns that are dependent on task contexts. Here, we demonstrate the use of the author-topic model to automatically determine if heterogeneities within ALE-type meta-analyses can be robustly explained by a small number of latent patterns. In the first application, the author-topic modeling of experiments involving self-generated thought (N = 179) revealed cognitive components fractionating the default network. In the second application, the author-topic model revealed that the left inferior frontal junction (IFJ) participated in multiple task-dependent co-activation patterns (N = 323). Furthermore, the author-topic model estimates compared favorably with spatial independent component analysis in both simulation and real data. Overall, the results suggest that the author-topic model is a flexible tool for exploring heterogeneity in ALE-type meta-analyses that might arise from functional sub-domains, mental disorder subtypes or task-dependent co-activation patterns. Code for this study is publicly available (https://github.com/ThomasYeoLab/CBIG/tree/master/stable_projects/meta-analysis/Ngo2019_AuthorTopic).

## 1 Introduction

Brain imaging experiments are often underpowered (Carp, 2012; Poline et al., 2012; Button et al., 2013). Coordinate-based meta-analysis provides an important framework for analyzing underpowered studies across different experimental conditions and analysis piplines to reveal reliable trends (Wager et al. 2003; Fox et al. 2014; Poldrack and Yarkoni, 2016). Large-scale coordinate-based meta-analyses synthesize thousands of experiments across diverse experimental designs to discover broad and general principles of brain organization and disorder (Laird et al., 2011; Poldrack et al., 2011; Crossley et al., 2014). By contrast, the vast majority of meta-analyses involve smaller number of experiments that are expertly chosen (curated) to generate consensus on specific functional domains (e.g., Binder et al. 2009), brain regions (e.g., Shackman et al., 2011) or disorders (e.g., Cortese et al., 2012).

A popular approach for smaller-scale meta-analyses is activation likelihood estimation or ALE (Laird et al., 2005; Eickhoff et al. 2009, 2012; Turkeltaub et al. 2012). ALE identifies brain regions consistently activated across neuroimaging experiments within a functional domain (Costafreda et al., 2008; Spaniol et al., 2009; Beissner et al., 2013) or within a disorder (e.g., Fitzgerald et al., 2008; Minzenberg et al., 2009; Di Martino et al., 2009). Thus, ALE treats heterogeneities across studies as noise. Consequently, ALE analysis might miss out on genuine biological heterogeneity indicative of functional sub-domains or disorder subtypes.

For example, Figure 1 (middle panel) illustrates activation foci from experiments associated with a hypothetical functional domain. These foci are generated by two latent sub-domains activating distinct, but overlapping, brain regions. Without prior knowledge of the two sub-domains from theory or previous empirical work, ALE will converge on regions commonly activated across both sub-domains (Figure 1 left panel). To get around this issue, meta-analytic studies can sub-divide experiments into hypothetical functional sub-domains before applying ALE. For example, a recent meta-analysis divided working memory experiments into verbal versus non-verbal tasks, as well as tasks involving object identity versus object locations (Rottschy et al., 2012). However, manually subdividing experiments requires prior knowledge of the sub-domains and may reinforce biases towards existing concepts. By contrast, in this study, we explored whether a previously published data-driven approach (author-topic model; Yeo et al., 2015) can help uncover heterogeneities within ALE-type meta-analyses in a bottom-up, data-driven fashion (Figure 1 right panel).

**Figure 1.**
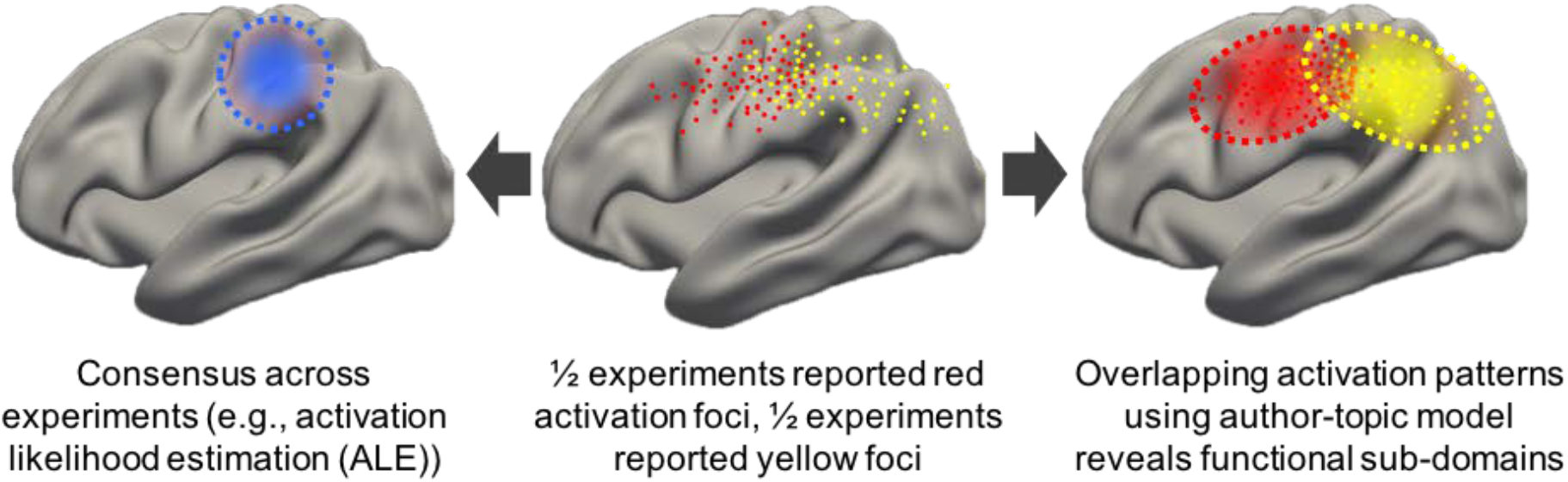
Example of heterogeneity in neuroimaging meta-analysis. The middle panel shows activation peaks reported from neuroimaging experiments within a functional domain. Half the experiments are red dots; half the experiments are yellow dots. The left panel illustrates a possible outcome of activation likelihood estimation (ALE), which converges on regions consistently activated across experiments (blue dotted circle). The right panel illustrates a possible estimate by the author-topic model (Yeo et al., 2015), which recovers overlapping patterns (red and yellow ovals) corresponding to two functional sub-domains.

### 1.1 Discovering sub-domains of self-generated thoughts

A good example in which ALE might miss out on functional sub-domains is the default network and self-generated thought (Smallwood, 2013; Andrews-Hanna et al., 2014). Self-generated thought involves associative and constructive processes that take place within an individual, and depends upon an internal representation to reconstruct or imagine a situation, understand a stimulus, or generate an answer to a question. The term “self-generated thought” serves to contrast with thoughts where the primary referent is based on immediate perceptual input. By virtue of being largely stimulus independent or task unrelated, self-generated thought has been linked with the functions of the default network (Buckner et al., 2008; Andrews-Hanna et al., 2014). Previous ALE meta-analyses have implicated the default network in many tasks involving self-generated thought, including theory of mind, narrative fiction, autobiographical memory and moral cognition (Spreng et al. 2009; Binder et al. 2009; Mar, 2011; Sevinc and Spreng, 2014).

However, many studies have suggested that the default network might be fractionated into sub-systems. For example, Andrews-Hanna and colleagues have proposed a dorsomedial prefrontal subsystem preferentially specialized for social cognition and narrative processing (Andrews-Hanna et al., 2014; Spreng & Andrews-Hanna, 2015) and a medial temporal lobe sub-system preferentially specialized for mnemonic constructive processes (Andrews-Hanna et al., 2014; Christoff et al., 2016). Both sub-systems might spatially overlap or inter-digitate across multiple brain regions (Andrews-Hanna et al., 2014; Rodrigo and Buckner, 2017), which would be challenging to ALE without assuming prior knowledge of the sub-systems (Figure 1). Furthermore, specific default network fractionation details differed across studies (Laird et al. 2009; Andrews-Hanna et al., 2010; Mayer et al. 2010; Humphreys et al., 2015; Kernbach et al., 2018), so application of the author-topic model might potentially clarify sub-systems subserving self-generated thought.

### 1.2 Discovering multiple co-activation patterns of the left inferior frontal junction (IFJ)

Another common application of ALE is meta-analytic connectivity modeling (MACM), which identifies brain regions that consistently co-activate with a particular seed region (Toro et al., 2008; Koski and Paus, 2010; Robinson et al., 2010; Eickhoff et al., 2010). The assumption is that the seed region exhibits a *single* co-activation pattern regardless of the actual task activating the seed region. However, studies have shown the existence of multiple hub regions in the brain (e.g., dorsal anterior insula, dorsal anterior cingulate cortex) that are activated across many different tasks and might adapt their connectivity pattern depending on task context (Cole et al., 2013; Uddin 2015; Bertolero et al., 2017). Thus, a seed region might be involved in *multiple* task-dependent co-activation patterns (McIntosh, 2000).

A good example in which MACM might miss out on multiple co-activation patterns is the left inferior frontal junction (IFJ; Muhle-Karbe et al., 2015). The IFJ has been implicated in many cognitive processes (Brass et al. 2005; Chikazoe et al. 2009; Asplund et al. 2010) and is a key node of the multiple-demand system (Duncan et al., 2010; Fedorenko et al., 2010). IFJ might also coordinate information among modules by adapting its connectivity patterns across different resting and task states (Cole et al., 2013; Bertolero et al., 2018). Therefore, one might expect the IFJ region to exhibit multiple co-activation patterns that are dependent on task contexts. Since ALE cannot capture heterogeneity across experiments, MACM might be insensitive to such task-dependent co-activation patterns. On the other hand, application of the author-topic model to the IFJ region might yield multiple meaningful co-activation patterns.

### 1.3 Author-topic model

In this work, we propose the use of the author-topic model to automatically make sense of heterogeneity within ALE-type meta-analyses. We have previously utilized the author-topic model (Figure 2; Yeo et al. 2015; Bertolero et al., 2015) to encode the intuitive notion that a behavioral task recruits multiple cognitive components, which are in turn supported by overlapping brain regions (Poldrack 2006; Leech et al. 2012; Barrett and Satpute, 2013). While our previous work focused on large-scale meta-analysis across many functional domains (Yeo et al. 2015; Bertolero et al., 2015), the current study focuses on heterogeneity within a functional domain (self-generated thought) or co-activation heterogeneity of a seed region (left IFJ). These applications of the author-topic model are made possible by the development of a novel inference algorithm for the author-topic model (Ngo et al., 2016) that is sufficiently robust for smaller-scale meta-analyses.

**Figure 2.**
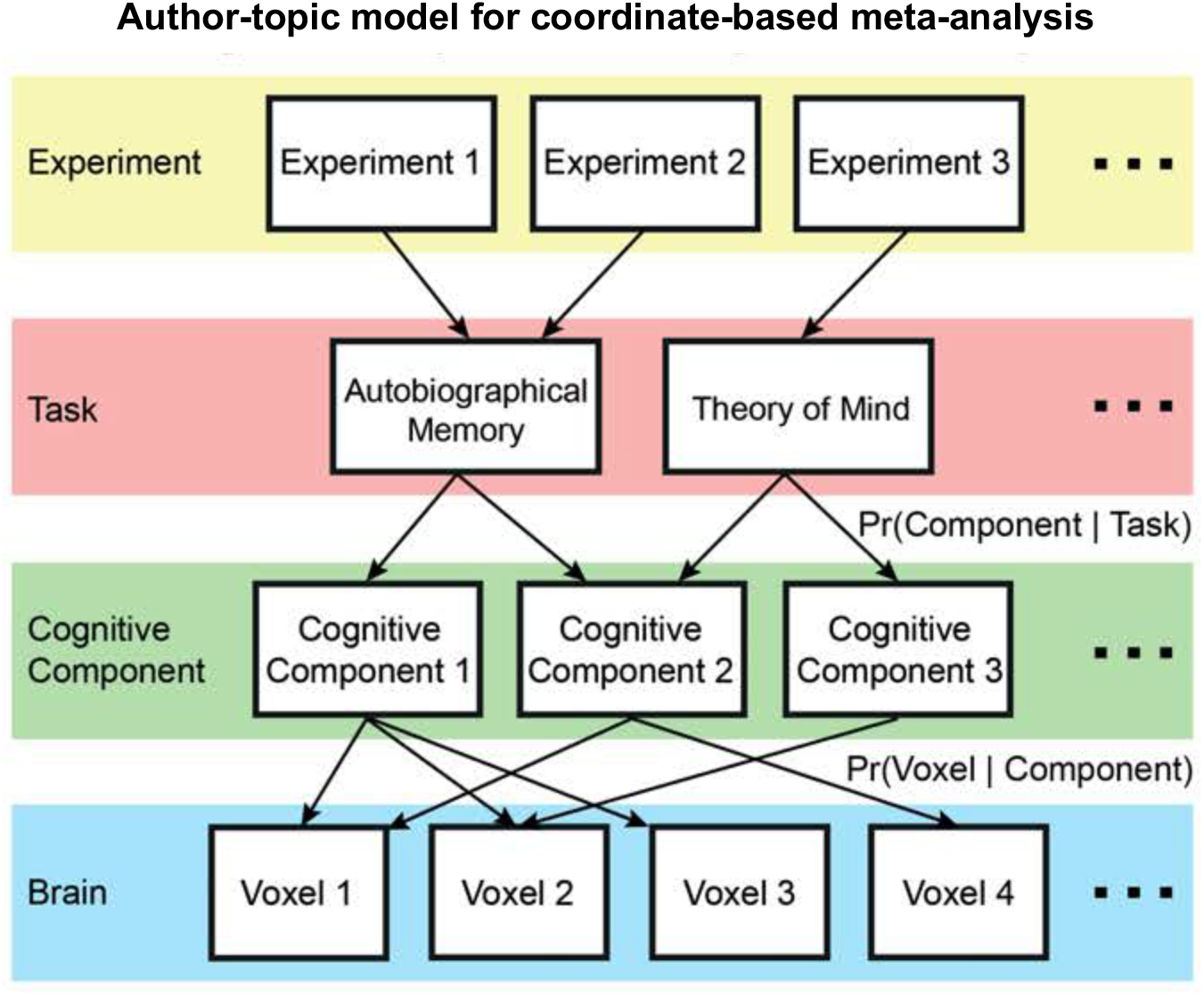
Author-topic model for coordinate-based meta-analysis (Yeo et al. 2015). The underlying premise of the model is that behavioral tasks recruit multiple cognitive components, which are in turn supported by overlapping brain regions. The model parameters are the probability that a task would recruit a cognitive component (Pr(component | task)) and the probability that a component would activate a brain voxel (Pr(voxel | component)). The author-topic model can be directly applied to estimate cognitive components (sub-systems) of self-generated thought.

Our choice of self-generated thought is motivated by previous work suggesting the possibility of fractionating self-generated thought into functional sub-domains (Section 1.1). Similarly, our choice of left IFJ is motivated by previous work suggesting that IFJ might adaptively modify its connectivity patterns across task contexts (Section 1.2). There are of course other functional domains (e.g., executive function) that might be fractionated and other hub regions (e.g., dorsal anterior insula) that might exhibit task-dependent co-activation patterns. Therefore, we have made our code publicly available for researchers to explore the heterogeneity of their preferred functional domain, hub region or mental disorder.

## 2. Methods

### 2.1 Overview

In Section 2.2, we reviewed the author-topic model and how it could be applied to coordinate-based meta-analysis (Yeo et al. 2015). Section 2.3 discussed simulations and comparisons with spatial independent component analysis. Finally, the model was utilized in two different applications. In the first application (Section 2.4), we applied the author-topic model to discover cognitive components subserving self-generated thought. In the second application (Section 2.5), we estimated the co-activation patterns of the left IFJ.

### 2.2 Author-topic model

#### 2.2.1 Intuition behind the model

The author-topic model was originally developed to discover topics from a corpus of text documents (Rosen-Zvi et al., 2010). The model represents each text document as an unordered collection of words written by a group of authors. Each author is associated with a probability distribution over topics, and each topic is associated with a probability distribution over a dictionary of words. Given a corpus of text documents, there are algorithms to estimate the distribution of topics associated with each author and the distribution of words associated with each topic. A topic is in some sense abstract, but is made concrete by its association with certain words and its association with certain authors. For example, if the author-topic model was applied to neuroimaging research articles, the algorithm might yield a topic associated with the author “Stephen Smith” and words like “fMRI”, “resting-state” and “ICA”. One might then interpret the topic posthoc as a “resting-state fMRI” research topic.

In a previous study (Yeo et al., 2015), the author-topic model was applied to neuroimaging meta-analysis (Figure 2) by treating task contrasts in the BrainMap database (Fox and Lancaster, 2002) as text documents, 83 BrainMap task categories (e.g., n-back) as authors, cognitive components as topics, and activation foci as words in the documents. Thus, the model encodes the premise that different behavioral tasks recruit multiple cognitive components, supported by overlapping brain regions.

Suppose a study utilizes one or more task categories, resulting in an experimental contrast yielding a collection of activation foci. Under the author-topic model, each activation focus is assumed to be generated by first randomly selecting a task from the set of tasks utilized in the experiment. Given the task, a component is randomly chosen based on the probability of a task recruiting a component (Pr(component | task)). Given the component, the location of the activation focus is then randomly chosen based on the probability that the component would activate a voxel (Pr(voxel | component)). The entire collections of Pr(component | task) and Pr(voxel | component) are denoted as matrices *θ* and *β*, respectively. For example, the 2nd row and 3rd column of *θ* corresponds to Pr(3rd component | 2nd task) and the 4th row and 28th column of *β* corresponds to Pr(28th voxel | 4th component). Therefore, each row of *θ* and *β* sums to 1. The formal mathematical definition of the model is provided in Supplemental Method S1.

A key property of the author-topic model is that the ordering of words within a document is exchangeable. When applied to meta-analysis, the corresponding assumption is that the ordering of activation foci is arbitrary. Although the ordering of words within a document is obviously important, the ordering of activation foci is not. For example, in the context of text documents, “dog has a bone” has a different meaning from “bone has a dog”. On the other hand, in the context of a fMRI experiment, reporting parietal activation coordinates followed by prefrontal activation coordinates is equivalent to reporting prefrontal activation coordinates followed by parietal activation coordinates. Therefore, the author-topic model is arguably more suitable for meta-analysis than topic discovery from documents.

#### 2.2.2 Estimating the model parameters

Given a collection of experiments with their associated activation coordinates and task categories, as well as the number of cognitive components *K*, the probabilities *θ* and *β* can be estimated using various algorithms (Rosen-Zvi et al. 2010; Yeo et al., 2015; Ngo et al., 2016). Here, we chose to utilize the CVB algorithm because the algorithm was more robust to the choice of hyperparameters in smaller datasets. Although the CVB algorithm for the author-topic model was first introduced in a conference article (Ngo et al., 2016), detailed derivations have not been published. For completeness, detailed derivations of the author-topic CVB algorithm are provided in Supplemental Method S2. Explanations of why the CVB algorithm is theoretically better than the EM algorithm and standard variational Bayes inference are found in Supplemental Method S3. In this work, Bayesian information criterion (BIC) was used to estimate the optimal number of cognitive components (Supplemental Method S4). Further implementation details are found in Supplemental Method S5.

#### 2.2.3 Input to the author-topic model

Each task activation contrast was associated with a set of activation foci. The spatial locations (i.e., coordinates) of the activation foci were reported in or transformed to the MNI152 coordinate system (Lancaster et al., 2007). Using standard meta-analysis procedure (Wager et al., 2009; Yarkoni et al. 2011; Yeo et al. 2015), a 2mm-resolution binary activation image was created for each experimental contrast, in which a voxel was given a value of 1 if it was within a 10mm-radius of any activation focus, and 0 otherwise. Thus, the set activated voxels of each experiment in the author-topic model corresponds to the set of voxels with a value of 1 in the corresponding 2mm-resolution binary activation image. We note that the exact choice of smoothing radius did not significantly affect the results (see Section 3.4).

### 2.3 Simulations

#### 2.3.1 Independent component analysis (ICA)

ICA is a data-driven technique that has been widely applied to fMRI (Calhoun et al. 2001; Beckmann and Smith, 2004). ICA has also been successfully applied to coordinate-based meta-analysis (Smith et al., 2009). However, the author-topic model has a few significant advantages over ICA in the case of coordinate-based meta-analysis. First, activation foci are binary data in the sense that a voxel is either reported to be activated or not in an experiment. However, ICA requires positive and negative values in the input data, which involves demeaning the binary values at each voxel (across experiments). In contrast, the author-topic model makes direct use of the binary activation data. Second, the author-topic model is able to exploit task categorical information (red task layer in Figure 2), which is non-trivial to introduce in ICA.

Most importantly, ICA estimates can be negative, which do not make sense in the case of coordinate-based meta-analysis. For example, a task should not be allowed to be negatively associated with a component, since task activation and de-activation in a coordinate-based meta-analysis are typically handled separately. Similarly, it does not make sense for the activation maps associated with each component to be negative. The situation is of course reversed in image-based meta-analysis (Salimi-Khorshidi et al., 2009), where there might be both activation and de-activation. For image-based meta-analysis, it does make sense to talk about components being negatively recruited by a task and ICA makes more theoretical sense than the author-topic model.

#### 2.3.2 Simulation details

Here, we considered simulations to compare the effectiveness of the author-topic model and ICA. More specifically, we considered a hypothetical situation in which five tasks from a functional domain recruited two cognitive components with different probabilities (Figure 3A). The two components have distinct activation patterns on a 2D “brain” of 256 by 256 pixels. More specifically, each component is associated with activations within two Gaussian distributions centered at two opposite quadrants of the 2D brain (Figure 3B). Given the activation foci of multiple experiments (task contrasts), the goal was to automatically recover the two cognitive components using either the author-topic model or ICA.

**Figure 3.**
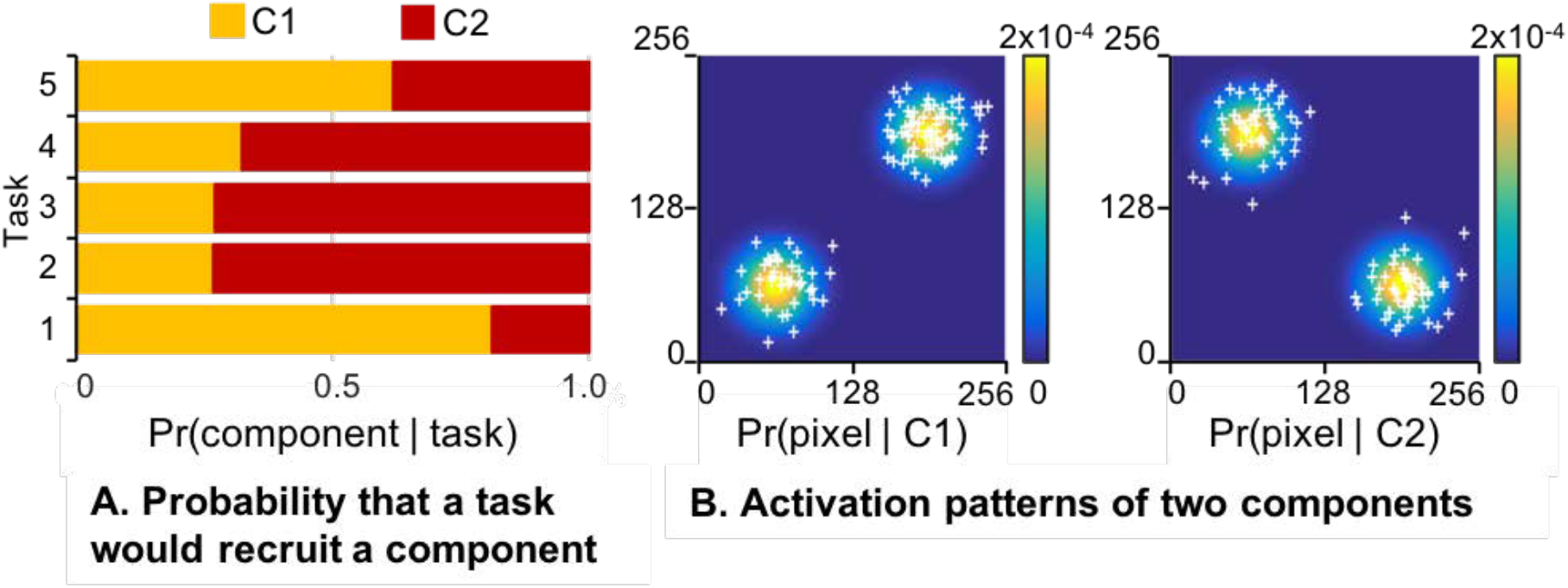
Simulation of heterogeneity in coordinate-based meta-analysis. (A) Bar chart shows five tasks from a functional domain recruiting two cognitive components with different probabilities. (B) Activation patterns of two components on a 2D “brain” of 256 by 256 pixels. Each component is associated with activations (white crosses) within two Gaussian distributions centered at two opposite quadrants of the 2D “brain”. For each simulation run, the probability of a task recruiting a component and the covariances of each component’s 2D Gaussian distributions were randomly generated. The author-topic model and ICA were then applied to recover the two components. We note that ICA mixture weights can be negative, which does not make sense in the context of coordinate-based meta-analysis. As such, we discarded simulation runs if any of the ICA estimates yielded negative weights.

A single simulation run comprised 150 experiments (task contrasts), which is comparable to a typical meta-analysis (c.f. self-generated thought in Section 2.4). Each experiment (task contrast) was randomly assigned to one of the five tasks, with the contrast distributions skewed towards two of the five tasks to simulate the fact that some tasks are more popular than others in the literature. Furthermore, each task contrast is randomly chosen to have between 1 to 10 activation foci. For each activation focus, a component was randomly sampled based on the probability of components given the task assigned to the experiment. For the given component, one of the 2-D Gaussian distributions of each component was randomly chosen with equal probabilities (Figure 3B). The spatial location of the activation focus was then randomly sampled from the Gaussian distribution. The activation focus was smoothed with a binary smoothing kernel, such that all pixels within 10 voxels from an activation focus were given a value of 1, and 0 otherwise.

For a given simulation run, the latent components were estimated using either ICA or the author-topic model. We considered three possible ICA setups. The first two setups (ICA1 and ICA2) utilized CanICA (Varoquaux et al., 2010), an ICA decomposition implementation provided with Nilearn (Abraham et al. 2014). CanICA extracts representative patterns of multi-subject fMRI data by performing ICA on a data subspace common to the group (Varoquaux et al., 2010). In the two setups ICA1 and ICA2, each task was treated as a subject. In ICA1, the activation maps of all experiments assigned to the same task were summed together, i.e., each task was treated as a single subject with a single time point. In ICA2, each task was treated as a single subject, but the experiments assigned to the given task were treated as separate time points of the subject. The third setup (ICA3) utilized the MELODIC implementation of ICA from the FSL package (Beckmann & Smith 2004; Smith et al. 2004).

To evaluate the estimation quality, Pearson’s correlation coefficient was computed between the groundtruth probability distribution of a component activating a vertex (Pr(vertex | component)) against the estimates from the author-topic model or ICA. Pearson’s correlation coefficient was also computed between the groundtruth distribution of components given a task (Pr(component | task)) and estimates from the author-topic model or ICA.

The simulation was repeated multiple times. For a given simulation run, the covariances of each component’s 2D Gaussian distributions were randomly generated (Figure 3B). The probability of a task recruiting a component was also randomly generated (Figure 3A). As explained previously (Section 2.3.1), ICA’s mixture weights can be negative, which implies negative associations between tasks and components. This does not make sense in the case of coordinate-based meta-analysis, so we discarded simulation runs if any of the ICA estimates yielded negative weights. Overall, we ran roughly 300 simulation runs in order to yield exactly 100 simulation runs, in which ICA estimates were valid.

### 2.4 Self-generated thought

#### 2.4.1 Activation foci of experiments involving self-generated thought

To explore cognitive components subserving self-generated thought, we considered 1812 activation foci from 179 experimental contrasts across 167 imaging studies, each employing one of seven task categories subjected to prior meta-analysis with GingerALE (Fox and Lancaster, 2002; Laird et al., 2009, 2011; Fox et al., 2014; http://brainmap.org/ale). Of the 167 studies, 48 studies employed “Autobiographical Memory” (N = 19), “Navigation” (N = 13) or “Task Deactivation” (N = 16) tasks. The 48 studies were employed in a previous meta-analysis examining the default network (Spreng et al., 2009). There were 79 studies involving “Story-based Theory of Mind” (N = 18), “Nonstory-based Theory of Mind” (N = 42) and “Narrative Comprehension” (N = 19) tasks. The 79 studies were utilized in a previous meta-analysis examining social cognition and story comprehension (Mar, 2011). Finally, there were 40 studies involving the “Moral Cognition” task that was again utilized in a previous meta-analysis (Sevinc and Spreng, 2014). The list of all experiments included in the dataset are provided in Supplemental Method S7. The criteria for selecting the experiments can be found in the original meta-analyses (Spreng et al., 2009; Mar, 2011; Sevinc and Spreng, 2014). All foci coordinates were in or transformed to the MNI152 coordinate system (Lancaster et al., 2007).

#### 2.4.2 Discovering cognitive components of self-generated thought

The application of the author-topic model to discover cognitive components subserving self-generated thought (Figure 2) is conceptually similar to the original application to the BrainMap (Yeo et al., 2015). The key difference is that the current application is restricted to seven related tasks in order to discover heterogeneity within a single functional domain, while the original application sought to find common and distinct cognitive components across domains.

The model parameters are the probability of a task recruiting a component (Pr(component | task)) and the probability of a component activating a brain voxel (Pr(voxel | component)). The parameters were estimated from the 1812 activation foci from the previous section using the CVB algorithm (Supplemental Methods S2 and S5). BIC was used to estimate the optimal number of cognitive components (Supplemental Method S4).

#### 2.4.3 Interpreting cognitive components of self-generated thought

The matrix Pr(voxel | component), *β*, can be interpreted as *K* brain images in MNI152 coordinate system (Lancaster et al., 2007), where *K* is the number of cognitive components. Volumetric slices highlighting specific subcortical structures were displayed using FreeSurfer (Fischl, 2012). The cerebral cortex was visualized by transforming the volumetric images from MNI152 space to fs_LR surface space using Connetome Workbench (Van Essen et al., 2013) via the FreeSurfer surface space (Buckner et al., 2011; Fischl et al., 2012). For visualization purpose, isolated surface clusters with less than 20 vertices were removed, Pr(component | task) was thresholded at 1/K, and Pr(voxel | component) was thresholded at 1e-5, consistent with previous work (Yeo et al., 2015). Unthresholded maps of the components are available on NeuroVault (Gorgolewski et al., 2015) at https://neurovault.org/collections/4684/.

#### 2.4.4 Goodness of fit

For each task, we computed the weighted average of the cognitive components (Pr(voxel | component)), where the weights corresponded to the probabilities of the task recruiting the components (Pr(component | task)). This weighted average spatial map could be interpreted as the model estimate of the “ideal” (reconstructed) activation map for a particular task. The model fit was good if a task’s reconstructed activation map was similar to the empirical activation map of the task (obtained by averaging the activation maps of all experiments employing the task). Therefore, we computed Pearson’s correlation coefficient between all pairs of reconstructed and empirical activation maps, yielding a 7 x 7 correlation matrix (since there were 7 tasks).

#### 2.4.5 Correspondence between cognitive components and resting-state networks

Motivated by similarities between task and resting-state networks (Smith et al., 2009; Laird et al., 2011; Yeo et al., 2015), we compared the cognitive components of self-generated thought with a previously published set of 17 resting-state networks (Yeo et al., 2011). For each resting-state network and each cognitive component, the probability of the cognitive component activating a voxel (Pr(voxel | component)) was averaged across all voxels within the network, resulting in an average probability of a component activating the given network.

### 2.5 Left inferior frontal junction (IFJ)

#### 2.5.1 Activation foci of experiments activating the left IFJ

To explore task-dependent co-activation patterns expressed by the left IFJ, we considered activation foci from experiments reporting activation within a left IFJ seed region (Figure S1) delineated by a previous study (Muhle-Karbe et al. 2015). Muhle-Karbe and colleagues performed a co-activation-based parcellation of a left lateral prefrontal region into six parcels, including an IFJ region (Muhle-Karbe et al., 2015). The parcellation procedure assumed that voxels within a parcel exhibited a single co-activation pattern. Thus, the advantage of using this particular IFJ seed region (instead of an IFJ region from a different study) is that this region is thought to exhibit a single co-activation pattern (according to MACM).

This seed region is publicly available on ANIMA (Reid et al. 2016; http://anima.fz-juelich.de/studies/MuhleKarbe_2015_IFJ). We selected experiments from the BrainMap database with at least one activation focus falling within the IFJ seed region. We further restricted our analyses to experimental contrasts involving normal subjects. Overall, there were 323 experiment contrasts from 238 studies with a total of 5201 activation foci. The list of all experiments included in the dataset are provided in Supplemental Method S8.

#### 2.5.2 Discovering co-activation patterns of the IFJ

To apply the author-topic model to discover co-activation patterns, we consider each of the 323 experimental contrasts to employ its own unique task category (Figure 4). In the parlance of the author-topic model, we assumed each document (experiment) has its own unique author (task). The premise of the model is that the IFJ expresses one or more overlapping co-activation patterns depending on task contexts. A single experiment activating the IFJ might recruit one or more co-activation patterns. The model parameters are the probability that an experiment would recruit a co-activation pattern (Pr(co-activation pattern | experiment)), and the probability that a voxel would be involved in a co-activation pattern (Pr(voxel | co-activation pattern)). The parameters were estimated from the 5201 activation foci from the previous section using the CVB algorithm (Supplemental Method S2 and S5). BIC was used to estimate the optimal number of co-activation patterns (Supplemental Method S4).

**Figure 4.**
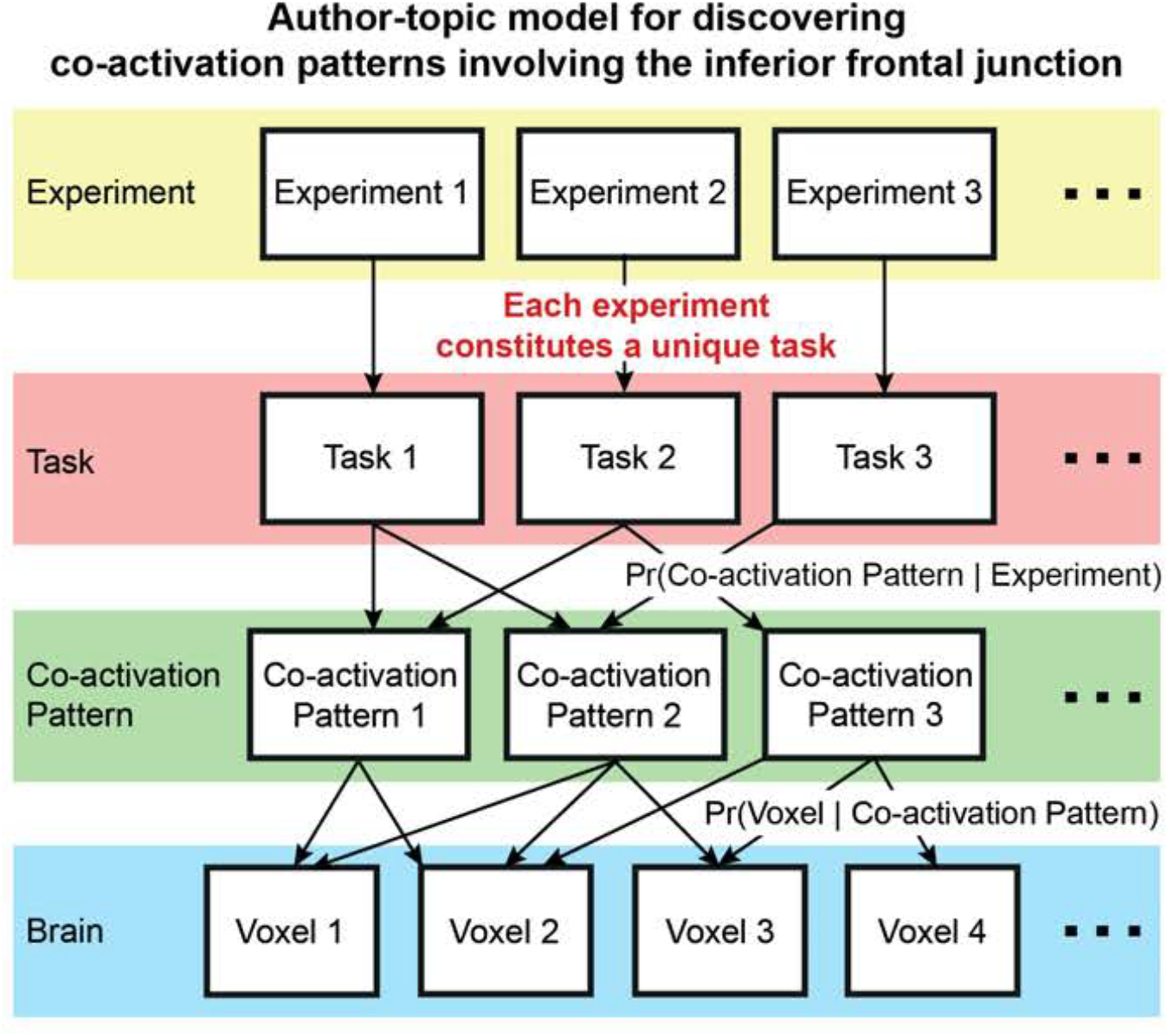
Author-topic model for discovering co-activation patterns of the inferior frontal junction (IFJ). In contrast to Figure 2, this instantiation of the model assumes that each experiment constitutes a unique task. The premise of the model is that the IFJ expresses multiple overlapping task-dependent co-activation patterns. The model parameters are the probability of an experiment recruiting a co-activation pattern (Pr(co-activation pattern | experiment)), and the probability of a voxel being associated with a co-activation pattern (Pr(voxel | co-activation pattern)).

#### 2.5.3 Interpreting co-activation patterns of the IFJ

Similar to the previous application on self-generated thought, the matrix Pr(voxel | co-activation pattern), *β*, was visualized as *K* brain images in both fsLR surface space and MNI152 volumetric space. Like before, isolated surface clusters with less than 20 vertices were removed for the purpose of visualization. Unthresholded spatial maps of the co-activation patterns are available on NeuroVault (Gorgolewski et al., 2015) at https://neurovault.org/collections/4718/.

Because each of the 323 experiments was treated as employing a unique task category, Pr(co-activation pattern | experiment), *θ*, is a matrix of size *K* × 323. *θ* was further mapped onto BrainMap task categories to assist in the interpretation. More specifically, since the experiments were extracted from the BrainMap database, each experiment was tagged with one or more BrainMap task categories (Table S1). The Pr(co-activation pattern | experiment) was averaged across experiments employing the same task category to estimate the probability that a task category would recruit a co-activation pattern (Pr(co-activation pattern c | task t)). Further details of this procedure are found in Supplemental Method S6.

We note that directly using the BrainMap task categories to interpret the co-activation patterns is tricky. This is because a BrainMap task category might only have a very small percentage of experiments activating the IFJ, so these experiments might not be representative of the task category. For example, of the 230 experiments in the BrainMap database labeled as the “Encoding” task category, only 13 experiments reported activations in the left IFJ. Thus, the 13 experiments were not simply encoding tasks, but encoding tasks that happened to activate the IFJ. This is the reason why the BrainMap task categories were not directly utilized in the author-topic model for the IFJ analysis and that each experiment was treated as employing a unique task category (c.f. self-generated thought in Section 2.4).

To ensure an appropriate interpretation, we inspected the original publications associated with the top three experiments with the highest Pr(co-activation pattern | experiment) for each of the top three tasks associated with each co-activation pattern, i.e., nine publications for each co-activation pattern. The literature analysis allowed us to determine if there were common neural processes underlying the subset of experiments within each task category that strongly activated the IFJ.

#### 2.5.4 Goodness of fit

For each co-activation pattern, activation maps of the top three experiments with the highest probability of recruiting a co-activation pattern (i.e., Pr(co-activation pattern | experiment)) for each of the top three tasks associated with the co-activation pattern (i.e., nine activation maps in total) were averaged, resulting in an empirical activation map associated with each co-activation pattern. The model fit was good if the empirical activation map was similar to the estimated co-activated pattern. Therefore, we computed Pearson’s correlation coefficient between all pairs of empirical activation maps and co-activation maps, yielding a *K* × *K* correlation matrix, where K is the number of co-activation patterns estimated by BIC.

### 2.6 Data and code availability

Activation foci from the meta-analysis of self-generated thought and the source code of the author-topic model, including the visualization and analysis tools, are publicly available at https://github.com/ThomasYeoLab/CBIG/tree/master/stable_projects/meta-analysis/Ngo2019_AuthorTopic. The activation foci from the meta-analysis of IFJ can be obtained via a collaborative-use license agreement with BrainMap (http://www.brainmap.org/collaborations.html).

## 3 Results

### 3.1 Overview

In Section 3.2, we show simulation results suggesting that the author-topic model compares favorably with ICA in the goal of discovering latent patterns in coordinate-based meta-analysis. We then explored the cognitive components of self-generated thought (Section 3.3) and the co-activation patterns of the IFJ (Section 3.4). Finally, Section 3.5 discusses a few control analyses.

### 3.2 Simulations

Figure 5 shows the results of one representative simulation (see Section 2.3 for details). Figure 5A shows the groundtruth 2D “brain” maps for this representative simulation run. The two leftmost columns show simulated activation foci as white crosses overlaid on top of the 2D Gaussian distributions used to generate the foci. The rightmost bar chart shows the probability of each of the 5 tasks recruiting a component.

**Figure 5.**
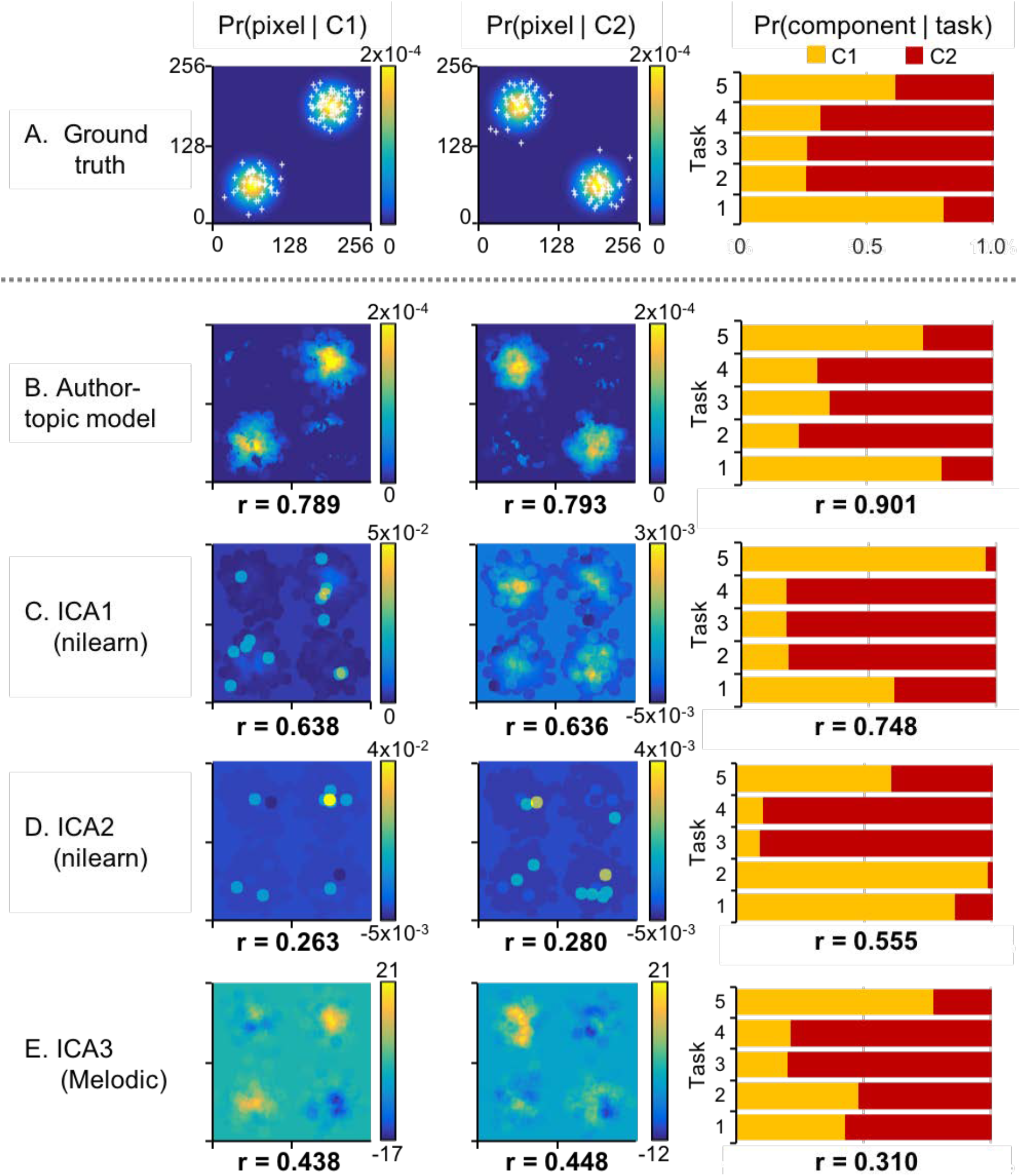
Simulation comparing the author-topic model and ICA. (A) Single representative simulation run. Two leftmost columns show activation foci (white crosses) on top of Gaussian distributions used to generate the foci. Rightmost bar chart shows the probability of each of the 5 tasks recruiting a component. (B) Author-topic model estimates. (C) ICA estimates. Number below each panel is the correlation between model estimates and groundtruth averaged across 100 simulation runs. Observe that ICA can yield negative weights, which do not make sense in the context of a coordinate-based meta-analysis (see discussion in Section 2.3.1). We note that about 300 simulation runs were run in order to generate 100 simulation runs in which ICA estimates of mixture weights were non-negative.

The rightmost column of Figure 5B shows the author-topic model estimates of the probability of each of the 5 tasks recruiting a component. The rightmost column of Figures 5C to 5E shows the ICA mixture weights, normalized so they sum to one1. The mixture weights represent the association between the tasks and the components. The numbers at the bottom of each panel are the correlations between the estimates and groundtruth averaged across 100 simulation runs. In general, the author-topic model yielded better estimates of the associations between tasks and components.

Figure 5B shows the author-topic model estimates, while Figures 5C to 5E show the ICA estimates. The two leftmost columns show the spatial maps of the two components estimated by the author-topic model or ICA. The numbers at the bottom of each panel are the correlations between the estimated and groundtruth “brain” maps averaged across 100 simulation runs. In general, the author-topic model yielded better estimates of the groundtruth “brain” maps. It is also worth noting that the ICA spatial maps showed negative values, even though the simulation runs had been constrained to those where ICA mixture weights were positive2. As previously explained (Section 2.3.1), negative values are not meaningful in the context of coordinate-based meta-analysis.

### 3.3 Self-generated thought

#### 3.3.1 ALE meta-analysis of self-generated thought

Figure 6 shows the activation likelihood estimate (ALE) of experiments involving self-generated thought. Statistical significance was established with 1000 permutations. The map was thresholded at a voxel-wise uncorrected threshold of p < 0.001 and cluster-level family-wise error rate threshold of p < 0.01. Consistent with previous studies, ALE reveals a constellation of regions typically referred to as the default network (Raichle et al., 2001; Buckner et al. 2008; Spreng et al. 2009). However, as previously discussed, ALE cannot reveal functional sub-domains within self-generated thought without prior assumptions about the sub-domains. Therefore, in the next section, we explored the use of the author-topic model.

**Figure 6.**
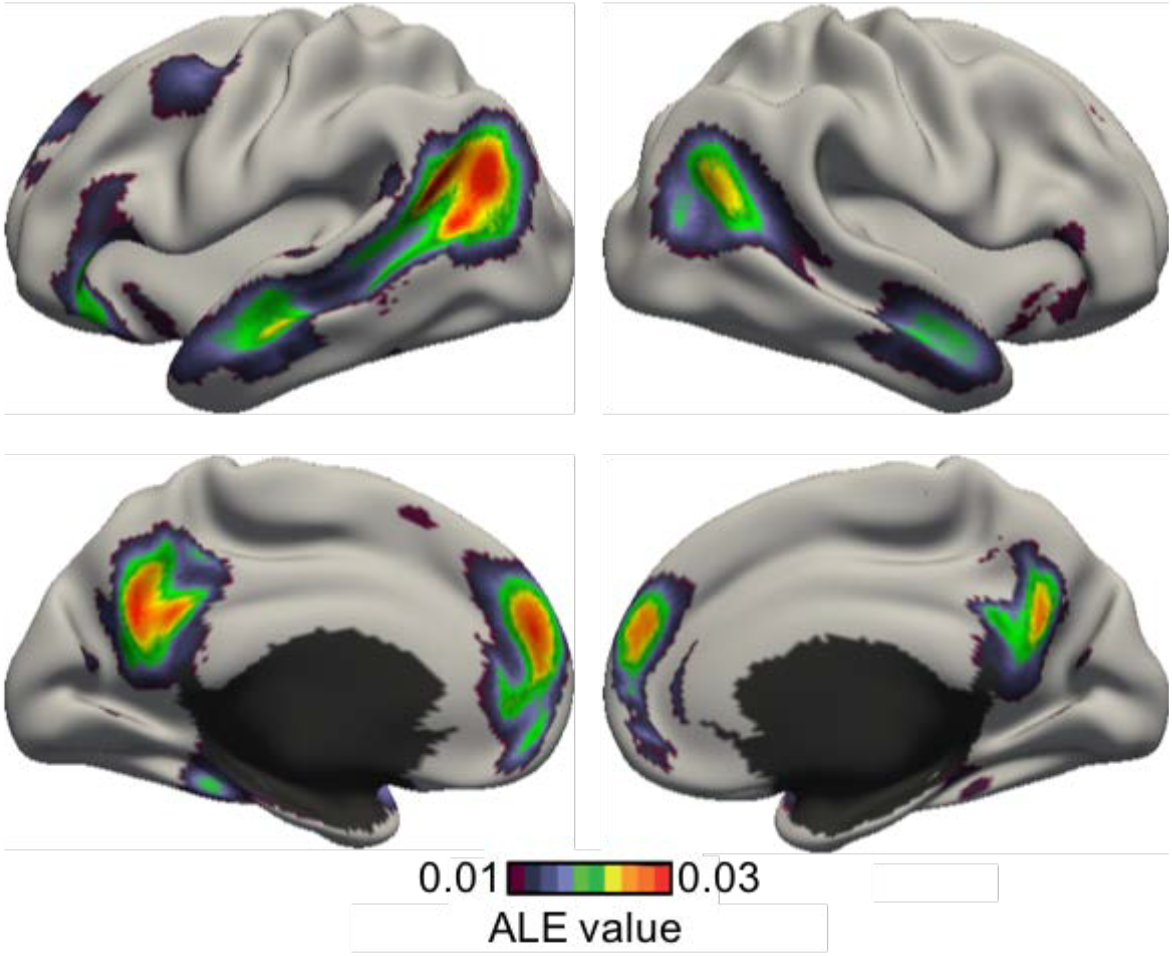
Activation likelihood estimate (ALE) of experiments involving self-generated thought. Consistent with previous studies, ALE reveals a set of regions corresponding to the default network. However, ALE cannot provide insights into functional sub-domains without prior assumptions about the sub-domains.

#### 3.3.2 Cognitive components of self-generated thought

Figure 7 shows the cognitive components of self-generated thought estimated by the author-topic model. Figure 7A shows the BIC score as a function of the number of estimated cognitive components. A higher BIC score indicates a better model. Because the 2-component estimate achieved the highest BIC score, subsequent results will focus on the 2-component estimate.

**Figure 7.**
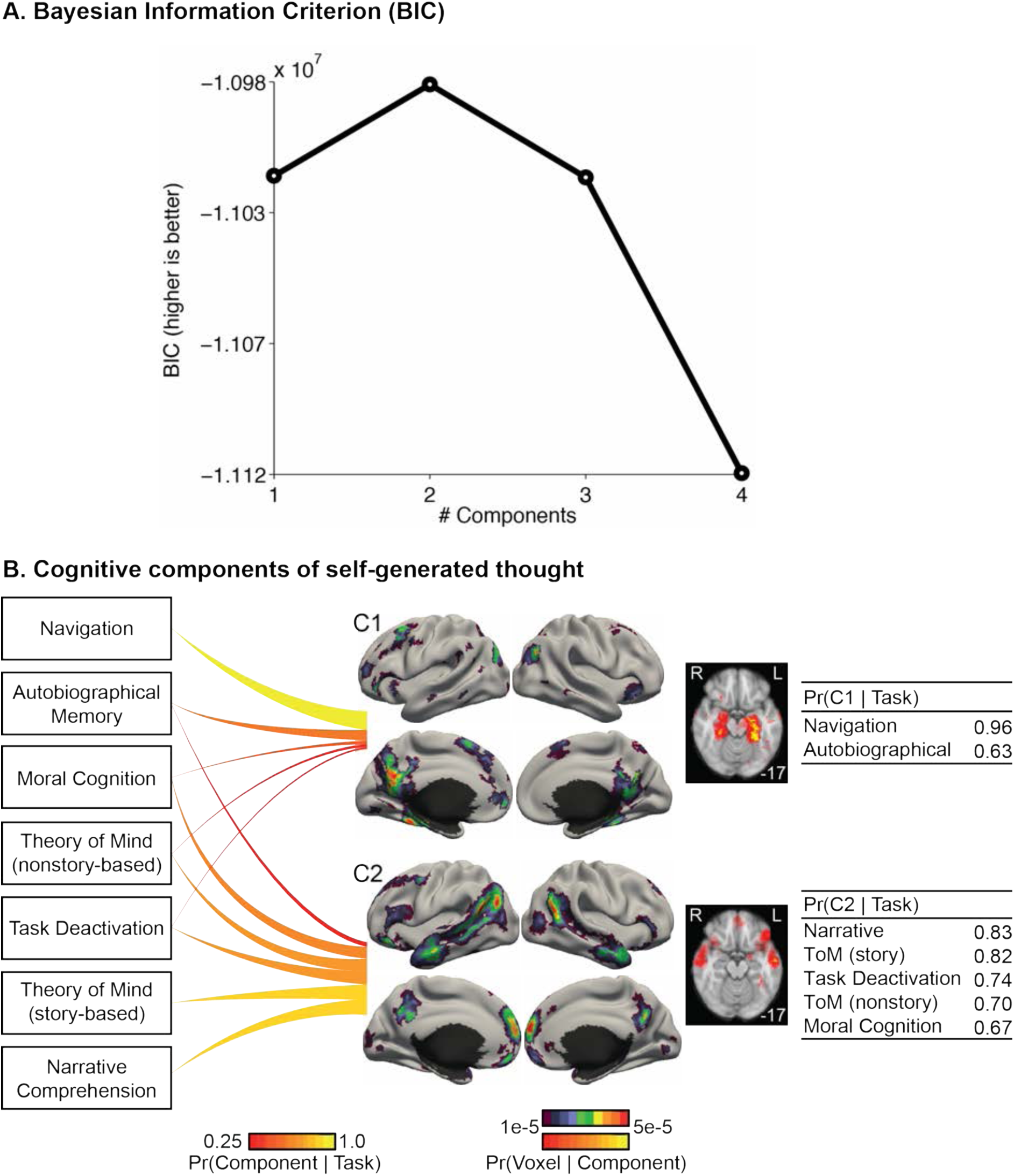
Cognitive components of self-generated thoughts. (A) Bayesian Information Criterion (BIC) plotted as a function of the number of estimated cognitive components. A higher BIC indicates a better model. BIC peaks at 2 components. (B) 2-component model estimates. Each line connects 1 task with 1 component. The thickness and brightness of the lines are proportional to the magnitude of Pr(component | task). For each component, the four leftmost figures show the surface-based visualization for the probability of components activating different brain voxels (i.e., Pr(voxel | component)), whereas the rightmost figure show a volumetric slice highlighting subcortical structures being activated differently across components. The top color bar is utilized for the surface-based visualization, whereas the bottom color bar is utilized for the volumetric slices. The tables on the right show the top tasks most likely to recruit the two components. The numbers in the right column correspond to Pr (component | task). Navigation and Autobiographical Memory preferentially recruited component C1, whereas Narrative Comprehension, Theory of Mind (ToM), Task Deactivation and Moral Cognition preferentially recruited component C2.

The 2-component estimate is shown in Figure 7B. The seven tasks recruited the two cognitive components to different degrees. The top tasks recruiting component C1 were “Navigation” and “Autobiographical Memory”. In contrast, the top tasks recruiting component C2 were “Narrative Comprehension”, “Theory of Mind (story-based)”, “Task deactivation”, “Theory of Mind (nonstory-based)”, and “Moral Cognition”.

Compared with Figure 6, the two cognitive components appeared to decompose the activation pattern revealed by ALE. The two cognitive components appeared to activate different portions of the default network (Figure 7B). Focusing our attention to the medial cortex, both components had high probability of activating the medial parietal cortex. However, while component C2’s activation was largely limited to the precuneus, component C1’s activation also included the posterior cingulate and retrosplenial cortices in addition to the precuneus. Both components also had high probability of activating the medial prefrontal cortex (MPFC). However, component C1’s activations were restricted to the middle portion of the MPFC, while component C2’s activations were restricted to the dorsal and ventral portions of the MPFC. Finally, component C1, but not component C2, had high probability of activating the hippocampal complex.

Switching our attention to the lateral cortex, component C1 had high probability of activating the posterior inferior parietal cortex, while component C2 had high probability of activating the entire stretch of cortex from the temporo-parietal junction to the temporal pole. Component C2 was significantly more likely than component C1 to activate the inferior frontal gyrus.

#### 3.3.3 Goodness of fit

Figure 8 shows the correlation matrix between the empirical activation maps of seven tasks involving self-generated thought (rows) and seven task activation maps reconstructed from the author-topic model parameter estimates (columns). The diagonal entries of the correlation matrix were significantly higher than the off-diagonal entries: average diagonal entry was 0.69, while the average off-diagonal entry was 0.50. Overall, this suggests a good model fit. However, the diagonal entries were not always the highest and there was a clear block-diagonal structure. Not surprisingly, the top left block corresponded to the top two tasks recruiting component C1 (Figure 7), while the bottom right block corresponded to the top five tasks recruiting component C2 (Figure 7).

**Figure 8.**
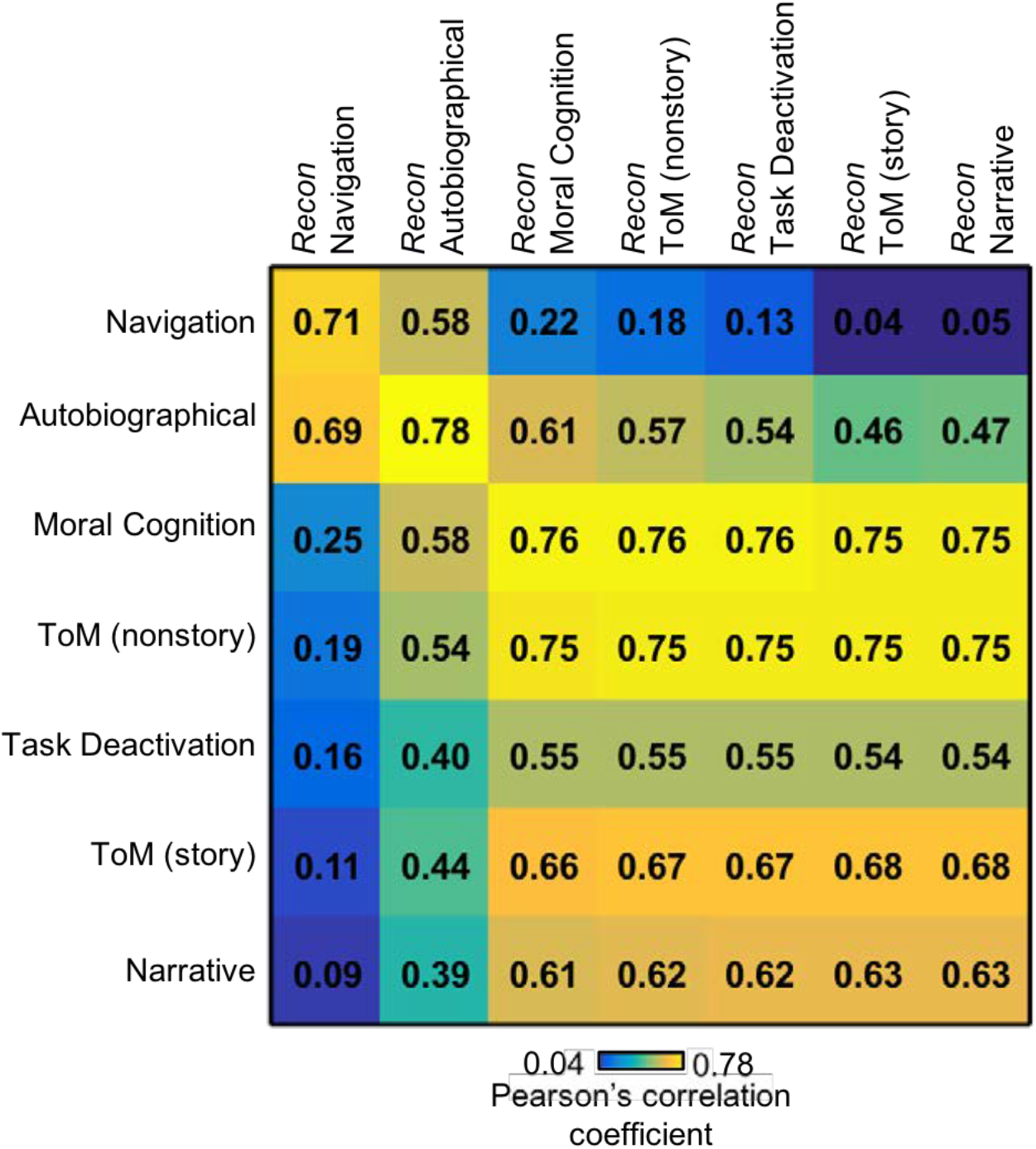
Goodness of fit of the author-topic model for self-generated thought. The matrix represents the correlations between the empirical activation maps (rows) and reconstructed activation maps (columns) of seven tasks. The tasks follow the same ordering as in Figure 7. The diagonal values (average r = 0.69) were larger than off-diagonal values (average r = 0.50), suggesting a good model fit.

#### 3.3.4 Correspondence with resting-state networks

The average probability of each self-generated thought cognitive component activating each resting-state network (Yeo et al., 2011) is shown in Figure S2. Four resting-state networks with the highest probability of being activated by either component are shown in Figure 9. Three of these resting-state networks were previously considered to be fractionation of the default network (Yeo et al., 2014).

**Figure 9.**
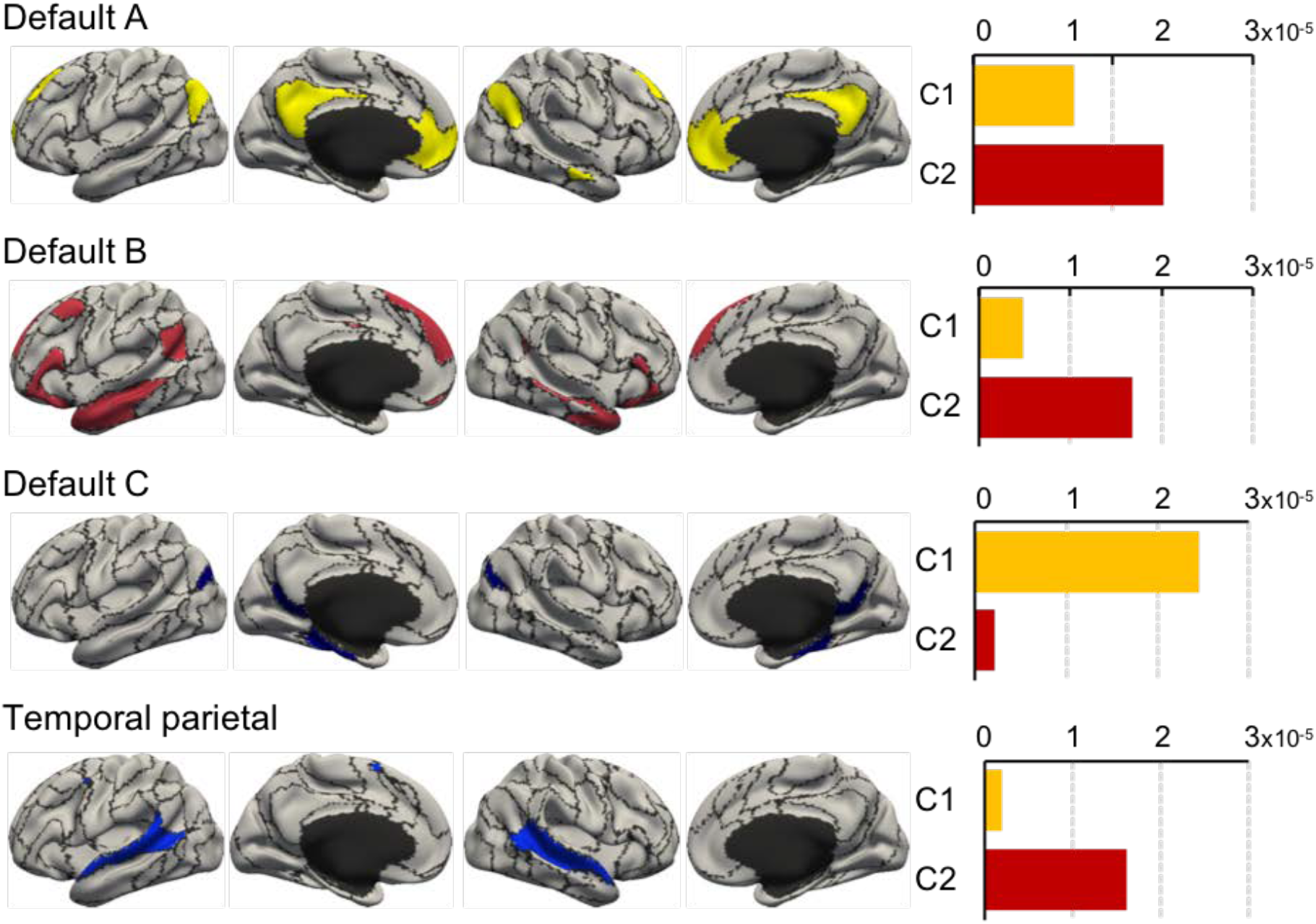
Average probability of self-generated thought cognitive components activating voxels within 4 resting-state networks (Yeo et al. 2011). The naming of the four resting-state networks followed the convention of previous literature (Kong et al., 2018; Li et al., 2018). Default C resting-state network was primarily activated by component C1, while the temporal parietal resting-state network was primarily activated by component C2. On the other hand, Default A and B resting-state networks were preferentially activated by component C2.

The Default C resting-state network was most strongly activated by component C1, while the temporal parietal resting-state network was most strongly activated by component C2. On the other hand, Default A and B resting-state networks were preferentially activated by components C2.

### 3.4 Left inferior frontal junction (IFJ)

#### 3.4.1 ALE meta-analysis of the left IFJ’s co-activation pattern

Figure 10 shows the co-activation pattern of the left IFJ estimated by the application of ALE to meta-analytic co-activation modeling (Muhle-Karbe et al., 2015). Statistical significance was established with 1000 permutations. The map was thresholded at a voxel-wise uncorrected threshold of p < 0.001 and cluster-level family-wise error rate threshold of p < 0.01. The co-activation pattern was mostly bilateral and involved dorsolateral prefrontal cortex, anterior insula, superior parietal lobules, posterior medial frontal cortex and the fusiform gyri. As previously discussed, ALE delineates regions consistently activated across studies, but cannot reveal potential task-dependent co-activation patterns. Therefore, in the next section, we explored the use of the author-topic model.

**Figure 10.**
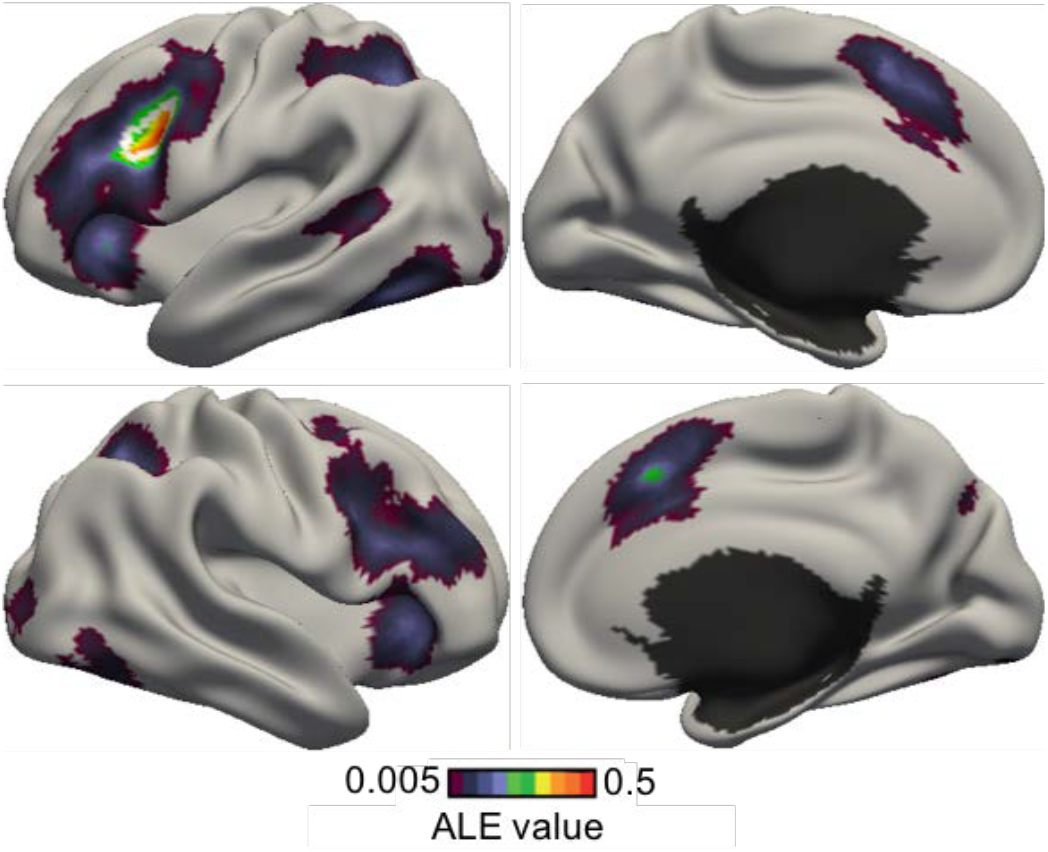
Co-activation pattern of the left inferior frontal junction (IFJ) estimated by the application of ALE to perform meta-analytic co-activation mapping. The IFJ seed region is delineated by a white boundary.

#### 3.4.2 Task-dependent co-activation patterns of the left IFJ

Figure 11 shows the co-activation patterns of the left IFJ estimated by the author-topic model. Figure 11A shows the BIC score as a function of the number of estimated co-activation patterns. There were two peaks corresponding to the 3-pattern and 5-pattern estimates. Figure S3 shows the 5-pattern estimate. Although the 5-pattern estimate had a higher BIC score than the 3-pattern estimate, the co-activation patterns appeared to fractionate the IFJ into smaller territories. While this fractionation was intriguing, our goal was to examine if the IFJ exhibited task-dependent co-activation patterns and not whether it can be further fractionated. Thus, the 5-pattern estimate represented a degenerate solution from this perspective.

**Figure 11.**
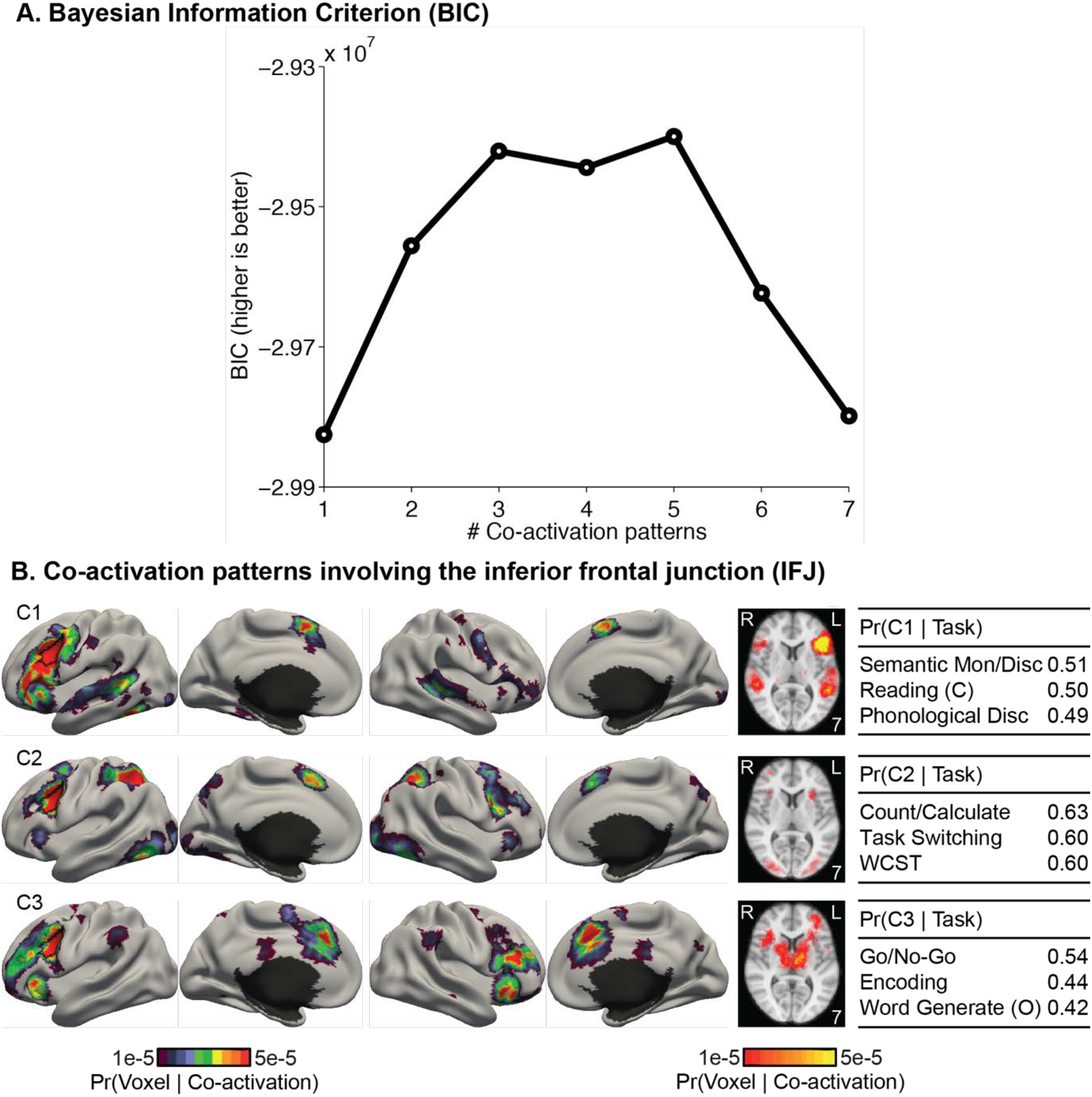
Co-activation patterns involving the inferior frontal junction (IFJ). (A) Bayesian Information Criterion (BIC) plotted as a function of the number of estimated co-activation patterns. BIC peaks at 3 co-activation patterns. (B) 3-coactivation-pattern model estimates for the IFJ. Format follows Figure 7. “(C)” and “(O)” indicate “covert” and “overt” respectively. “Mon”, and “Disc” are short for “monitor” and “discrimination” respectively. “Count/ Calculate” is short for “Counting/Calculation”. “WCST” is short for “Wisconsin Card Sorting Test”. The left IFJ is delineated by the black boundary in the left hemisphere.

Figure 11B shows the co-activation patterns from the 3-pattern estimate. Unlike the 5-pattern estimate, the 3 co-activation patterns appeared to overlap completely within the IFJ. Therefore, subsequent results will focus on the 3-pattern estimate. Overall the 3 co-activation patterns appeared to decompose the consensus co-activation pattern revealed by ALE (Figure 10).

Co-activation pattern C1 was left lateralized and might be recruited in tasks involving language processing. Co-activation pattern C2 involved bilateral superior parietal and posterior medial frontal cortices, and might be recruited in tasks involving attentional control. Co-activation pattern C3 involved bilateral frontal cortex, anterior insula and posterior medial frontal cortex, and might be recruited in tasks involving inhibition or response conflicts.

We now discuss in detail spatial differences among the co-activation patterns. Co-activation pattern C3 strongly engaged bilateral anterior insula, while co-activation pattern C1 only engaged left anterior insula. The activation of the anterior insula was much weaker in co-activation pattern C2.

In the frontal cortex, co-activation pattern C1 had high probability of activating the left inferior frontal gyrus, while co-activation pattern C3 had high probability of activating bilateral dorsal lateral prefrontal cortex. Although all three co-activation patterns also had high probability of activating the posterior medial frontal cortex (PMFC), the activation shifted anteriorly from co-activation patterns C1 to C2 to C3.

In the parietal cortex, co-activation pattern C2 included the superior parietal lobule and the intraparietal sulcus in both hemispheres. C1 and C3 did not activate the superior parietal cortex. Finally, co-activation pattern C1 engaged bilateral superior temporal cortex, which might overlap with early auditory regions. Both co-activation patterns C1 and C2 also had high probability of activating ventral visual regions, especially in the fusiform gyrus.

The top three tasks recruiting each co-activation pattern is shown in Figure 11B. For completeness, the top five tasks recruiting each co-activation pattern are shown in Table S2. The top tasks with the highest probability of recruiting co-activation pattern C1 were “Semantic Monitoring/Discrimination”, “Covert Reading”, and “Phonological Discrimination”. The top tasks recruiting co-activation pattern C2 were “Counting/Calculation”, “Task Switching”, and “Wisconsin Card Sorting Test”. The top tasks recruiting co-activation pattern C3 were “Go/ No-Go”, “Encoding”, and “Overt Word Generation”.

At first glance, the top three tasks for co-activation pattern C3 (“Go/ No-Go”, “Encoding”, and “Overt Word Generation”) might not seem to be similar tasks. The reason for this incongruence was previously explained in Section 2.5.3 and was due to the fact that the experiments activating IFJ might not be representative of their task categories. Indeed, of the 123 BrainMap experiments labeled as the “Overt Word Generation” task, only 6 experiments reported activation in the IFJ. Thus, the 6 experiments were not simply “Overt Word Generation” task, but “Overt Word Generation” experiments that happened to activate the IFJ. This motivated further examination of the original publications associated with the top experiments activating IFJ in order to interpret the co-activation patterns (see Section 4.2 for discussion).

Table S3A-S3C list the top three experiments with the highest Pr(co-activation pattern | experiment) for each of the top three tasks associated with each co-activation pattern. For example, Table S3-A lists the top three experiments employing “Semantic Monitoring/Discrimination”, “Covert Reading” or “Phonological Discrimination” with the highest Pr(co-activation pattern C1 | experiment).

To further ensure that the 3 co-activation patterns were not fractionating IFJ (like the 5-pattern estimate), Figure S4 illustrates the activation foci of the top three experiments with the highest Pr(co-activation pattern | experiment) for each of the top three tasks associated with each co-activation pattern falling inside the IFJ. Table 1 shows the mean and standard deviation of the coordinates of these activation foci (within IFJ) for each co-activation pattern. The mean locations of the IFJ activations across co-activation patterns did not differ by more than 2.5mm along any dimension, suggesting that the co-activation patterns were probably not simply sub-dividing the IFJ.

**Table 1.** Spatial locations of activation foci within the IFJ. Each row of the table shows the mean (standard deviation) of the coordinates of the activation foci (within IFJ) reported by the top 3 experiments with the highest Pr(co-activation pattern | experiment) for each of the top three tasks associated with each co-activation pattern falling inside the IFJ. See Figure S4 for volumetric slices illustrating the locations of the activation foci within the IFJ. Across the 3 co-activation patterns, the mean coordinates of the top experiments do not differ by more than 2.5mm in any dimension, suggesting that the co-activation patterns were not fractionating the IFJ

|  | x/mm | y/mm | z/mm |
| --- | --- | --- | --- |
| **Co-activation pattern C1** | -40.33 (1.80) | 3.89 (3.55) | 30.67 (4.21) |
| **Co-activation pattern C2** | -39.33 (3.16) | 5.33 (4.92) | 31.78 (5.45) |
| **Co-activation pattern C3** | -39.40 (4.60) | 6.40 (5.68) | 29.70 (4.64) |

#### 3.4.3 Goodness of fit

Figure 12 shows the correlation matrix between IFJ’s co-activation patterns (columns) and the average activation maps of the top three tasks associated with each co-activation pattern (rows). The diagonal entries of the correlation matrix were significantly higher than the off-diagonal entries: average diagonal entry was 0.75, while the average off-diagonal entry was 0.31. Overall, this suggests a good model fit.

**Figure 12.**
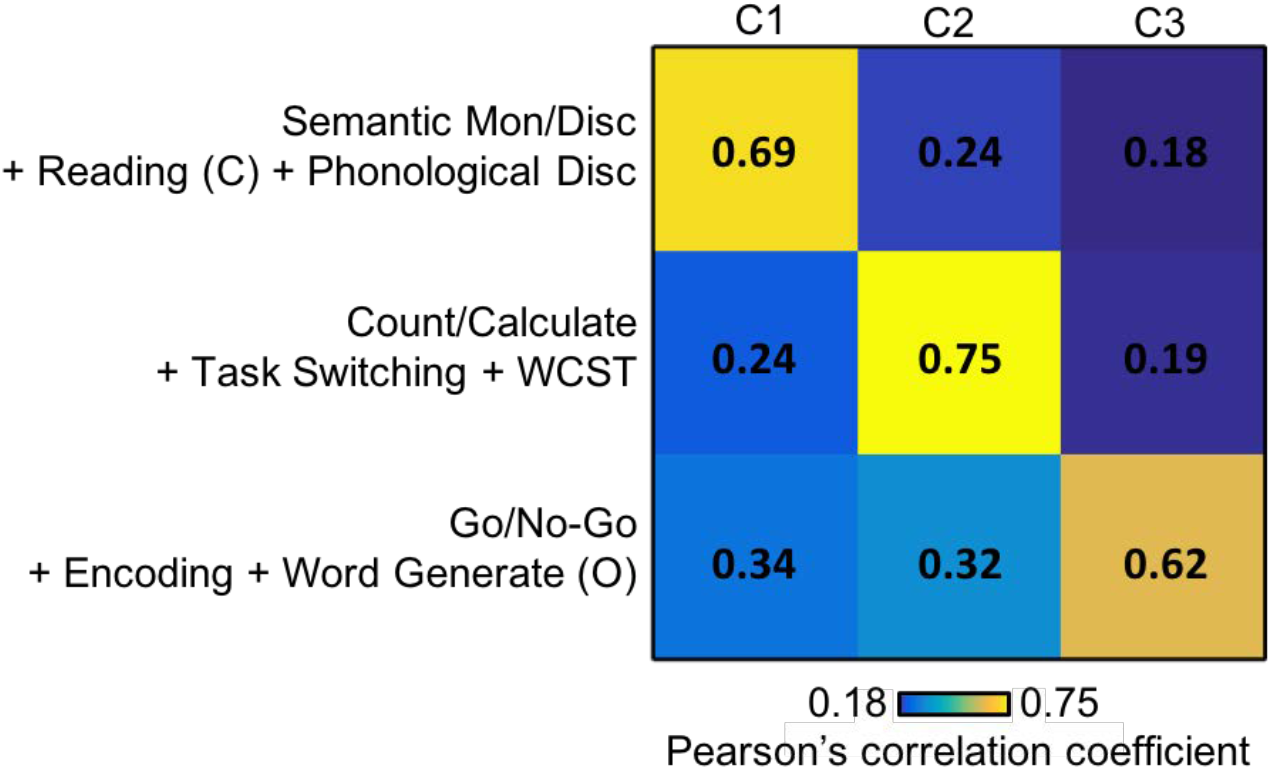
Goodness of fit of the author-topic model for IFJ. The matrix represents the correlations between IFJ’s co-activation patterns (columns) and the average activation maps of the top three tasks associated with each co-activation pattern (rows). The top tasks of each co-activation patterns are shown in Figure 11. The diagonal values (average r = 0.75) were larger than off-diagonal values (average r = 0.31), suggesting a good model fit.

### 3.5 Control analyses

#### 3.5.1 Smoothing

To create the input data for the author-topic model, the activation foci were smoothed with a 10mm binary smoothing kernel (see Section 2.2.3), consistent with previous work (Wager et al. 2003; Yarkoni et al. 2011; Yeo et al. 2015). Using different smoothing radii yielded similar cognitive components of self-generated thought (Figure S5) and co-activation patterns of the IFJ (Figure S6).

#### 3.5.2 Independent component analysis

For comparison, Figure S7 shows the ICA (ICA1-nilearn) estimate of 2 components of self-generated thought. The estimates were quite similar to the author-topic estimate. However, the spatial maps contained negative values, which was inappropriate in the context of coordinate-based meta-analysis.

Figure S8 shows the ICA (ICA1-nilearn) estimate of 3 co-activation patterns of left IFJ. However, the 3 independent components appeared to fractionate the left IFJ into smaller territories (Figure S8), suggesting a degenerate solution to our problem, similar to the situation with the 5-pattern author-topic estimate (Figure S3). Furthermore, both the mixture weights and spatial maps contained negative values, which were not interpretable in the context of coordinate-based meta-analysis (Section 2.3.1).

## 4 Discussion

The author-topic model encodes the intuitive notion that behavioral tasks recruited multiple cognitive components, supported by multiple brain regions (Mesulam 1990; Poldrack 2006; Barrett & Satpute, 2013). We have previously utilized the author-topic model for large-scale meta-analysis across functional domains (Yeo et al., 2015; Bertolero et al., 2015). By exploiting a recently developed CVB algorithm for the author-topic model (Ngo et al., 2016), we show that the model can also be utilized for small-scale meta-analyses focusing on discovering functional sub-domains or task-dependent co-activation patterns.

A dominant approach for small-scale meta-analyses is ALE, which seeks to find consistent activations across neuroimaging experiments within a functional domain or mental disorder or seed region (also known as MACM). ALE treats heterogeneity across experiments as noise. By contrast, the author-topic model evaluates whether the heterogeneity might be indicative of robust latent patterns within the data. We applied the author-topic model to two applications: one on fractionating a functional sub-domain and one on discovering multiple task-dependent co-activation patterns.

In the first application, the author-topic model encoded the notion that tasks involving self-generated thought might recruit one or more spatially overlapping. cognitive components. The model revealed two cognitive components that appeared to delineate two overlapping default sub-networks, consistent with the hypothesized functional organization of the default network (Andres-Hanna et al., 2014). In the second application, the author-topic model encoded the notion that experiments activating a brain region might recruit one or more co-activation patterns dependent on task contexts (McIntosh, 2000). In the current application, the model revealed that the IFJ participated in three co-activation patterns, suggesting that IFJ flexibly co-activate with different brain regions depending on the cognitive demands of different tasks. Overall, our work suggests that the author-topic model is a versatile tool suitable for both small-scale and large-scale meta-analyses.

### 4.1 Cognitive components of self-generated thought

Self-generated thought is a heterogeneous set of cognitive processes that includes inferring other people’s mental states, dealing with challenging moral scenarios, understanding narratives, retrieving autobiographical memories, internalizing semantic information, and mind-wandering. These processes are characterized by an absence of external stimuli, self-related, and often involve simulation or inferential reasoning (Buckner et al., 2008; Spreng et al. 2009; Smallwood et al., 2011; Baird et al., 2011; Prebble at al. 2013; Smallwood, 2013). Studies of tasks involving self-generated thought have consistently found the activation of the default network, suggesting its functional importance (Buckner et al. 2008; Spreng et al. 2009; Andrews-Hanna et al., 2010; Andrews-Hanna, 2012; Gorgolewski et al., 2014; Callard and Margulies, 2014). Additionally, the default network has been fractionated into sub-networks supporting different aspects of these stimulus independent cognitive processes (Buckner et al., 2008; Uddin et al. 2009; Sestieri et al., 2011; Andrews-Hanna et al., 2010; Kim, 2012; Seghier and Price, 2012; Salomon et al., 2013; Bzdok et al., 2013).

The author-topic model revealed two cognitive components of self-generated thought that appeared to fractionate the default network (Figure 7). The default network has been defined as the set of brain regions that are more active during passive task conditions relative to active task conditions (Shulman et al., 1997; Buckner et al., 2008). While there have been multiple studies fractionating the default network (Andrews-Hanna et al., 2010; Mayer et al. 2010; Kim, 2012; Yeo et al. 2014; Humphreys et al., 2015), the specific patterns of fractionation have differed across studies. The spatial topography of components C1 and C2 in this paper corresponded well to the previously proposed “medial temporal subsystem” and “dorsal medial subsystem” respectively (Figure 3A of Andrews-Hanna et al. 2014; Andrews-Hanna et al., 2010).

The first cognitive component C1 was strongly recruited by navigation and autobiographical memory tasks, suggesting its involvement in constructive mental simulation based upon mnemonic content (Andrews-Hanna et al., 2014). Constructive mental simulation is the process of combining information from the past in order to create a novel mental representation, such as imagining the future (Buckner and Carroll, 2007; Hassabis and Maguire, 2007; Schacter et al., 2007). “Navigation” tasks require constructive mental simulation to create a mental visualization (“scene construction”) for planning new routes and finding ways in unfamiliar contexts (Burgess et al., 2002; Byrne et al. 2007). On the other hand, “Autobiographical Memory” tasks require constructive mental simulation to project past experience (“constructive episodic simulation”; Atance and O’Neil, 2001; Schacter et al. 2007) or previously acquired knowledge (“semantic memory”; Irish et al., 2012; Brown et al. 2014) across spatiotemporal scale to enact novel perspectives. Overall, cognitive component C1 seems to support the projection of self, events, experiences, images and knowledge to a new temporal or spatial context based upon an associative constructive process, likely mediated by the hippocampus and connected brain structures (Moscovitch et al., 2016, Christoff et al., 2016).

The second cognitive component C2 was strongly recruited by narrative comprehension and theory of mind, suggesting its involvement in mentalizing, inferential, and conceptual processing (Andrews-Hanna et al., 2014). Mentalizing is the process of monitoring one’s own mental states or predicting others’ mental states (Frith and Frith, 2003), while conceptual processing involves internalizing and retrieving semantic or social knowledge (Binder and Desai, 2011; Overwalle, 2009). “Narrative Comprehension” engages conceptual processing to understand the contextual settings of the story, and requires mentalizing to follow and infer the characters’ thoughts and emotions (Gernsbacher et al., 1998; Mason et al. 2008). “Theory of Mind” tasks require the recall of learned knowledge, social norms and attitudes to form a meta-representation of the perspectives of other people (Leslie, 1987; Frith and Frith, 2005; Binder and Desai, 2011). The grouping of Narrative Comprehension and Theory of Mind tasks echoes the link between the ability to comprehend narratives and the ability to understanding others’ thoughts in developmental studies of children (Guajardo and Watson, 2001; Slaughter et al. 2007; Mason et al. 2008).

The two cognitive components had high probability of activating common and distinct brain regions. Both components engaged the posterior cingulate cortex and precuneus, which are considered part of the “core” sub-network that subserves personally relevant information necessary for both constructive mental simulation and mentalizing (Andrews-Hanna et al. 2014). The distinct brain regions supporting each cognitive component also corroborated the distinct functional role of each component. For instance, component C1, but not C2, had high probability of activating the medial temporal lobe and hippocampus. This is consistent with neuropsychological literaure showing that patients with impairment of the medial temporal lobe and hippocampus retain theory of mind and narrative construction capabilities, while suffering deficits in episodic memories and imagining the future (Hassabis et al., 2007; Rosenbaum et al., 2007; Rosenbaum et al., 2009; Race et al., 2011;).

The cognitive components of self-generated thought estimated by the author-topic model overlapped with default sub-networks A, B and C, as well as the temporal parietal network from a previously published resting-state parcellation (Yeo et al., 2011; Kong et al., 2018). The components loaded differentially on the resting-state networks, thus providing insights into the functions of distinct resting-state networks. Although resting-state fMRI is a powerful tool for extracting brain networks, participants do not actively perform a task during resting-state fMRI. Thus, coordinate-based meta-analysis can be used in conjunction with resting-state fMRI to discover new insights into brain networks and their functions (Seeley et al., 2007; Smith et al., 2009; Laird et al., 2011).

### 4.2 Co-activation patterns of the left IFJ

The inferior frontal junction (IFJ) is located in the prefrontal cortex at the intersection between the inferior frontal sulcus and the inferior precentral sulcus (Brass et al., 2005; Derrfuss et al., 2005). The IFJ has been suggested to be involved in a wide range of cognitive functions, including task switching (Brass and Cramon, 2002; Derrfuss et al., 2004, 2005), attentional control (Asplund et al. 2010; Baldauf and Desimone, 2014), detection of conflicting responses (Chikazoe et al. 2009; Levy and Wagner, 2011), short-term memory (Zanto et al. 2010; Sneve et al. 2013), construction of attentional episodes (Duncan, 2013) and so on. Using the author-topic model, we found that the IFJ participated in three task-dependent co-activation patterns.

*Co-activation pattern C1* might be involved in some aspects of language processing, such as phonological processing for lexical understanding. Phonological processing is an important linguistic function, concerning the use of speech sounds in handling written or oral languages (Wagner and Torgesen 1987; Poldrack et al. 1999; Friederici 2002). The top tasks associated with C1 were “Semantic Monitoring/ Discrimination”, “Covert Reading”, and “Phonological Discrimination” (Figure 11B). Inspecting the top three experiments recruiting these three tasks (Table S3-A) offered more insights into the functional characteristics of co-activation pattern C1. The top “Semantic Monitor/Discrimination” experiments with the highest probability of recruiting co-activation pattern C1 examined retrieval of semantic meaning (Thompson-Schill et al. 1999; Wagner et al. 2001) and an experiment requiring lexical perception and not just perception of elementary sounds (Poeppel et al., 2004). The top “Covert Reading” experiments most strongly associated with C1 identified a common brain network activated by both reading and listening (Jobard et al., 2007), as well as language comprehension across different media (Small et al., 2009), suggesting the involvement of C1 in generic language comprehension. Among “Phonological Discrimination” experiments, C1 was most highly associated with experiments engaging transcoding of phonological representation for semantic perception (Xu et al. 2001; Démonet et al. 1994). The language and phonological processing interpretation was supported by C1’s strong left lateralization with high probability of activating classical auditory and language brain regions, including the left (but not right) inferior frontal gyrus and bilateral superior temporal cortex.

*Co-activation pattern C2* might be engaged in attentional control, especially aspects of task maintenance and shifting of attentional set. Attentional set-shifting is the ability to switch between mental states associated with different reactionary tendencies (Omori et al. 1999, Konishi et al. 1998). The top three tasks most highly associated with C2 were “Counting/Calculation”, “Task Switching”, and “Wisconsin Card Sorting Test” (Figure 11B). Inspecting the top three experiments under the top task paradigms provided further insights into the functional characteristics of co-activation pattern C2 (Table S3-B). The top “Counting/ Calculation” experiments most strongly recruiting co-activation pattern C2 involved switching of resolution strategies in executive function. For example, one experimental contrast seeks to isolate demanding mental calculation but not retrieval of numerical facts (Zago et al. 2001; Rivera et al. 2002), suggesting C2’s involvement in the selection and application of strategies to solve arithmetic problems. The top “Task Switching” experiments most strongly associated with C2 involved the switching of mental states to learn new stimulus-response or stimulus-outcome associations (Omori et al. 1999; Nahagama et al. 2001; Sylvester et al. 2003). C2 was also strongly expressed by “Wisconsin Card Sorting Test” (WCST) experiments, which required attentional set-shifting to change behavioral patterns in reaction to changes of perceptual dimension (color, shape, or number) upon which the target and reference stimuli were matched (Berman 1995; Konishi 2002; Konishi 2003). Overall, the attentional control interpretation of co-activation pattern C2 is supported by C2’s high probability of activating classical attentional control regions, such as the superior parietal lobule and the intra-parietal sulcus, although there is a clear lack of DLPFC activation.

*Co-activation pattern C3* might be engaged in inhibition or response conflict resolution. Conflict-response resolution is a central aspect of cognitive control, which involves monitoring and mediating incongruous response tendencies (Pardo et al. 1990; Braver et al. 2001; Barch et al. 2001). Co-activation pattern C3 is most strongly recruited by experiments utilizing “Go/No-Go”, “Encoding” and “Overt Word Generation” tasks (Figure 11B). Closer examination of the top three experiments under each task paradigm provided further insights into the functional characteristics of C3 (Table S3-C). The top experiments utilizing “Go/No-Go” required the monitoring, preparing and reconciling of conflicting tendencies to either giving a “go” or “stop” (no-go) response (Chikazoe et al. 2009, Simoes-Franklin et al. 2010; Kawashima et al. 1996). It might be surprising at first glance that the “Go/No-Go” task was grouped together with “Encoding” and “Overt Word Generation” tasks. However, the top experiments utilizing the “Encoding” and “Overt Word Generation” task all required subjects to make competing decisions (Table S3-C). The top “Encoding” experiments most strongly associated with C3 required selective association of to-be-learnt items with existing memory or knowledge organization for effective enduring retention of new information (Kapur et al. 1996; Callan et al. 2010; Mickley et al. 2009). The top experiments utilizing the “Overt Word Generation” task required subjects to make competing decision, such as inhibiting verbalization of wrong words in verbal fluency task (Baker et al. 1997; Ravnkilde et al. 2002) or inhibiting a predominant pattern (regular past-tense verbs) in favor of generating less conventional forms (irregular past-tense verbs) (Desai et al. 2006). Overall, the inhibition or response conflict interpretation of co-activation pattern C3 is supported by C3’s high probability of activating classical executive function regions, including the bilateral dorsal lateral prefrontal cortex.

The intriguing location of the left IFJ and its functional heterogeneity suggests the role of IFJ as an integrative hub for different cognitive functions. For example, the IFJ has been suggested to consolidate information streams for cognitive control from its bordering brain regions (Brass et al., 2005). The involvement of the IFJ in three task-dependent co-activation patterns supported the view that the IFJ orchestrates different cognitive mechanisms to allow their operations in harmony.

## 5 Conclusion

Heterogeneities across neuroimaging experiments are often treated as noise in coordinate-based meta-analyses. Here we demonstrate that the author-topic model can be utilized to determine if the heterogeneities can be explained by a small number of latent patterns. In the first application, the author-topic model revealed two overlapping cognitive components subserving self-generated thought. In the second application, the author-topic revealed the participation of the left IFJ in three task-dependent co-activation patterns. These applications exhibited the broad utility of the author-topic model, ranging from discovering functional subdomains or task-dependent co-activation patterns.

## Acknowledgements

This work was supported by Singapore MOE Tier 2 (MOE2014-T2-2-016), NUS Strategic Research (DPRT/944/09/14), NUS SOM Aspiration Fund (R185000271720), Singapore NMRC (CBRG/0088/2015), NUS YIA and the Singapore National Research Foundation (NRF) Fellowship (Class of 2017). Simon Eickhoff is supported by the National Institute of Mental Health (R01-MH074457), the Helmholtz Portfolio Theme “Supercomputing and Modeling for the Human Brain” and the European Union’s Horizon 2020 Research and Innovation Programme under Grant Agreement No. 7202070 (HBP SGA1). R. Nathan Spreng is supported by the National Science and Engineering Research Council of Canada, the Canadian Institutes of Health Research, and received salary support from the Fonds de la Recherche du Quebec – Santé (FRQS). Comprehensive access to the BrainMap database was authorized by a collaborative-use license agreement (http://www.brainmap.org/collaborations.html). BrainMap database development is supported by NIH/NIMH R01 MH074457. Our computational work was partially performed on resources of the National Supercomputing Centre, Singapore (https://www.nscc.sg).

## 6 Appendix

| Pseudo-code | Function |
| --- | --- |
| (1) Read the input activation foci and task labels | CBIG_AuthorTopic_PreprocessInput |
| (2) Initialize the model's hyperparameters | CBIG_AuthorTopic_SetupParameters |
| (3) Repeat for $N = 1000$ re-initializations | |
| (a) Initialize the variational parameters $\phi$ | CBIG_AuthorTopic_InitializeParams |
| (b) Update the variational parameters $\phi$ (Eq. 14 in Supplemental S1) | |
| - Approximate $E_q(N_{..c.}^{-ef})$ and $Var_q(N_{..c.}^{-ef})$ | CBIG_AuthorTopic_ComputeVariationalTerm__N_c |
| - Approximate $E_q(N_{..cv_{ef}}^{-ef})$ and $Var_q(N_{..cv_{ef}}^{-ef})$ | CBIG_AuthorTopic_ComputeVariationalTerm__N_cv |
| - Approximate $E_q(N_{.t..}^{-ef})$ and $Var_q(N_{.t..}^{-ef})$ | CBIG_AuthorTopic_ComputeVariationalTerm__N_t |
| - Approximate $E_q(N_{.tc.}^{-ef})$ and $Var_q(N_{.tc.}^{-ef})$ | CBIG_AuthorTopic_ComputeVariationalTerm__N_tc |
| - Update the variational parameters $\phi$ (Eq. 14 in Supplemental S1) | |
| - Recompute the variational log likelihood. If it converges, go to step (3c), otherwise repeat from step (3b) |  |
| (c) Update the model parameters $\theta_{tc}$ and $\beta_{cv}$ (Eq. 16 and 17 in Supplemental S1) | CBIG_AuthorTopic_EstimateParams |

Pseudo-code of the Collapsed Variational Bayes (CVB) algorithm for estimating the author-topic model’s parameters. The left column outlines the main steps of the algorithm. The right column denotes the functions in the source code that correspond to each step. The source code of the CVB algorithm and the input file of the self-generated thought dataset are available at https://github.com/ThomasYeoLab/CBIG/tree/master/stable_projects/meta-analysis/Ngo2019_AuthorTopic. Note that steps (2) and (3) can be called by a single function *CBIG_AuthorTopic_RunInference.*

## Supplemental Material

This supplemental material is divided into *Supplemental Results* and *Supplemental Methods* to complement Results and Methods section in the main text, respectively.

### 1. Supplemental Results

#### 1.1 Supplemental Result Tables

**1.1.1 Table S1.**
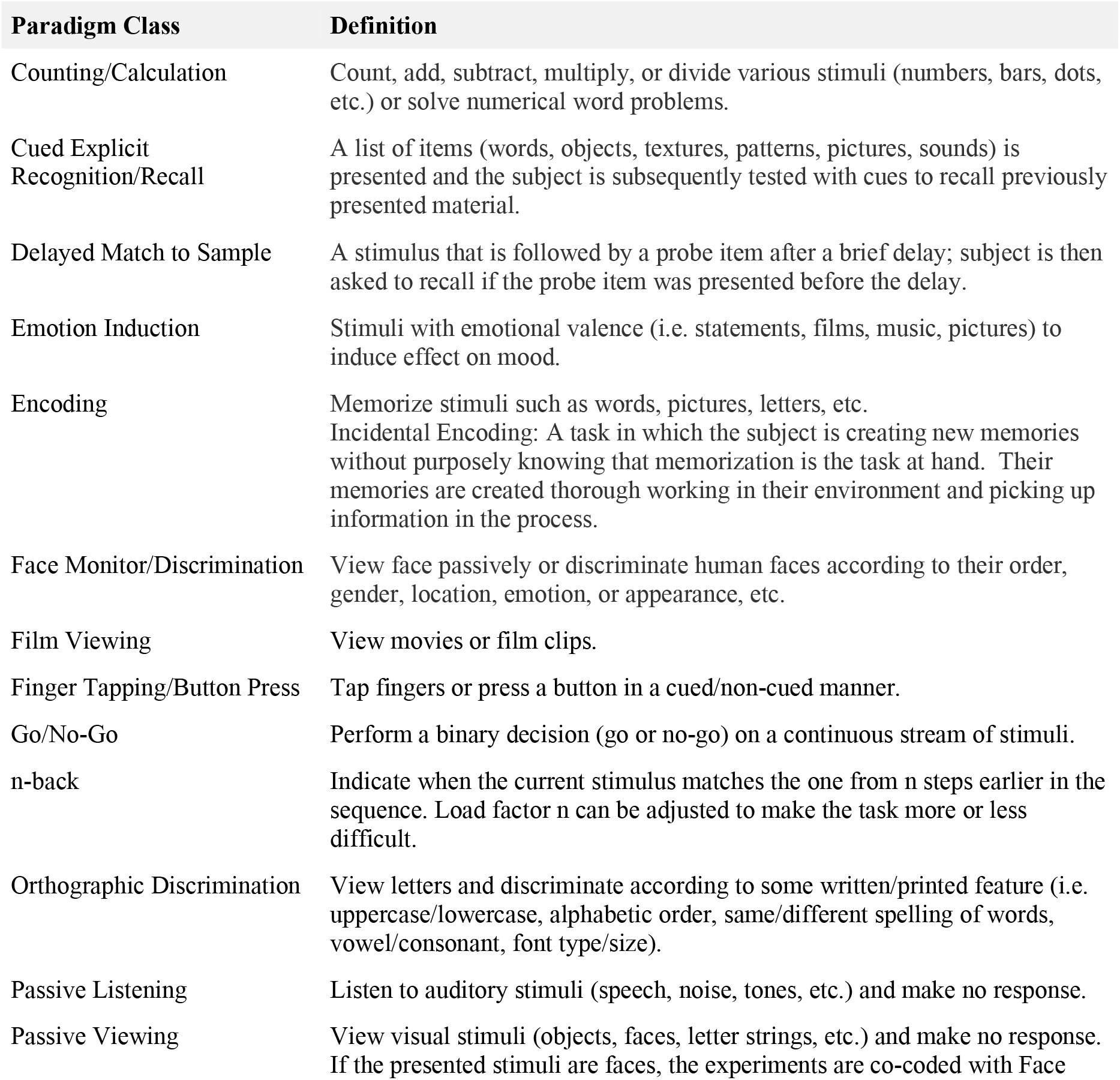
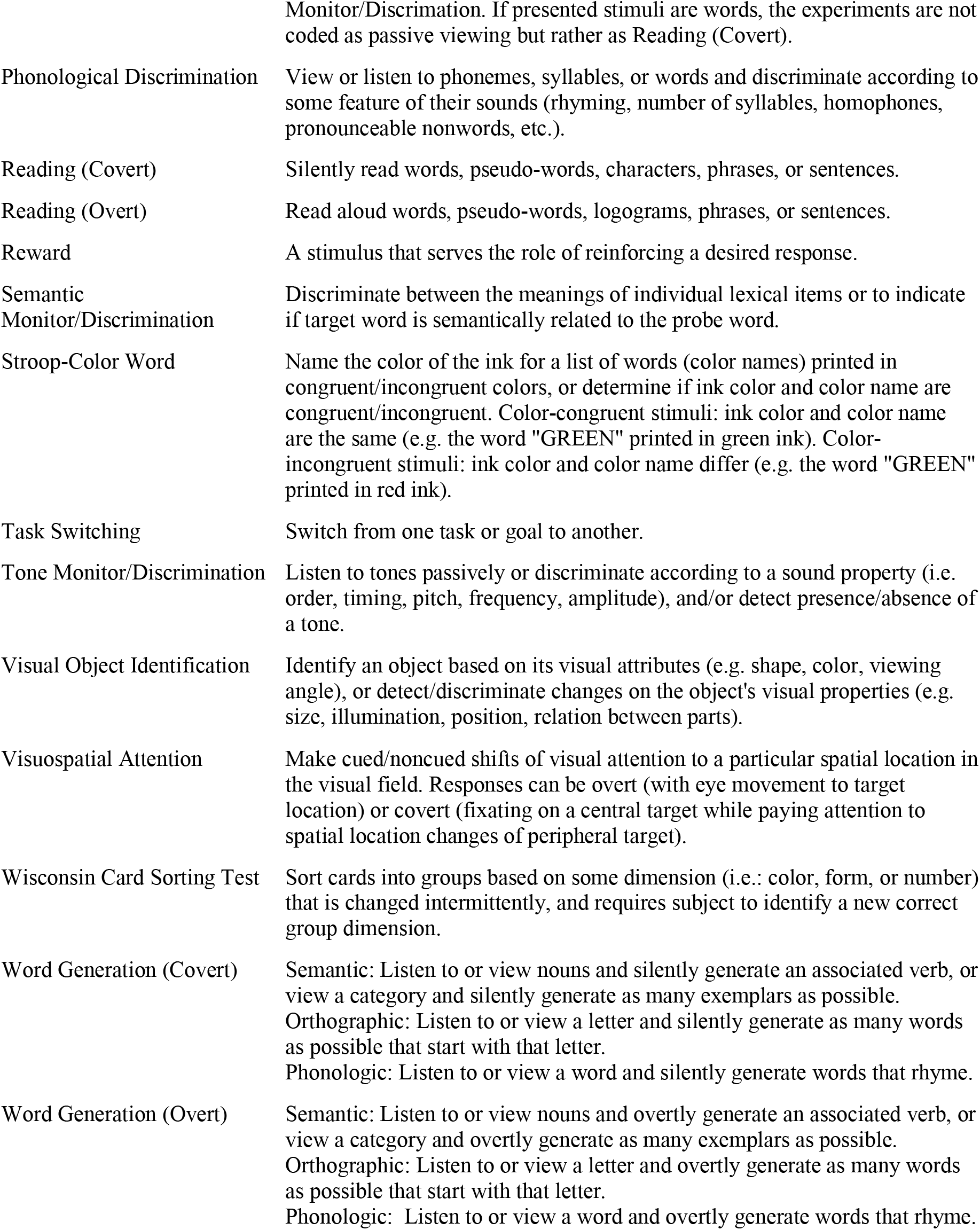
Definitions of 26 BrainMap task categories (paradigm classes) tagged to experiments activating the IFJ. Only paradigm classes tagged to at least 5 experiments were included. The definitions were extracted from BrainMap lexicon, available at http://www.brainmap.org/scribe/BrainMapLex.xls

**1.1.2 Table S2.** Top 5 tasks with the highest probabilities of recruiting a co-activation pattern involving the IFJ

| **Co-activation Pattern C1** |  |
| --- | --- |
| **Task** | **Pr (C1 \| Task)** |
| Semantic Monitor/Discrimination | 0.51 |
| Reading (Covert) | 0.50 |
| Phonological Discrimination | 0.49 |
| Film Viewing | 0.45 |
| Word Generation (Overt) | 0.42 |
| **Co-activation Pattern C2** |  |
| **Task** | **Pr (C2 \| Task)** |
| Counting/Calculation | 0.63 |
| Task Switching | 0.60 |
| Wisconsin Card Sorting Task | 0.60 |
| Stroop-Color Word | 0.49 |
| Delayed Match to Sample | 0.46 |
| **Co-activation Pattern C3** |  |
| **Task** | **Pr (C3 \| Task)** |
| Go/ No-Go | 0.54 |
| Encoding | 0.44 |
| Word Generation (Overt) | 0.42 |
| Reward | 0.41 |
| Reading (Overt) | 0.36 |

**1.1.3 Table S3-A.** Experiments with the highest probabilities of recruiting co-activation pattern C1 of IFJ

| **Semantic Monitor/Discrimination** |  |  |  |  |  |
| --- | --- | --- | --- | --- | --- |
| **PMID** | **Author** | **Title** | **Exp** | **Contrast** | **Pr** |
| 10390028 | Thompson-Schill 1999 | A neural basis for category and modality specificity of semantic knowledge | 3 | Retrieval of visual knowledge about nonliving things - baseline | 0.81 |
| 14644105 | Poeppel 2004 | Auditory lexical decision, categorical perception, and FM direction discrimination differentially engage left and right auditory cortex | 5 | Lexical decision of phonologically permissible targets vs. frequency-modulated signals | 0.80 |
| 11502262 | Wagner 2001 | Recovering meaning: Left prefrontal cortex guides controlled semantic retrieval | 4 | 4 target items > 2 target items (weak cue-correct response association) | 0.79 |
| **Reading (Covert)** |  |  |  |  |  |
| 17110132 | Jobard 2007 | Impact of modality and linguistic complexity during reading and listening tasks | 3 | Conjunction of reading and listening irrespective of the linguistic complexity | 0.87 |
| 19155745 | Small 2009 | Your brain on Google: patterns of cerebral activation during internet searching | 3 | Internet task in Internet-savvy group | 0.72 |
| 19155745 | Small 2009 | Your brain on Google: patterns of cerebral activation during internet searching | 2 | Reading task in Internet-naïve group | 0.64 |
| **Phonological Discrimination** |  |  |  |  |  |
| 11230098 | Xu 2001 | Conjoint and extended neural networks for the computation of speech codes: The neural basis of selective impairment in reading words and pseudowords | 4 | Real-word rhyming – color-matching with letters (Group) | 0.86 |
| 7922456 | Démonet 1994 | A PET study of cognitive strategies in normal subjects during language tasks. Influence of phonetic ambiguity and sequence processing on phoneme monitoring | 4 | Phoneme monitoring of perceptually ambiguous sequence of non-words – monitoring pure tones | 0.85 |
| 11230098 | Xu 2001 | Conjoint and extended neural networks for the computation of speech codes: The neural basis of selective impairment in reading words and pseudowords | 7 | Alternate-case rhyming – color-matching with letters (Group) | 0.83 |

Each row shows one of the top 3 experiments with the highest probabilities of recruiting co-activation pattern C1 of IFJ and employing one of the top 3 tasks with the highest probabilities of recruiting C1, namely “Semantic Monitoring/Discrimination”, “Covert Reading”, and “Phonological Discrimination”. The “PMID” and “Title” columns list the PubMed ID and title of each study respectively. The “Author” column lists the last name of the first author and the year of publication of each study. The “Exp” column lists the experiment’s order in the respective study as reported in BrainMap. The “Contrast” column lists the experimental contrast of each experiment. The “Pr” column shows the probability that each experiment would recruit co-activation pattern C1.

**1.1.4 Table S3-B.**
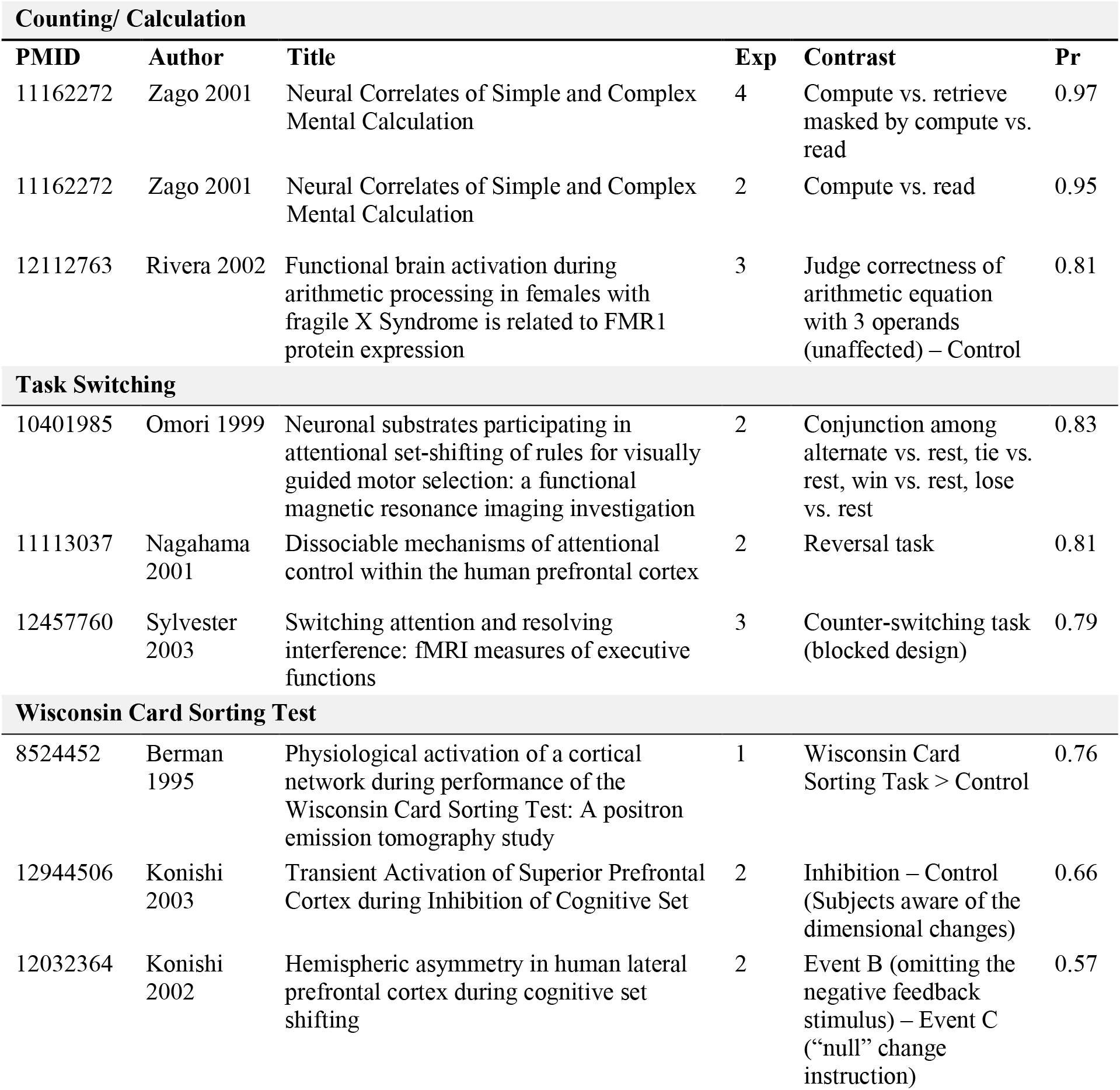
Experiments with the highest probabilities of recruiting co-activation pattern C2 of IFJ

| Counting/ Calculation |  |  |  |  |  |
| --- | --- | --- | --- | --- | --- |
| PMID | Author | Title | Exp | Contrast | Pr |
| 11162272 | Zago 2001 | Neural Correlates of Simple and Complex Mental Calculation | 4 | Compute vs. retrieve masked by compute vs. read | 0.97 |
| 11162272 | Zago 2001 | Neural Correlates of Simple and Complex Mental Calculation | 2 | Compute vs. read | 0.95 |
| 12112763 | Rivera 2002 | Functional brain activation during arithmetic processing in females with fragile X Syndrome is related to FMR1 protein expression | 3 | Judge correctness of arithmetic equation with 3 operands (unaffected) – Control | 0.81 |
| Task Switching |  |  |  |  |  |
| 10401985 | Omori 1999 | Neuronal substrates participating in attentional set-shifting of rules for visually guided motor selection: a functional magnetic resonance imaging investigation | 2 | Conjunction among alternate vs. rest, tie vs. rest, win vs. rest, lose vs. rest | 0.83 |
| 11113037 | Nagahama 2001 | Dissociable mechanisms of attentional control within the human prefrontal cortex | 2 | Reversal task | 0.81 |
| 12457760 | Sylvester 2003 | Switching attention and resolving interference: fMRI measures of executive functions | 3 | Counter-switching task (blocked design) | 0.79 |
| Wisconsin Card Sorting Test |  |  |  |  |  |
| 8524452 | Berman 1995 | Physiological activation of a cortical network during performance of the Wisconsin Card Sorting Test: A positron emission tomography study | 1 | Wisconsin Card Sorting Task > Control | 0.76 |
| 12944506 | Konishi 2003 | Transient Activation of Superior Prefrontal Cortex during Inhibition of Cognitive Set | 2 | Inhibition – Control (Subjects aware of the dimensional changes) | 0.66 |
| 12032364 | Konishi 2002 | Hemispheric asymmetry in human lateral prefrontal cortex during cognitive set shifting | 2 | Event B (omitting the negative feedback stimulus) – Event C (“null” change instruction) | 0.57 |

Each row shows one of the top 3 experiments with the highest probabilities of recruiting co-activation pattern C2 of IFJ and employing one of the top 3 tasks with the highest probabilities of recruiting C2, namely “Counting/Calculation”, “Task Switching”, and “Wisconsin Card Sorting Test”. The “PMID” and “Title” columns list the PubMed ID and title of each study respectively. The “Author” column lists the last name of the first author and the year of publication of each study. The “Exp” column lists the experiment’s order in the respective study as reported in BrainMap. The “Contrast” column lists the experimental contrast of each experiment. The “Pr” column shows the probability that each experiment would recruit co-activation pattern C2.

**1.1.5 Table S3-C.**
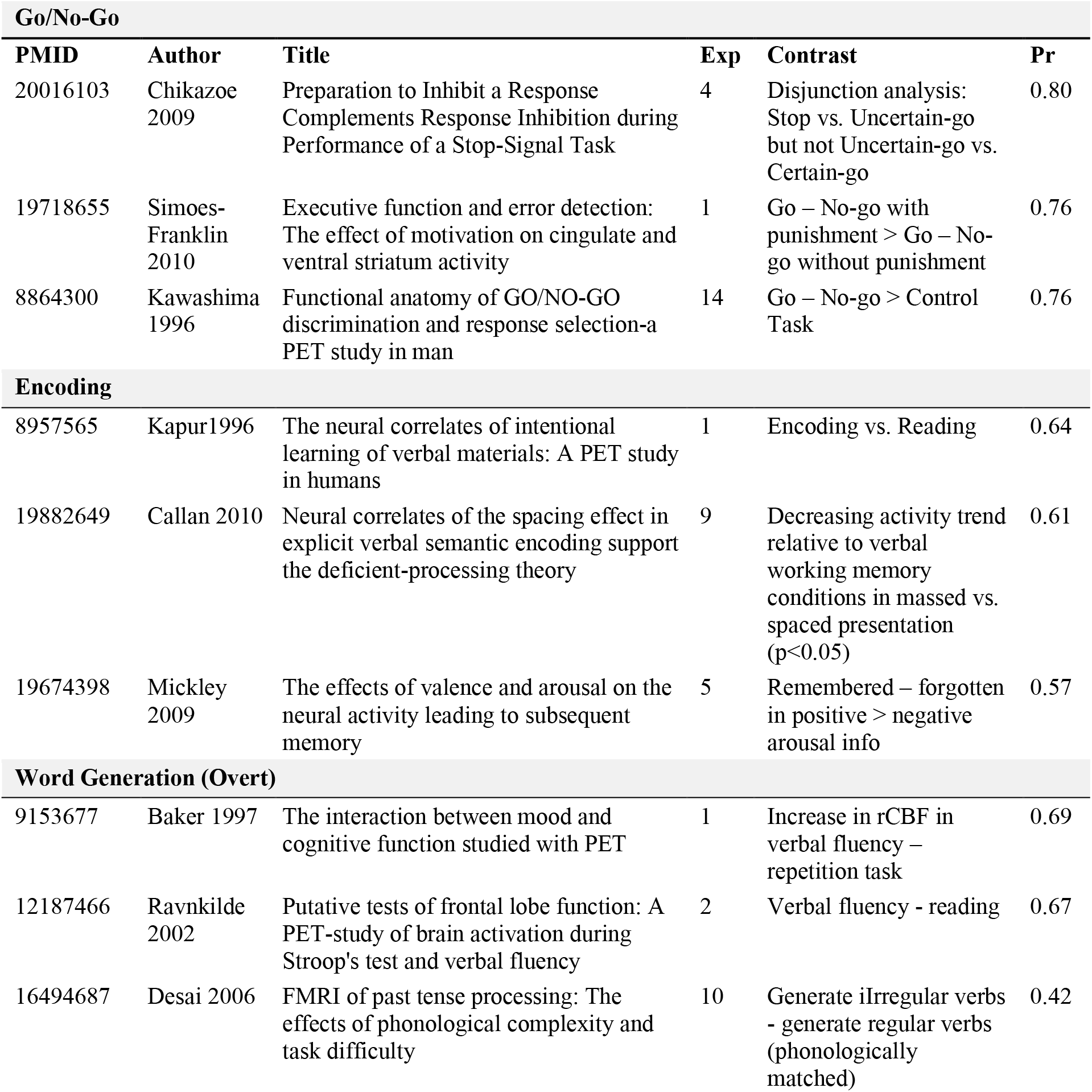
Experiments with the highest probabilities of recruiting co-activation pattern C3 of IFJ

Each row shows one of the top 3 experiments with the highest probabilities of recruiting co-activation pattern C3 of IFJ and employing one of the top 3 tasks with the highest probabilities of recruiting C3, namely “Go/No-Go”, “Encoding”, and “Overt Word Generation”. The “PMID” and “Title” columns list the PubMed ID and title of each study respectively. The “Author” column lists the last name of the first author and the year of publication of each study. The “Exp” column lists the experiment’s order in the respective study as reported in BrainMap. The “Contrast” column lists the experimental contrast of each experiment. The “Pr” column shows the probability that each experiment would recruit co-activation pattern C3.

#### 1.2 Supplemental Result Figures

**Figure S1.**
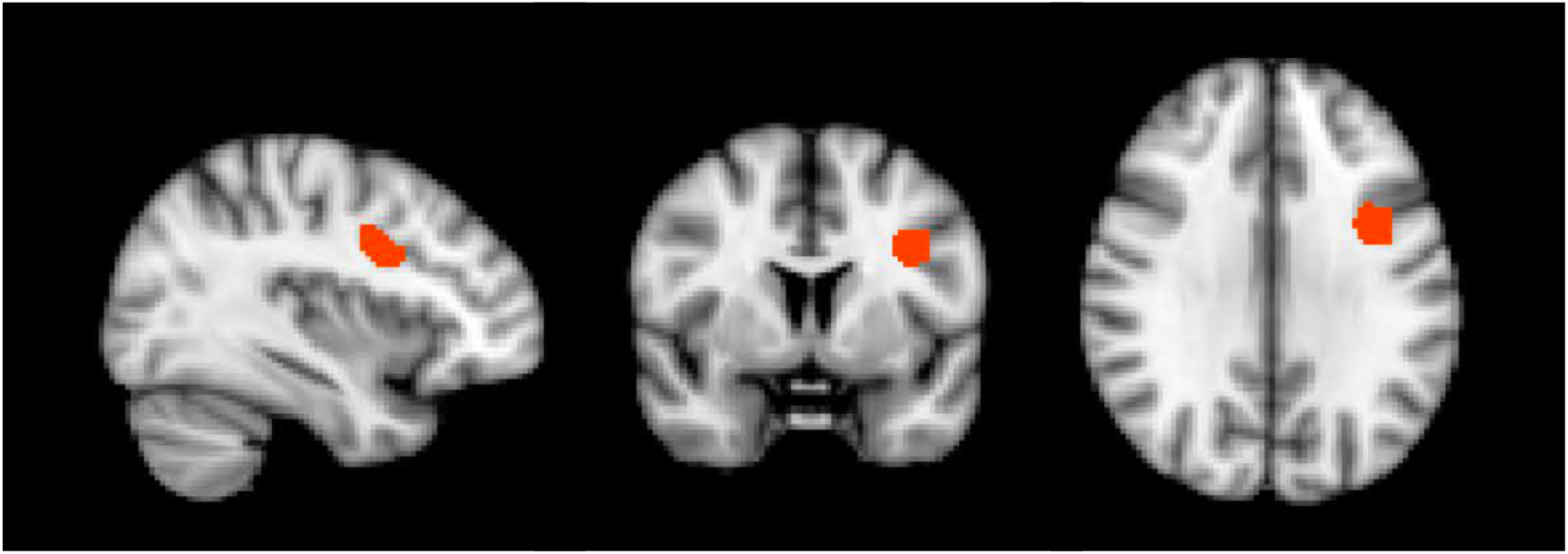
Illustration of the left IFJ seed region (Muhle-Karbe et al., 2015) in the sagittal, coronal and axial planes.

**Figure S2.**
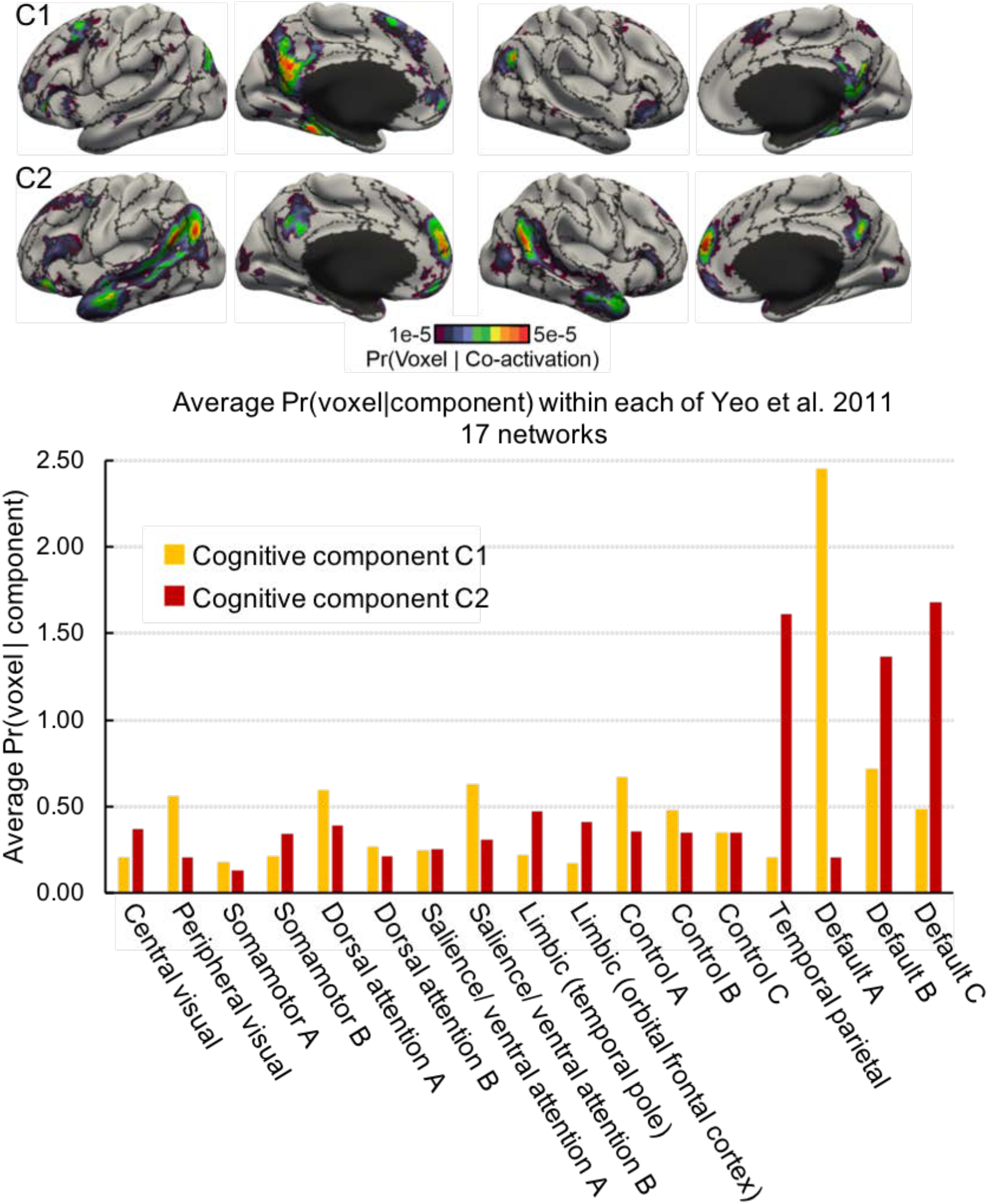
Comparison of the cognitive components of self-generated thought with Yeo et al. (2011) 17-network parcellation. The top figure shows the surface-based visualization of the probability of cognitive components of self-generated thought activating different brain voxels (i.e., Pr(voxel | component)) overlaid on top of the 17-network boundaries (black lines) from Yeo et al. (2011). The bottom bar chart shows the average Pr(voxel | component) of each network. The yellow and red columns correspond to cognitive components C1 and C2 respectively.

**Figure S3.**
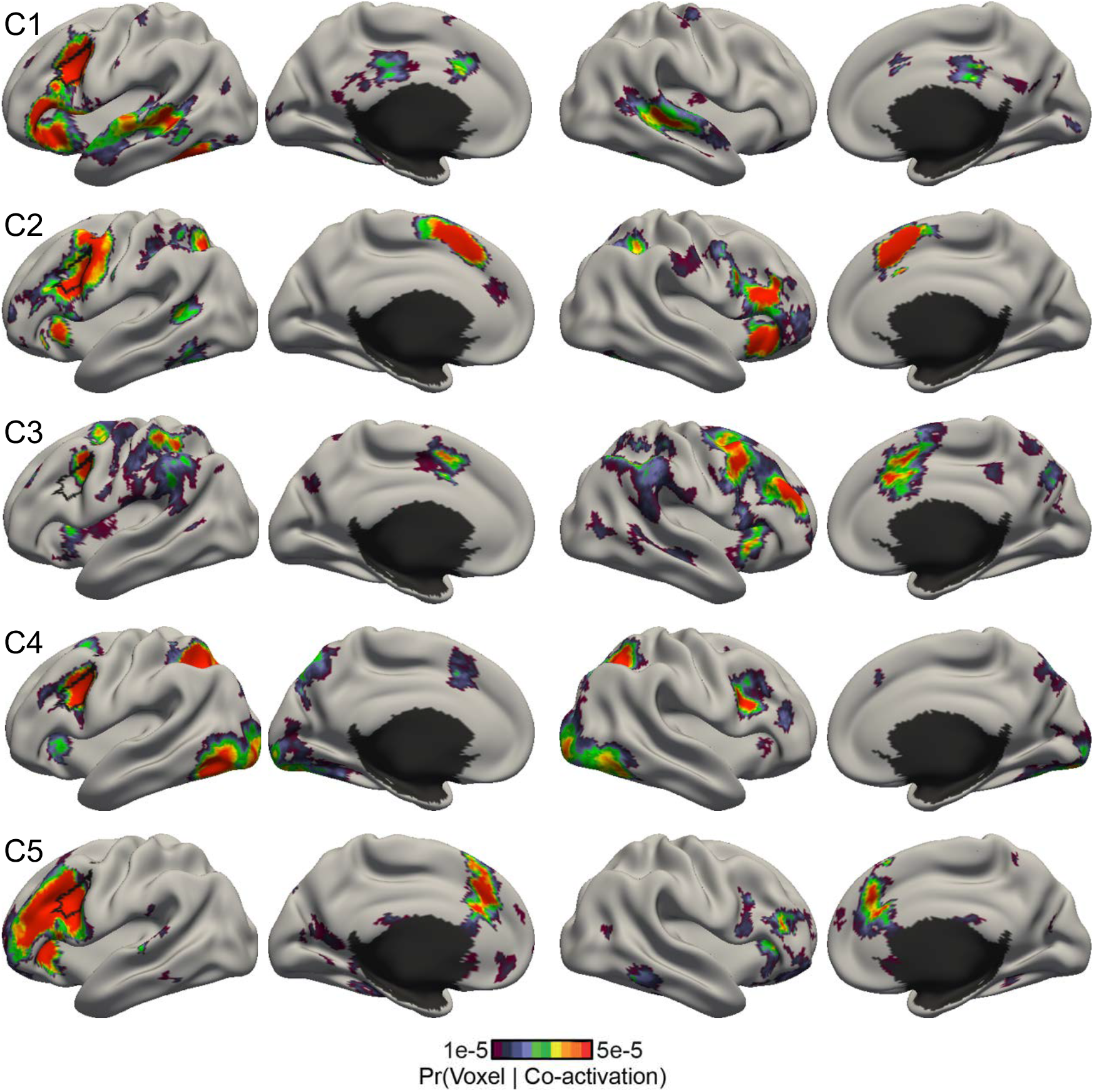
Model estimates of 5 co-activation patterns involving the left inferior frontal junction (IFJ). Each row shows the surface-based visualization for the probability of a co-activation pattern recruiting different brain voxels (i.e., Pr(voxel | co-activation pattern)). Co-activation pattern C3 only involves a part of the left IFJ, suggesting that the 5-pattern model estimates might fractionate the seed region.

**Figure S4.**
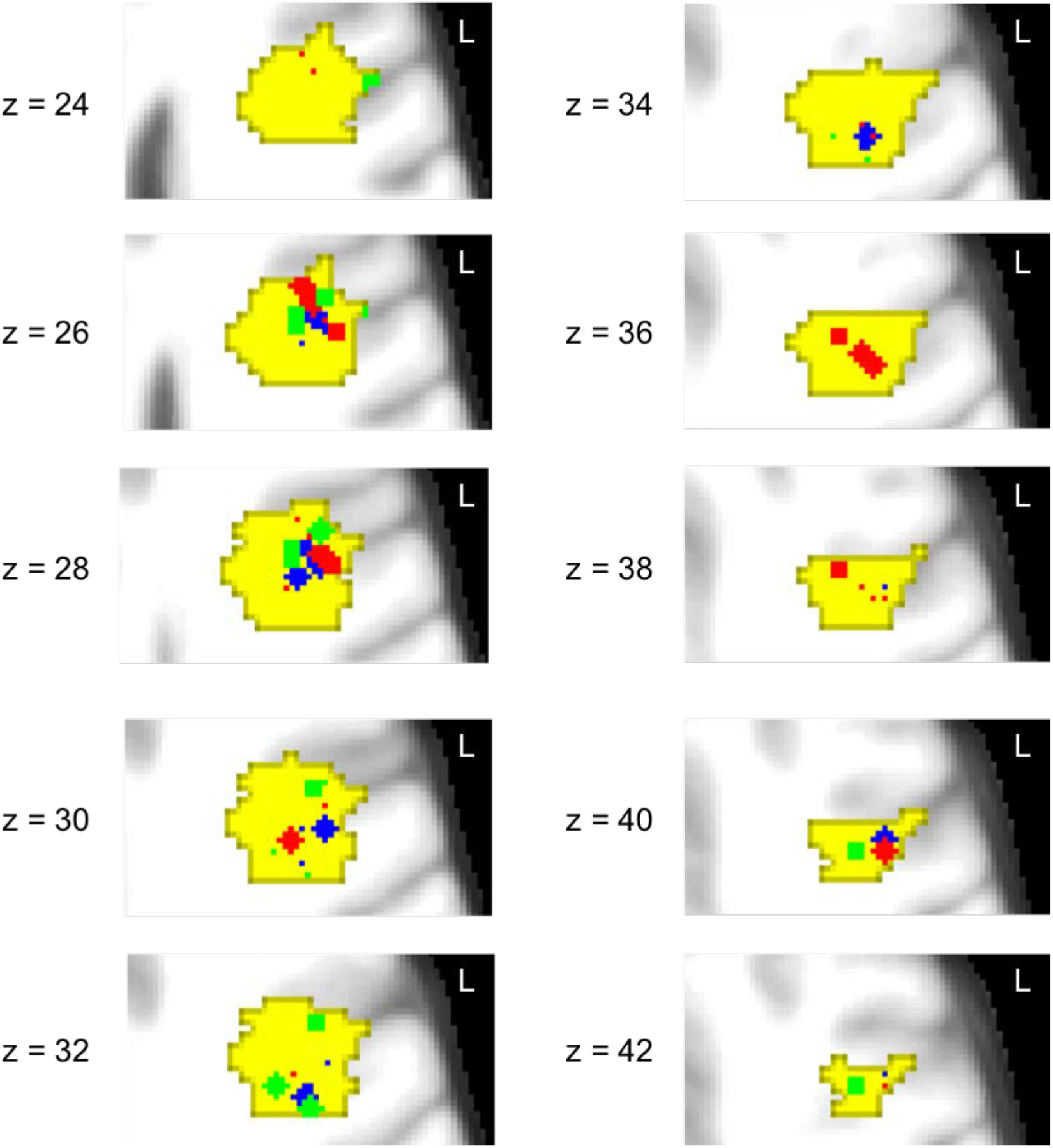
Volumetric slices suggesting the foci of co-activation patterns overlap within the left IFJ. Yellow indicates the left IFJ seed region. The colored dots correspond 2-mm-radius spheres centered about the activation foci reported by the top 3 experiments with the highest probabilities of recruiting IFJ co-activation patterns and employs one of the top 3 tasks with the highest probabilities of recruiting the given co-activation pattern. Blue, red and green dots correspond to the activation foci associated with co-activation patterns C1, C2, and C3, respectively.

**Figure S5.**
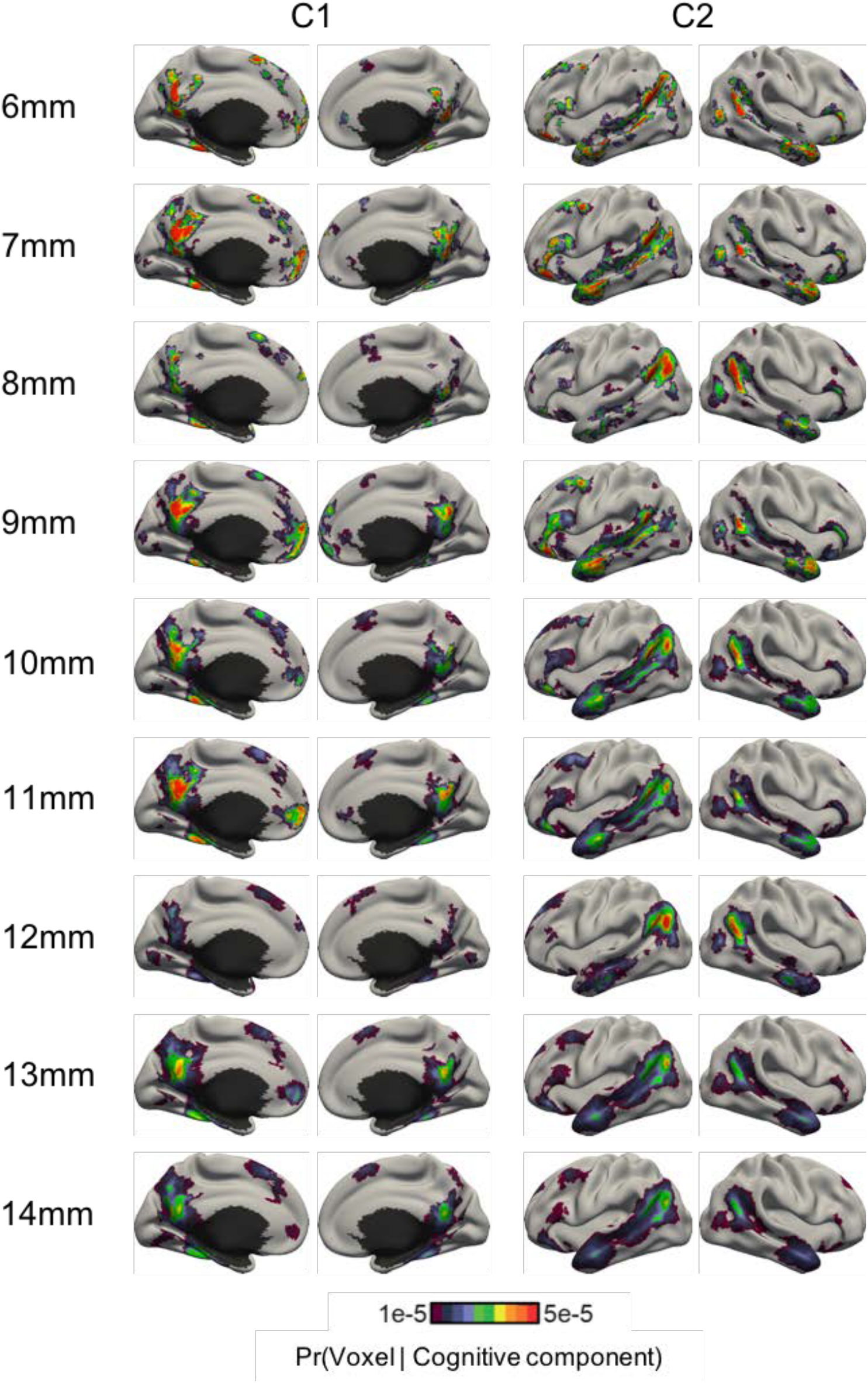
Cognitive components of self-generated thought estimated by the author-topic model using different radii of smoothing kernel applied to the activation foci. The smoothing radii range from 6mm to 14mm. Cognitive components estimated with different smoothing kernel radii are similar to the components estimated with a 10mm-radius smoothing kernel. The average Pearsons’ correlation coefficient of the cognitive components estimated with different smoothing kernel radii against those estimated with a 10mm-radius ranges from 0.60 (6mm) to 0.79 (14mm). The cognitive components estimated with a 10mm-radius smoothing kernel were shown in all results and analyses.

**Figure S6.**
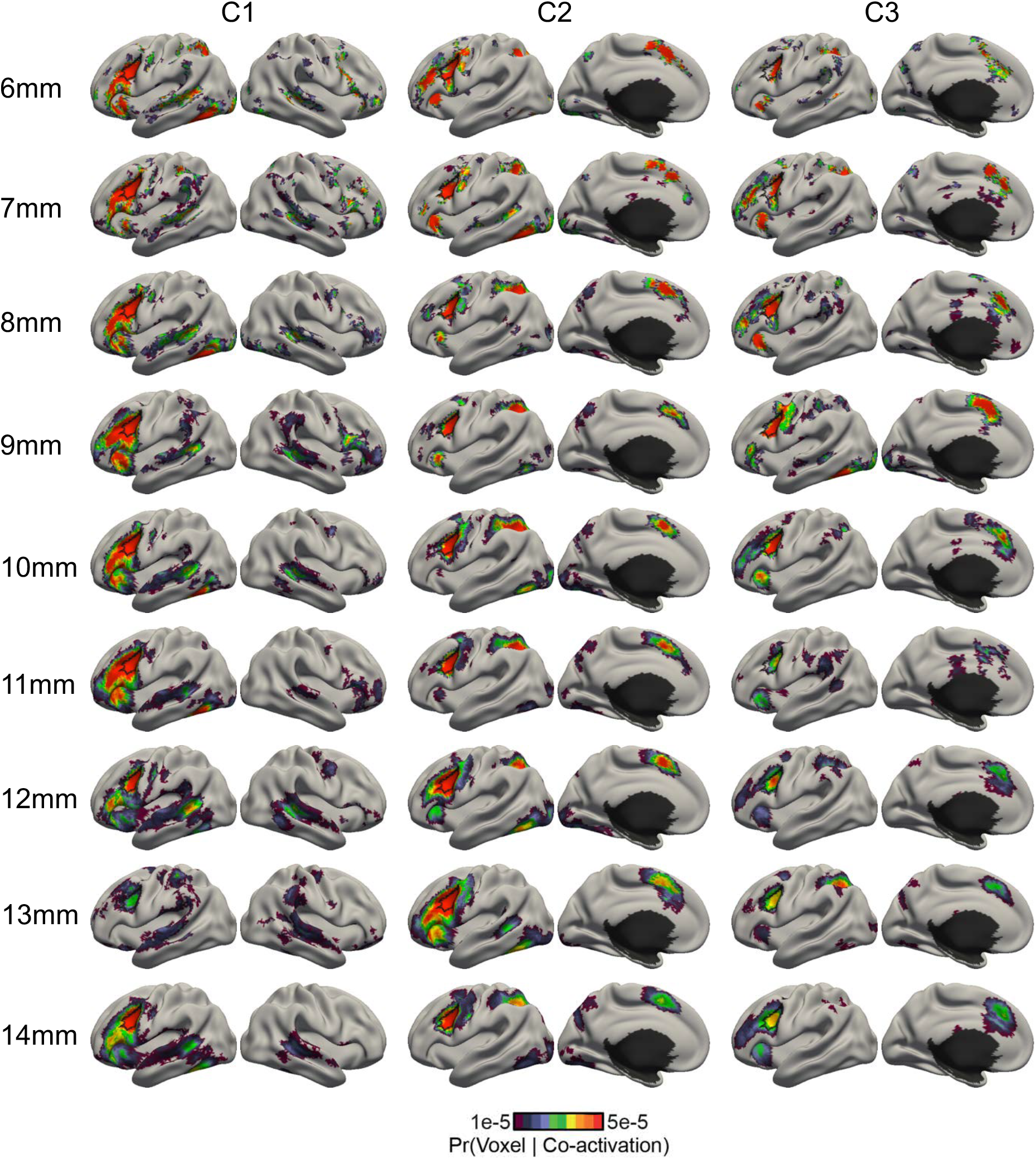
Co-activation patterns of the left inferior frontal junction estimated by the author-topic model across different radii of smoothing kernel applied to the input data. The smoothing radii ranged from 6mm to 14mm. Co-activation patterns estimated with different smoothing kernel radii are similar to the co-activation patterns estimated with a 10mm-radius smoothing kernel. The average Pearsons’ correlation coefficient of the co-activation patterns estimated with different smoothing kernel radii against those estimated with a 10mm-radius ranges from 0.54 (6mm) to 0.85 (14mm). The co-activation patterns estimated with a 10mm-radius smoothing kernel were shown in all results and analyses.

**Figure S7.**
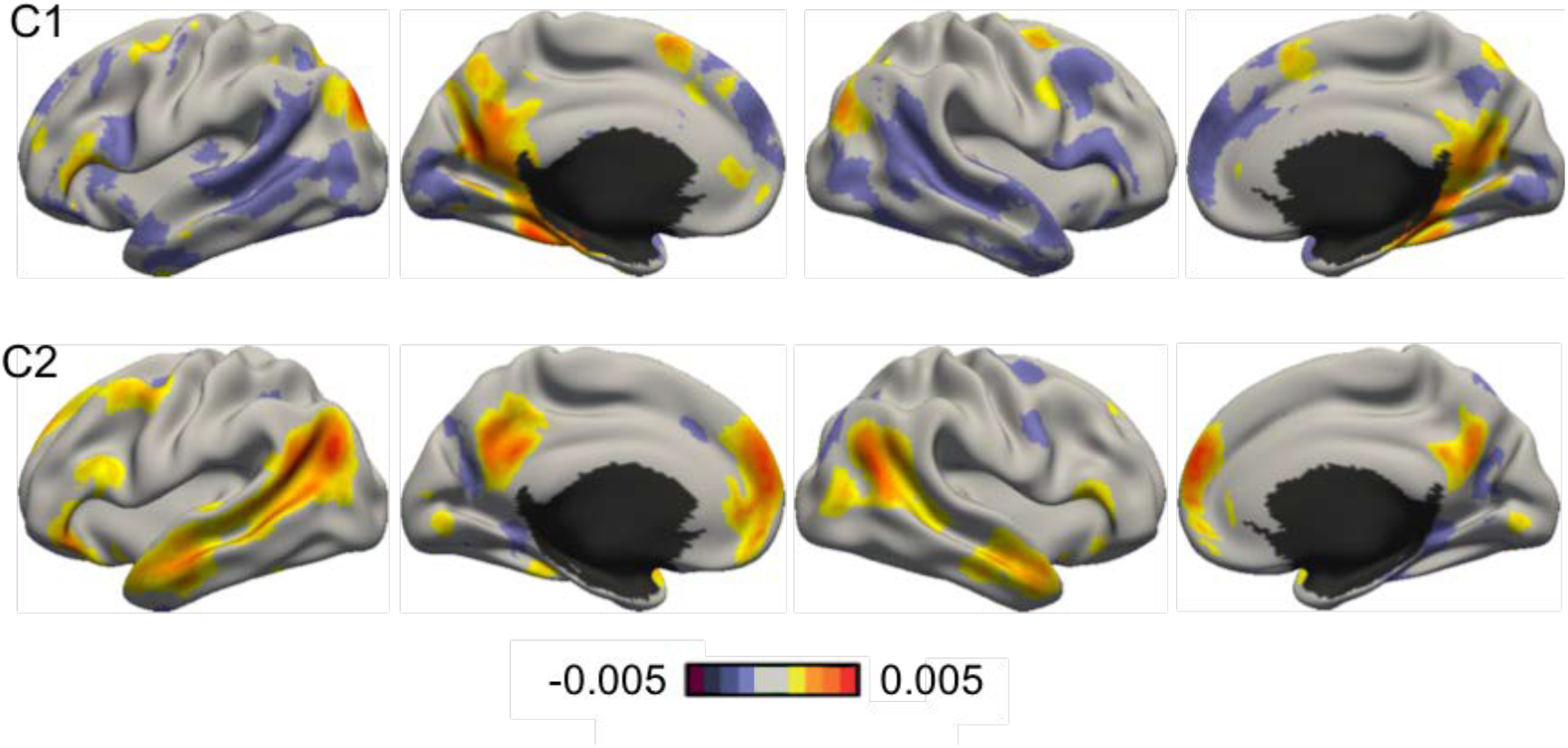
2-component ICA decomposition of self-generated thought dataset. Each independent component has both positive (yellow and red) and negative values (purple and maroon). The negative values in the independent components are not directly interpretable since the input data consisted of only “activation”, so it does not make sense to have “de-activation” in the component estimates.

**Figure S8.**
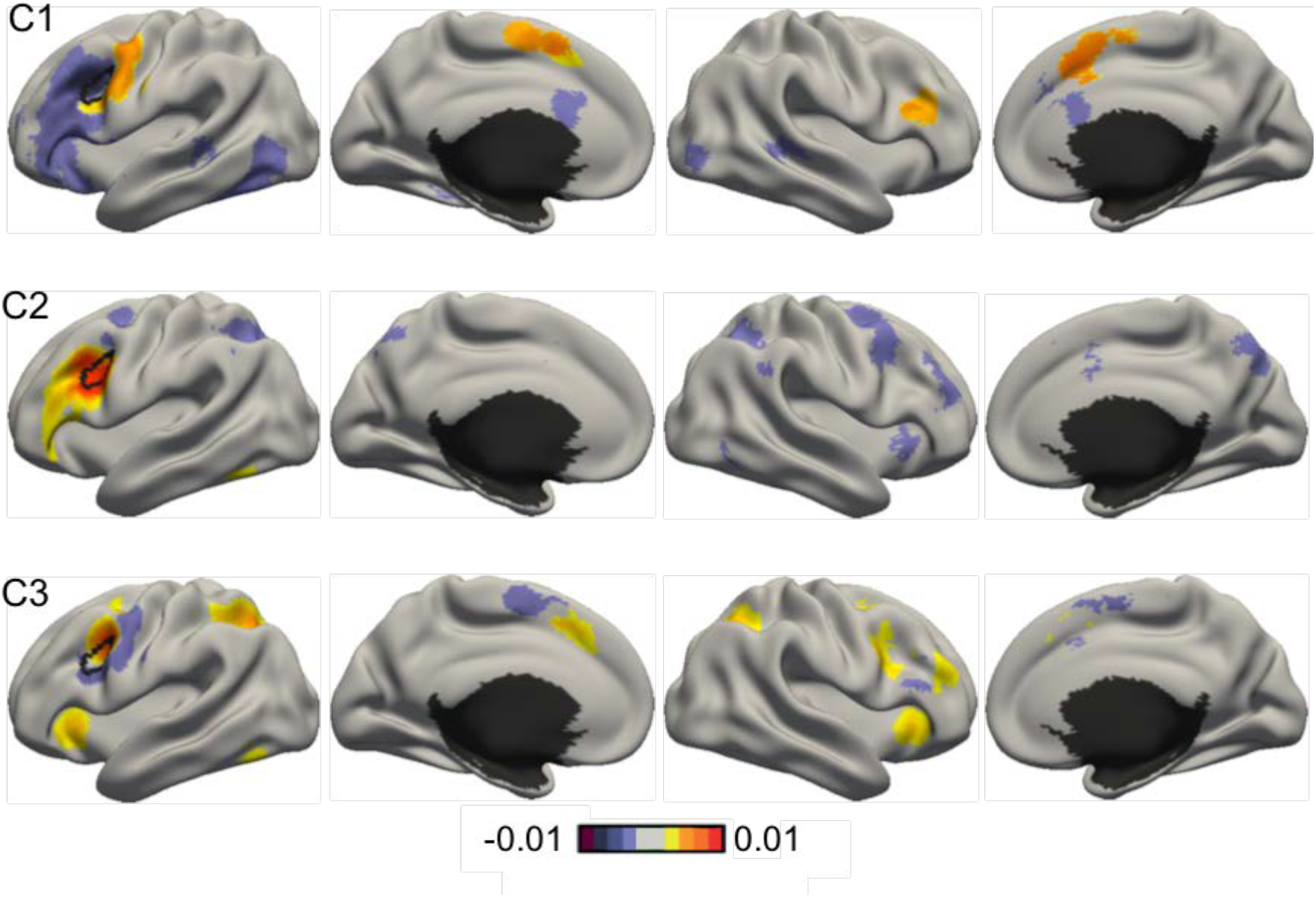
3-component ICA decomposition of the left inferior frontal junction (IFJ) activation data. Each independent component has both positive (yellow and red) and negative values (purple and maroon). The negative values in independent components are not directly interpretable, since the input data consisted of only “activation”, so it does not make sense to have “de-activation” in the component estimates. Component C1 does not engage most of the left IFJ seed region while component C3 has both positive and negative values within the seed region, suggesting that the ICA components might be fractionating the seed region, rather than discovering multiple co-activation patterns.

### 2. Supplemental Methods

#### 2.1 (S1) Mathematical Model

**Figure S9.**
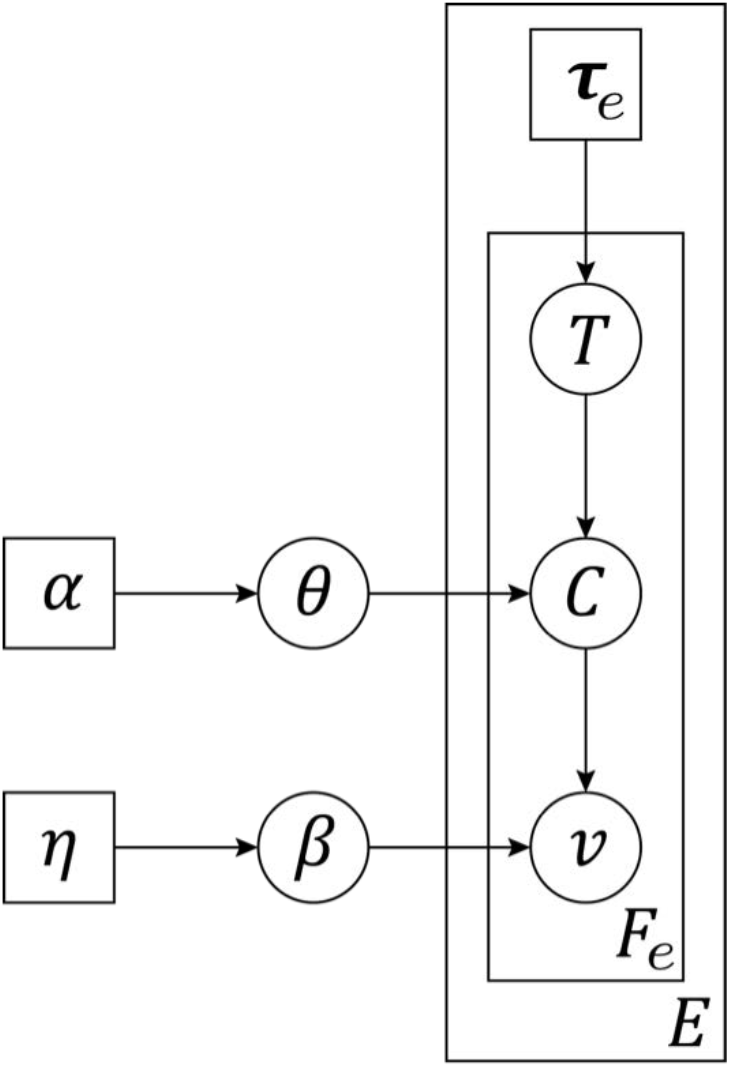
Formal graphical representation of the author-topic model for coordinate-based meta-analysis (Yeo et al. 2015). The circles represent random variables, while the squares represent non-random parameters. The edges represent statistical dependencies. There are *E* experiments. The *e*-th experiment utilizes a set of behavioral tasks **τ**_*e*_ and reports *F*_*e*_ number of activated voxels. The *f*-th activated voxel has an observed location *v*, and associated with a latent (unobserved) component *C* and a latent (unobserved) task *T* ∈ **τ**_*e*_. The variables at the corners of the rectangles (plates) indicate the number of times the variables within the rectangles were replicated. Therefore **τ**_*e*_ was replicated *E* times, once for each experiment. For the *e*-th experiment, the variables *v, C* and *T* were replicated *F*_*e*_ times, once for each activated voxel. *θ* denotes *P* (*component* | *task*) and *β* denotes *P r*(*voxel* | *component*). Thus *θ* and *β* are matrices, where each row is a categorical distribution summing to one. *α* and *η* are hyperparameters parameterizing the Dirichlet priors on *θ* and *β* respectively.

Figure S9 shows the formal graphical representation of the author-topic model for coordinate-based meta-analysis. The model assumes that there are *E* experiments. The *e*-th experiment is associated with a set of tasks **τ**_*e*_ and an unordered set *v*_*e*_ of *F*_*e*_ activated voxels. The location of the *f*-th activated voxel is denoted as *v*_*ef*_, corresponding to one of *V* = 284100 possible locations in MNI152 2mm space (Lancaster et al., 2007). The collection of activated voxels across all *E* experiments is denoted as *v*= {*v*_*e*_}. The collection of tasks across all *E* experiments is denoted as **τ**= {**τ**_*e*_}. Thus, {*v*, **τ**} are the input data for the meta-analysis.

We assume that there are *K* cognitive components and *M* unique tasks in the dataset. For example, *M* = 83 in Yeo et al. (2015). Each task has a certain (unknown) probability of recruiting a component Pr(component | task). The collection of all probabilities Pr(component | task) is denoted by a *M* × *K* matrix *θ*. The *t*-th row and *k*-th column of θ corresponds to the probability of the *t*-th task recruiting the *k*-th component. Each component has a certain (unknown) probability of activating a voxel pr(voxel | component). The collection of all probabilities Pr(voxel | component) is denoted by a *K* × *V* matrix *β*. The *k*-th row and *v*-th column of *β* corresponds to the probability of the *k* -th component activating the *v* -th MNI152 voxel. Symmetric Dirichlet priors with hyperparameter *α* are assumed on θ, and hyperparameter *η* on β.

We assume that the activated voxels of an experiment are independent and identically distributed (conditioned on knowing *θ* and *β*). To generate the *f*-th activated voxel *v*_*ef*_ in the *e*-th experiment, a task *T*_*ef*_ is sampled uniformly from the set of tasks **τ**_*e*_ utilized by the experiment. Given task *T*_*ef*_, a component *C*_*ef*_ is sampled based on the probability that the task would recruit a component (corresponding to the *T*_*ef*_-th row of the *θ* matrix). Given component *C*_*ef*_, the activation location *v*_*ef*_ is sampled based on the probability that the component would activate a voxel (corresponding to the *C*_*ef*_-th row of the *β* matrix). *T*_*ef*_ and *C*_*ef*_ are known as latent variables because they are not directly observed in the input data. We denote *T*= {*T*_*ef*_}, *C* = {*C*_*ef*_} as the collection of latent tasks and components across all experiments and activated voxels.

Given the number of cognitive components *K*, the fixed hyperparameters *α* and *η*, and the activated voxels and behavioral task categories {*v*, **τ**} of all experiments, the parameters Pr(component | task) *θ*, and Pr(voxel | component) *β* can be estimated using different algorithms. Gibbs sampling was proposed in the original author-topic paper (Rosen-Zvi et al. 2010). We also proposed a faster expectation maximization (EM) algorithm that was highly efficient on large amount of data (Yeo et al. 2015). In the present work, we used collapsed variational Bayes (CVB) inference (Ngo et al. 2016), which is less sensitive to choice of hyperparameters compared to the EM algorithm.

#### 2.2 (S2) Collapsed Variational Bayes (CVB) Inference

The CVB algorithm for the latent dirichlet allocation model (Blei et al., 2006) was introduced by Teh et al. (2006). We subsequently extended the CVB algorithm to the author-topic model (Ngo et al. 2016). Here we provide the derivation of the algorithm in detail.

We start by following the standard variational Bayesian inference procedure (Beal, 2003) of constructing the lower bound of the log data likelihood:

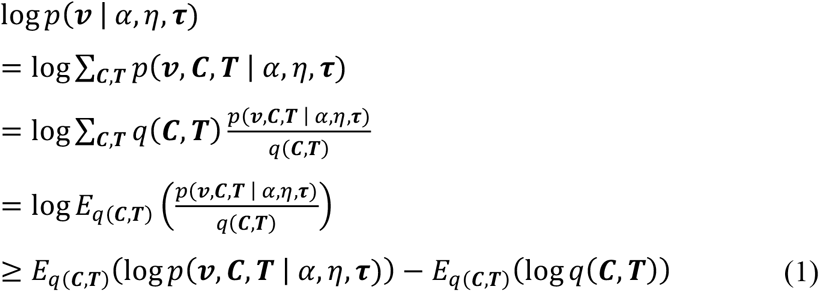

where q(***C, T***) can be any probability distribution and the inequality (Eq. (1)) utilizes the Jensen inequality (Beal, 2003). We can indirectly maximize the data likelihood (***v*** | *α, η*, **τ**) by finding the variational distribution q(***C, T***) that maximizes the lower bound (Eq. (1)). The equality in Eq. (1) occurs when q(***C, T***) = *p*(***C, T*** | ***v***, *α, η*, **τ**), i.e., when the variational distribution is equal to the true posterior distribution. However, computing the true posterior distribution of the latent variables is intractable because of dependencies among the variables constituting ***C*** and ***T***. Instead, the posterior of the latent variables (***C, T***) is approximated to be factorizable (Teh et al., 2006):

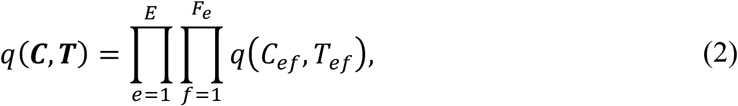

where q(***C, T***) is a categorical distribution with parameters *φ*:

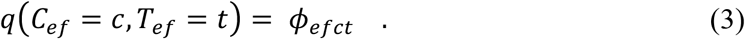

By plugging Eq. (3) into Eq. (1), we get

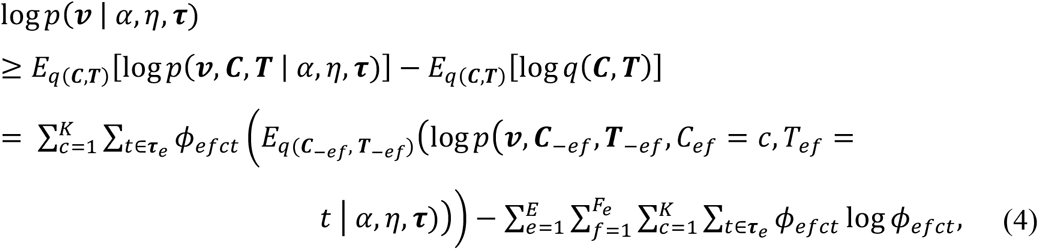

where the subscript “–*ef*” indicates the exclusion of corresponding variables *C*_*ef*_ and *T*_*ef*_. Maximizing the lower bound (Eq. (4)) by differentiating with respect to *φ* and using the constraint that 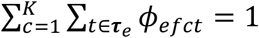, we get the update equation

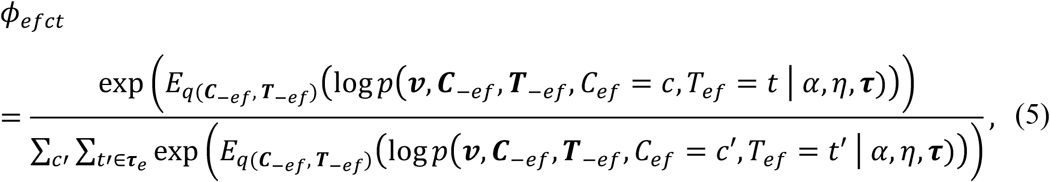

The CVB algorithm involves iterating Eq. (5) till convergence. The remaining derivations concern the evaluation of Eq. (5). We first apply the conditional independence assumptions of the author-topic model to simplify the joint probability distribution in Eq. (5):

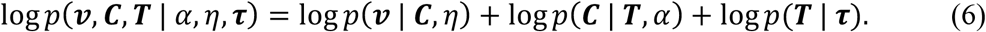

By exploiting the properties of Dirichlet-multinomial compound distribution (Teh et al., 2006), the first term on the right hand side of Eq. (6) is given by

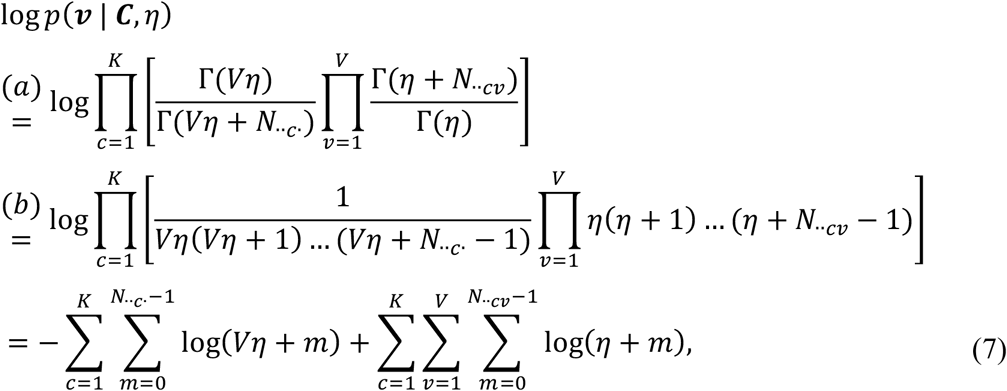

where equality (7a) arises from the definition of the Dirichlet-multinomial compound distribution and Γ(·) is the Gamma function. *N*_*etcv*_ is the number of activation foci in experiment *e* generated by task *t*, cognitive component *c*, and located at brain location *v*. The dot ‘·’ indicates that the corresponding variable is summed out. For example, *N* _*c*_ is the number of activation foci generated by component *c* across all experiments. Equation (7b) arises from the identity Γ(*z* + 1) = *z*Γ(*z*) for *z* > 0. Using the same procedure, the second term on the right hand side of Eq. (6) can be written as

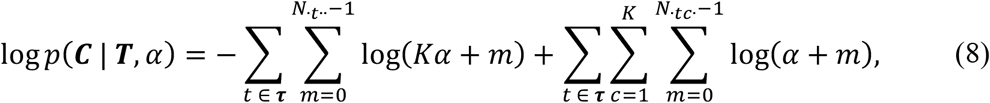

Substituting Eq. (7) and Eq. (8) back into Eq. (6), we get

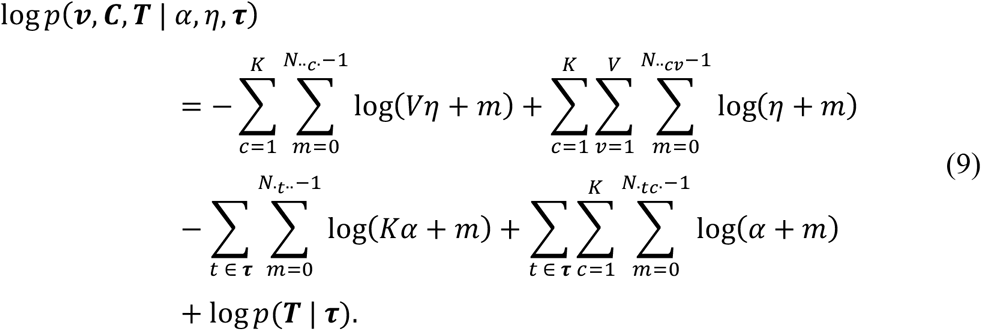

We are now ready to substitute Eq. (9) back into the update Eq. (5). The last term (log (***T*** | **τ**)) of Eq. (9) exists in both the numerator and denominator of Eq. (5) and thus cancels out. The remaining terms in Eq. (9) can be similarly simplified as follows. For example, the first term of Eq. (9) can be written as

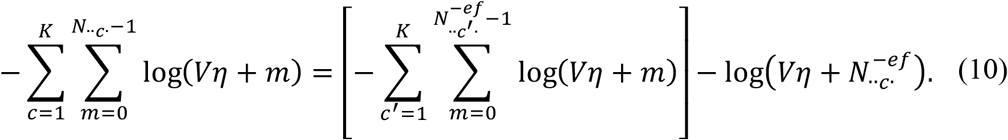

Therefore when Eq. (9) is substituted back into Eq. (5), the first term of Eq. (10) would be present in both the numerator and denominator of Eq. (5) and cancel out. Using the similar evaluation for the remaining terms, update Eq. (5) becomes

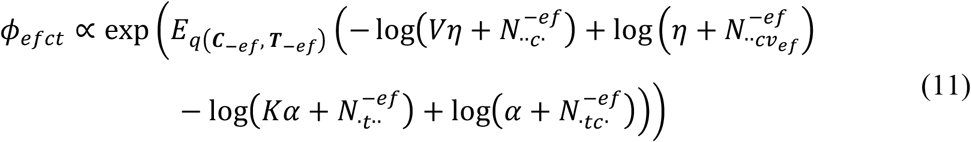

for *t* ∈ **τ**_*e*_ (otherwise *φ*_*efct*_ is zero), where the first log (·) term in Eq. (11) comes from the first term in Eq. (9), the second log (·) term in Eq. (11) comes from the second term in Eq. (9), and so on.

The log (·) terms in Eq. (10) can be approximated by a second-order Taylor’s series expansion about their means (Teh at al. 2006). Consider the Taylor’s series expansion of the log(*b* + *x*) function about a particular constant *a*:

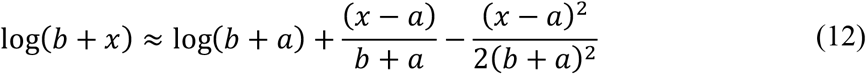

Applying the expansion in Eq. (12) with *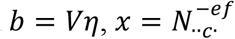,* and 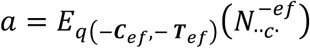, the first term in Eq. (11) can be approximated as

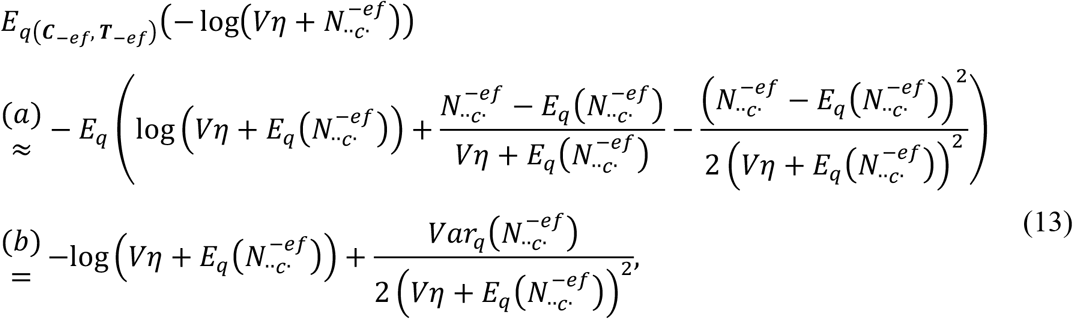

where the subscript (***C***_-*ef*_, ***T***_-*ef*_) in 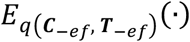 was omitted in Eq. (13a) to reduce clutter. Eq. (13b) was obtained because the expectation of a constant is itself. Therefore the first term corresponds to 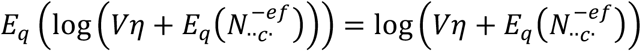. The second term evaluates to zero because 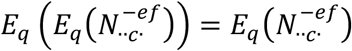.

Applying the same approximation for all log (·) terms in Eq. (11) and rearranging, the update equation for *φ* becomes

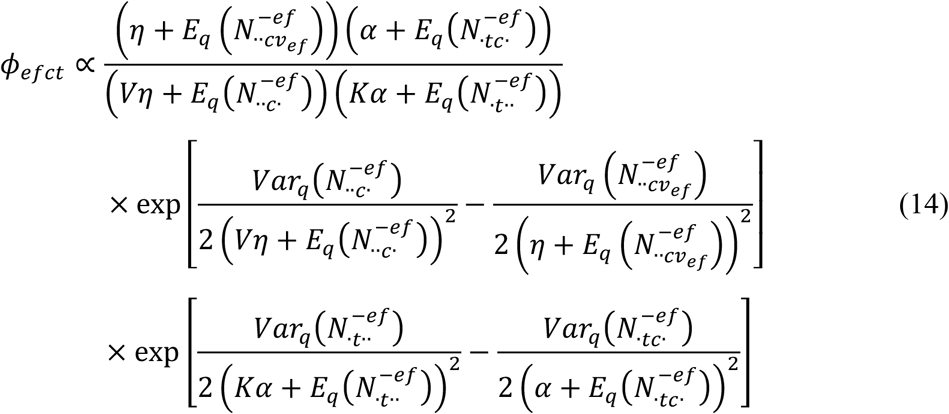

The mean and variance of the counts in Eq. (14) can be evaluated using the current estimate of *φ*. For example, 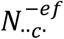 can be thought of as the number of heads obtained from tossing a coin independently for each focus of each experiment in the entire dataset (excluding the *f*-th focus of the *e*-th experiment), where the probability of getting a head for the *f*′-th activated voxel of the *e*′-th experiment is equal to 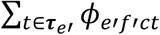 Thus, the expectation and variance of 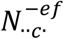 is given by

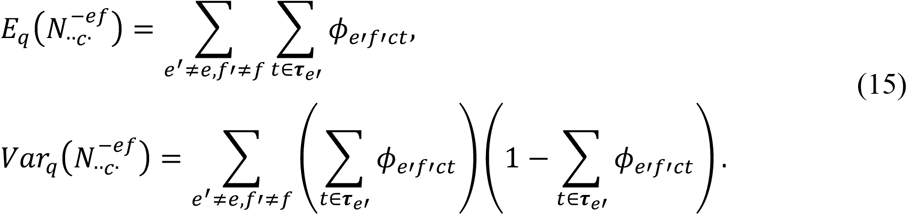

By using the same argument for the remaining terms of Eq. (14), we can evaluate the update equation for *φ*_*efct*_ given the current estimate of *φ*.

To summarize, the CVB algorithm proceeds by iterating Eq. (14) until convergence. Notice that under CVB inference, the posterior *φ* is estimated without using the point estimates of the model parameters *θ* and *β* (unlike the EM algorithm; see Supplemental Method S3). Given the final estimate of posterior distribution *φ*, the parameters *θ* and *β* can be estimated by the posterior means (Teh et al. 2006):

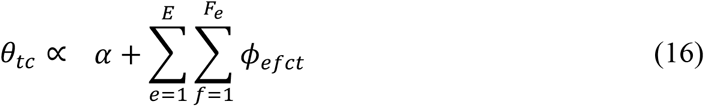

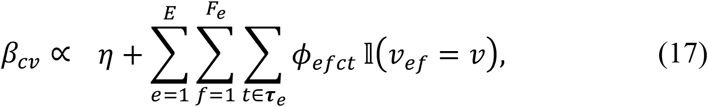

where 𝕝(*v*_*ef*_ = *v*) equals to one if the activation focus *v*_*ef*_ corresponds to location *v* in MNI152 space, and zero otherwise.

#### 2.3 (S3) Theoretical differences between CVB with Standard Variational Bayes (SVB) and EM algorithm

The CVB algorithm is theoretically better than standard variational Bayes (SVB) inference (Teh et al., 2006). As explained in the previous supplemental, CVB algorithm constructs a lower bound to the data log likelihood with respect to the latent variables (***C, T***). On the other hand, the SVB algorithm constructs a lower bound with respect to both latent variables (***C, T***) and model parameters (*θ, β*). Consequently, CVB provides a tighter lower bound to the data log likelihood:

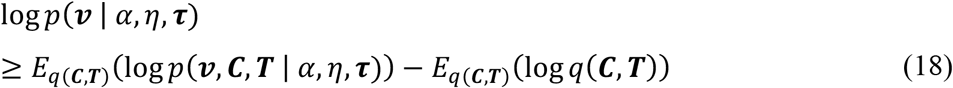

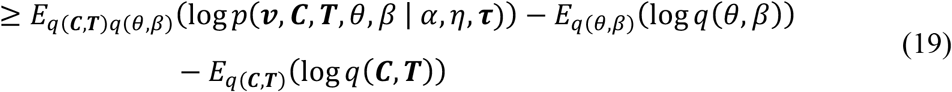

where the inequality (Eq. (18)) is the same as CVB Eq. (1) and the second inequality (Eq. (19)) corresponds to SVB.

One can also draw parallels between the CVB (Supplemental Method S2) and EM (Yeo et al., 2015) algorithms for the author-topic model. Both algorithms iterate between estimating the posterior distribution of the latent variables (***C, T***) and using the posterior distribution to update the model parameters estimates (*θ, β*). However, the EM algorithm uses *point* estimates of the model parameters to update the posterior distribution of (***C, T***). In contrast, the CVB algorithm avoids doing so (Eq. (14)) and might therefore produce better estimates of the parameters (Ngo et al. 2016).

In practice, we find the CVB algorithm to be less sensitive than the EM algorithm to the initialization of the hyperparameters *α* and *η*. This is not an issue for a big dataset (e.g., BrainMap; Yeo et al., 2015) because the data will overwhelm the priors. However, this issue is important for small datasets like those utilized in this work. For the CVB algorithm, the hyperparameters *α* and *η* were set to 100 and 0.01 respectively across all of our experiments. Perturbing *α* and *η* by two orders of magnitude did not significantly change the model parameters estimated by CVB algorithm, suggesting its robustness. This was not the case for the EM algorithm.

#### 2.4 (S4) Estimating Number of Components using Bayesian Information Criterion (BIC)

Bayesian Information Criterion (BIC) is commonly used for model selection in machine learning (Schwarz, 1978). BIC favors models that best fit the data, while also penalizing models with more parameters. The BIC for the author-topic model is given by:

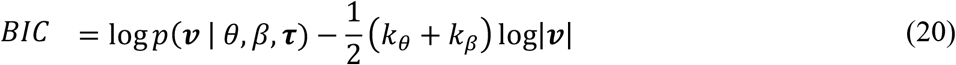

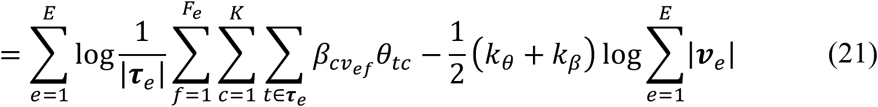

where the first term is the log likelihood of the activation foci *v* given the model parameters estimates *θ* and *β*, and the second term is the penalty based on the number of model parameters. |***v***_*e*_| and |**τ**_*e*_| are the number of foci and tasks employed in the *e*-th experiment. *k*_*θ*_ and *k*_*β*_ are the number of free model parameters. *k*_*θ*_ is the number of free parameters in the *M* × *K* matrix *θ*, which is equal to *M* × (*K* – 1) since each row sums to one. *k*_*β*_ is the approximated number of independent elements in the *K* × *V* matrix *β*. Each row of *β* can be interpreted as a spatially smoothed brain image (see Supplemental Method S5). Therefore we approximated the number of independent elements in each row of *β* by the number of resolution elements (resels) in the corresponding brain image (Worsley et al. 1992) using AFNI (Cox 1996).

Models with a higher number of components *K* fit the data better, resulting in a higher data log likelihood (first term of Eq. (20)). On the other hand, a higher *K* also increases the number of free parameters *k*_*θ*_ + *k*_*β*_ b, which results in a higher penalty (second term of Eq. (20)). A higher BIC values indicates a better model.

#### 2.5 (S5) Implementation Details

The model’s hyperparameters were set to be *α* = 100 and *η* = 0.01 across all experiments. Perturbing *α* and *η* by two orders of magnitude did not significantly change the results. The posterior distribution *φ* was randomly initialized. The CVB algorithm then updated the posterior distribution *φ* (Eq. (14)) until convergence. Given the estimate of *φ*, CVB algorithm then computed the model parameters *θ* and *β* (Eq. (16) and Eq. (17)). For a given number of components *K*, the procedure was repeated with 1000 random initializations resulting in 1000 estimates. The estimate resulting in the maximum lower bound of the data log likelihood (Eq. (1)) was taken as the final estimate.

We repeated the procedure with for different number of cognitive components *K*. BIC was computed for each value of *K* (Supplemental Method S4). Higher BIC implied better model parameters estimates. The model parameters with the highest BIC were presented in the Results and Discussion sections.

#### 2.6 (S6) Approximation of Pr(co-activation pattern | task)

For the co-activation analysis of IFJ, each experiment was treated as its own unique task. To help interpret the co-activation pattern in terms of BrainMap task categories (also known as paradigm classes), we estimated the probability of the *c*-th co-activation pattern being utilized by the *t*-th task posthoc:

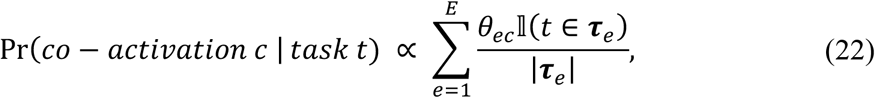

where *θ*_*ec*_ was the estimated probability that the *e*-th experiment would recruit the *c*-th co-activation pattern, **τ**_*e*_ was the set of tasks utilized in the *e*-th experiment, and 𝕝(*t* ∈ **τ**_*e*_) is an indicator function that was equal to 1 if the *t*-th task was one of the collection of tasks **τ**_*e*_ utilized by the *e*-th experiment and 0 otherwise. Eq. (22) can be interpreted as weighted average of *θ*_*e*i_ across all experiments utilizing task *t* with the weight being smaller if an experiment utilizes many tasks. For example, if the 3rd experiment utilized “n-back” and “Stroop” tasks, the probability contributed by this experiment to the computation of the probability of “n-back” recruiting co-activation pattern C1 (Pr(*co* – *activation C*1 | “*n* – *back*”)) would be the probability of the experiment recruiting co-activation pattern C1 (i.e., *θ*_31_), divided by the number of tasks, which is two.

#### 2.7 (S7) Self-generated thought studies

##### 2.7.1 Summary tables of self-generated thought studies

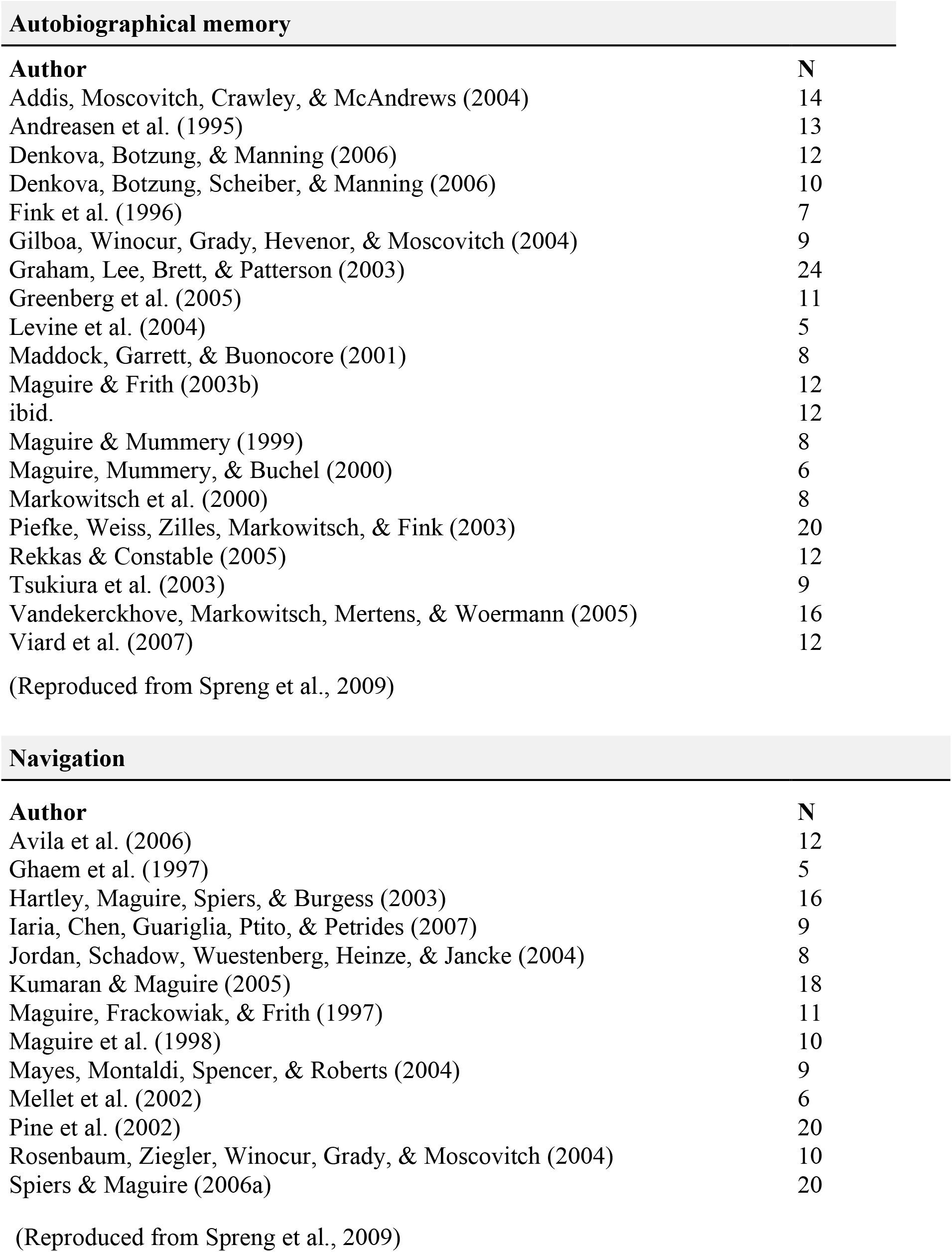

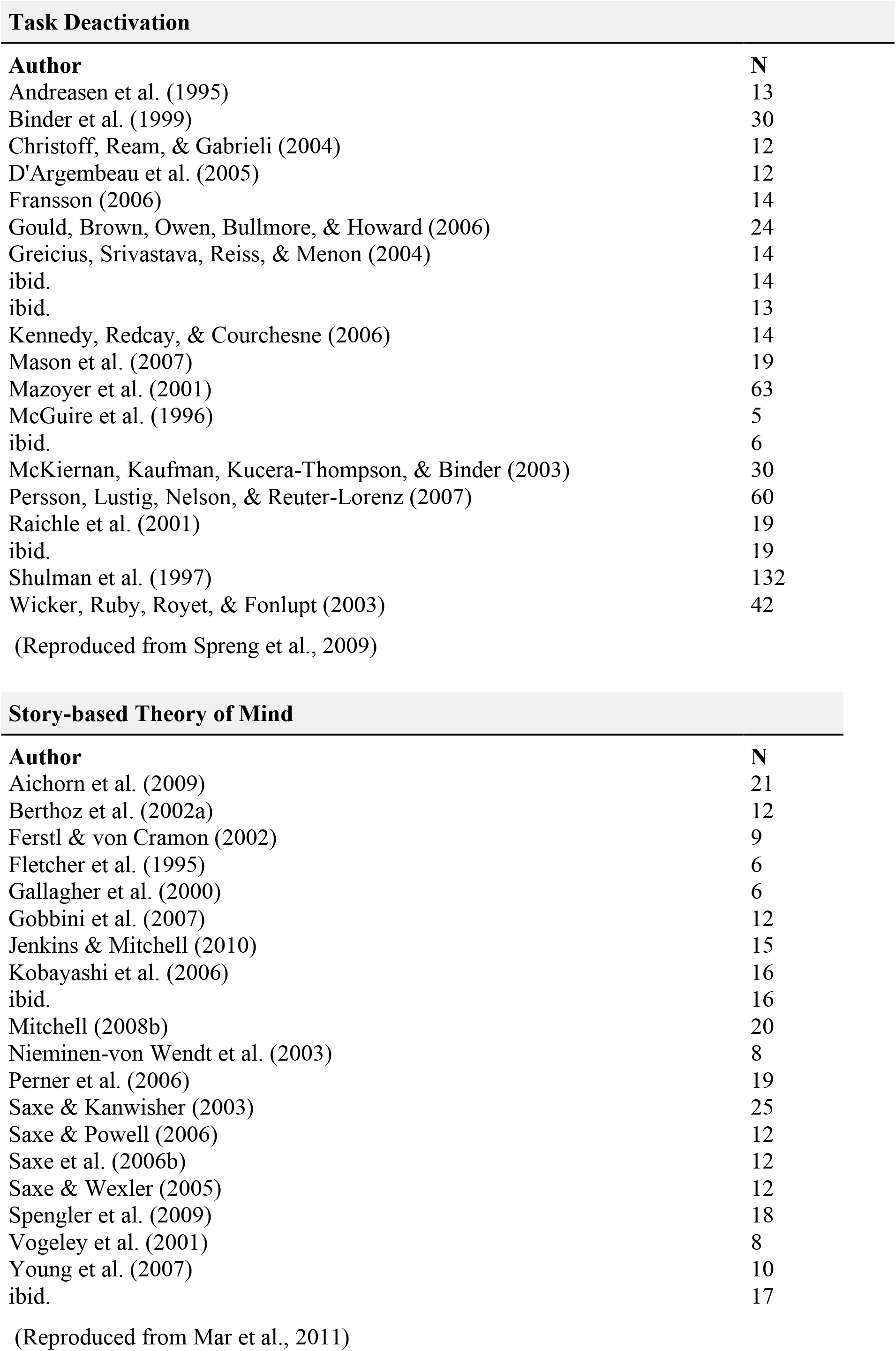

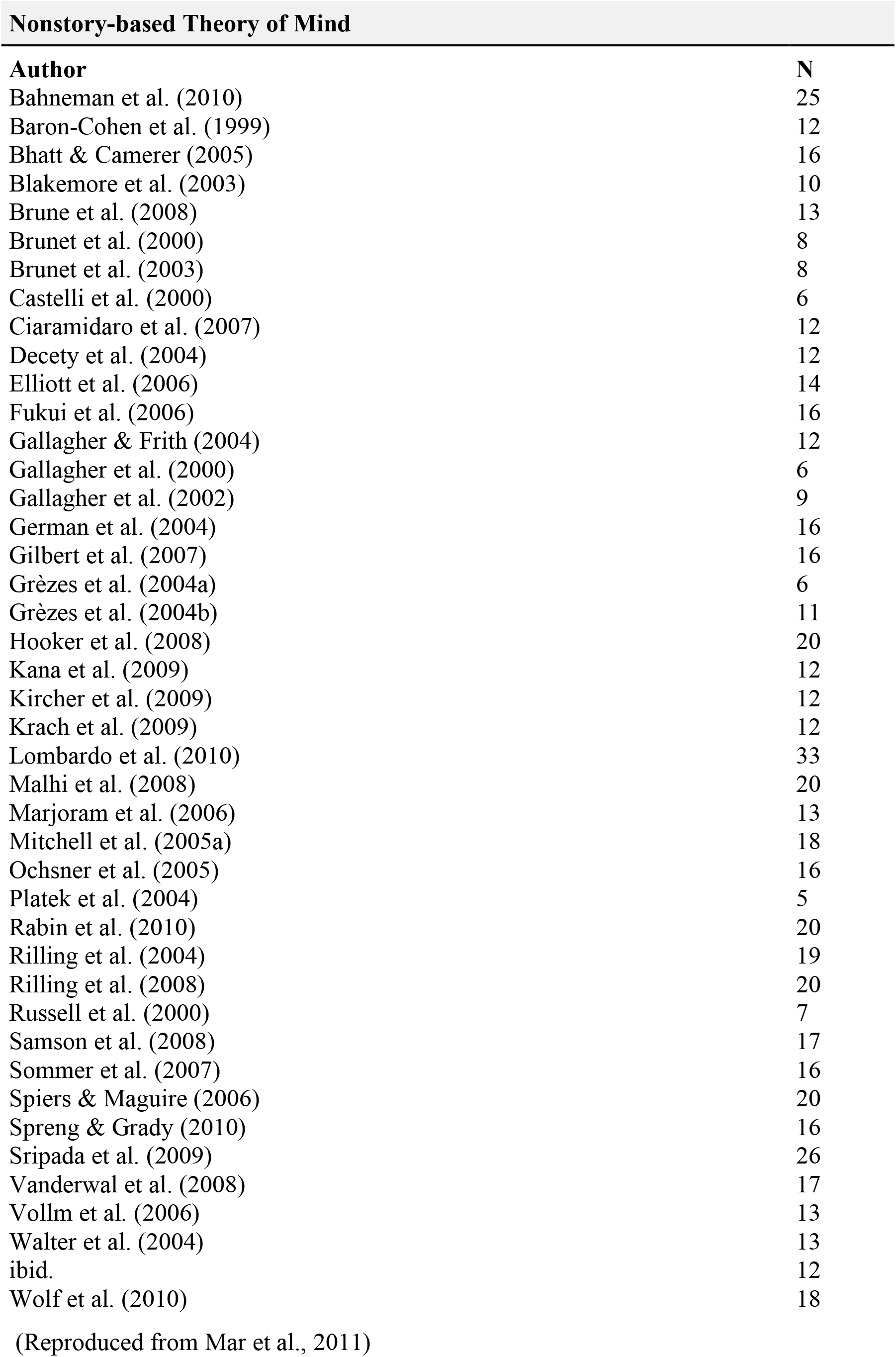

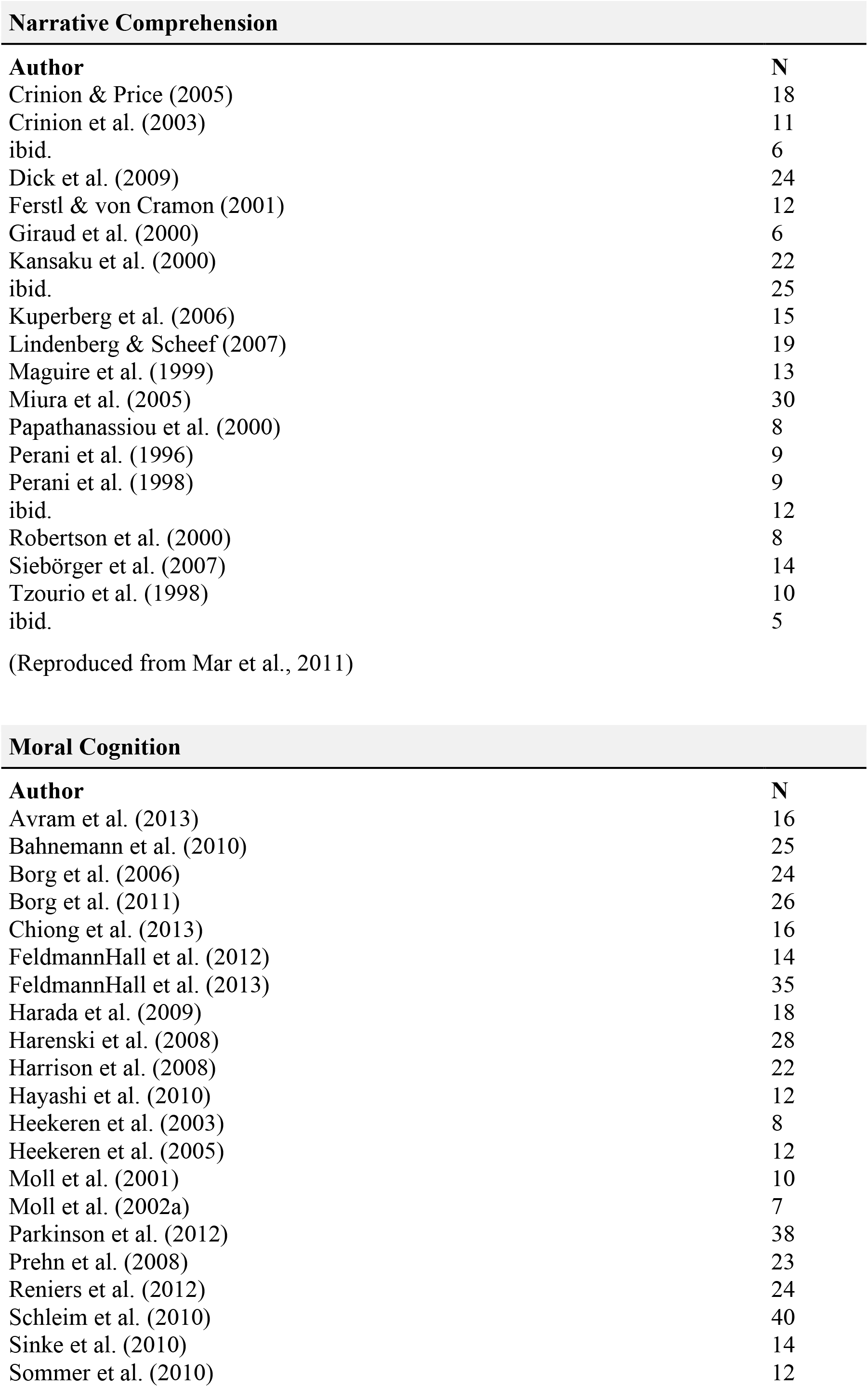

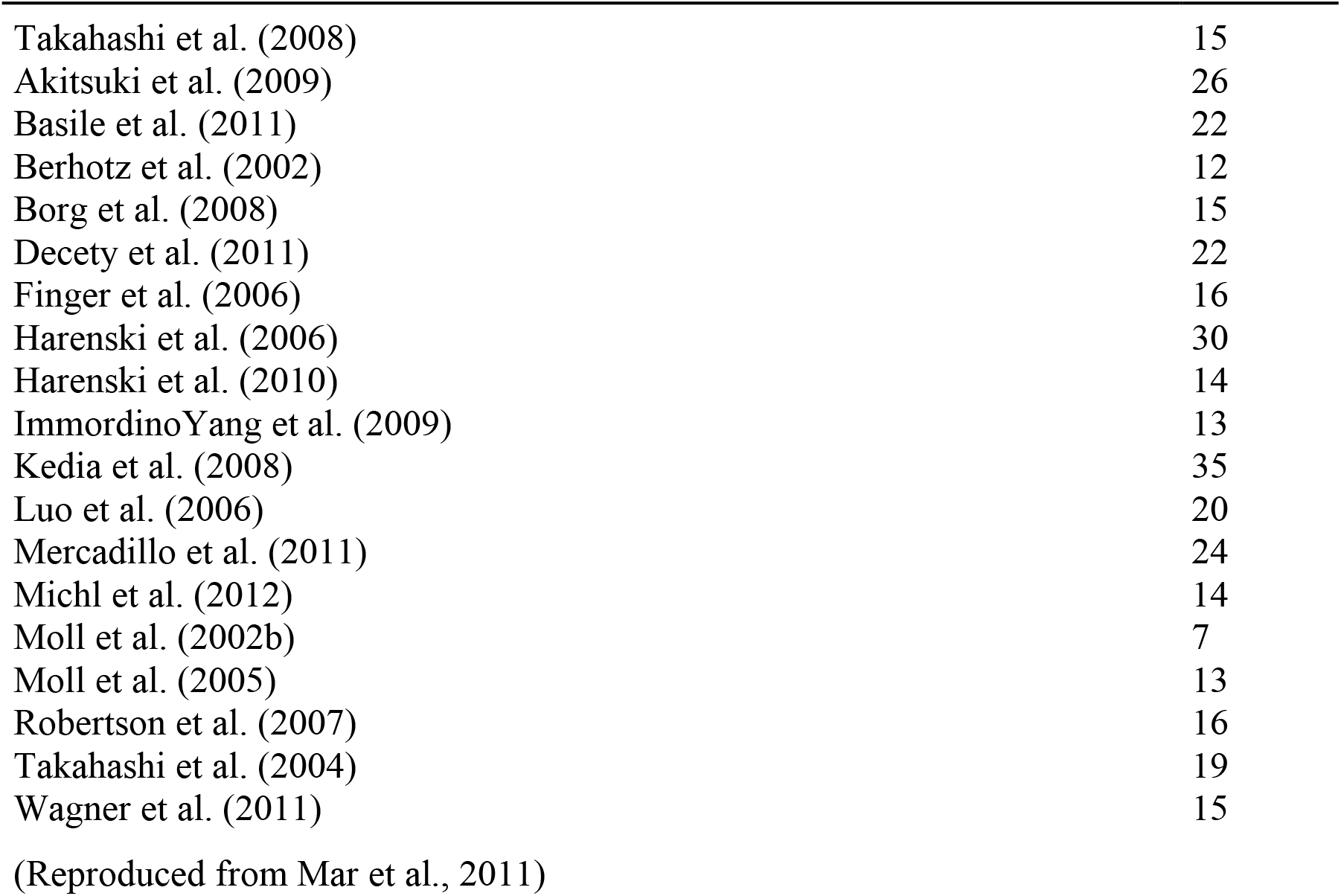

Autobiographical memory
| Author | N |
| --- | --- |
| Addis, Moscovitch, Crawley, & McAndrews (2004) | 14 |
| Andreasen et al. (1995) | 13 |
| Denkova, Botzung, & Manning (2006) | 12 |
| Denkova, Botzung, Scheiber, & Manning (2006) | 10 |
| Fink et al. (1996) | 7 |
| Gilboa, Winocur, Grady, Hevenor, & Moscovitch (2004) | 9 |
| Graham, Lee, Brett, & Patterson (2003) | 24 |
| Greenberg et al. (2005) | 11 |
| Levine et al. (2004) | 5 |
| Maddock, Garrett, & Buonocore (2001) | 8 |
| Maguire & Frith (2003b) | 12 |
| ibid. | 12 |
| Maguire & Mummery (1999) | 8 |
| Maguire, Mummery, & Buchel (2000) | 6 |
| Markowitsch et al. (2000) | 8 |
| Piefke, Weiss, Zilles, Markowitsch, & Fink (2003) | 20 |
| Rekkas & Constable (2005) | 12 |
| Tsukiura et al. (2003) | 9 |
| Vandekerckhove, Markowitsch, Mertens, & Woermann (2005) | 16 |
| Viard et al. (2007) | 12 |

Navigation
| Author | N |
| --- | --- |
| Avila et al. (2006) | 12 |
| Ghaem et al. (1997) | 5 |
| Hartley, Maguire, Spiers, & Burgess (2003) | 16 |
| Iaria, Chen, Guariglia, Ptito, & Petrides (2007) | 9 |
| Jordan, Schadow, Wuestenberg, Heinze, & Jancke (2004) | 8 |
| Kumaran & Maguire (2005) | 18 |
| Maguire, Frackowiak, & Frith (1997) | 11 |
| Maguire et al. (1998) | 10 |
| Mayes, Montaldi, Spencer, & Roberts (2004) | 9 |
| Mellet et al. (2002) | 6 |
| Pine et al. (2002) | 20 |
| Rosenbaum, Ziegler, Winocur, Grady, & Moscovitch (2004) | 10 |
| Spiers & Maguire (2006a) | 20 |

**Task Deactivation**
| **Author** | **N** |
| --- | --- |
| Andreasen et al. (1995) | 13 |
| Binder et al. (1999) | 30 |
| Christoff, Ream, & Gabrieli (2004) | 12 |
| D'Argembeau et al. (2005) | 12 |
| Fransson (2006) | 14 |
| Gould, Brown, Owen, Bullmore, & Howard (2006) | 24 |
| Greicius, Srivastava, Reiss, & Menon (2004) | 14 |
| ibid. | 14 |
| ibid. | 13 |
| Kennedy, Redcay, & Courchesne (2006) | 14 |
| Mason et al. (2007) | 19 |
| Mazoyer et al. (2001) | 63 |
| McGuire et al. (1996) | 5 |
| ibid. | 6 |
| McKiernan, Kaufman, Kucera-Thompson, & Binder (2003) | 30 |
| Persson, Lustig, Nelson, & Reuter-Lorenz (2007) | 60 |
| Raichle et al. (2001) | 19 |
| ibid. | 19 |
| Shulman et al. (1997) | 132 |
| Wicker, Ruby, Royet, & Fonlupt (2003) | 42 |
| (Reproduced from Spreng et al., 2009) |  |

**Story-based Theory of Mind**
| **Author** | **N** |
| --- | --- |
| Aichorn et al. (2009) | 21 |
| Berthoz et al. (2002a) | 12 |
| Ferstl & von Cramon (2002) | 9 |
| Fletcher et al. (1995) | 6 |
| Gallagher et al. (2000) | 6 |
| Gobbini et al. (2007) | 12 |
| Jenkins & Mitchell (2010) | 15 |
| Kobayashi et al. (2006) | 16 |
| ibid. | 16 |
| Mitchell (2008b) | 20 |
| Nieminen-von Wendt et al. (2003) | 8 |
| Perner et al. (2006) | 19 |
| Saxe & Kanwisher (2003) | 25 |
| Saxe & Powell (2006) | 12 |
| Saxe et al. (2006b) | 12 |
| Saxe & Wexler (2005) | 12 |
| Spengler et al. (2009) | 18 |
| Vogeley et al. (2001) | 8 |
| Young et al. (2007) | 10 |
| ibid. | 17 |

**Nonstory-based Theory of Mind**
| **Author** | **N** |
| --- | --- |
| Bahneman et al. (2010) | 25 |
| Baron-Cohen et al. (1999) | 12 |
| Bhatt & Camerer (2005) | 16 |
| Blakemore et al. (2003) | 10 |
| Brune et al. (2008) | 13 |
| Brunet et al. (2000) | 8 |
| Brunet et al. (2003) | 8 |
| Castelli et al. (2000) | 6 |
| Ciaramidaro et al. (2007) | 12 |
| Decety et al. (2004) | 12 |
| Elliott et al. (2006) | 14 |
| Fukui et al. (2006) | 16 |
| Gallagher & Frith (2004) | 12 |
| Gallagher et al. (2000) | 6 |
| Gallagher et al. (2002) | 9 |
| German et al. (2004) | 16 |
| Gilbert et al. (2007) | 16 |
| Grèzes et al. (2004a) | 6 |
| Grèzes et al. (2004b) | 11 |
| Hooker et al. (2008) | 20 |
| Kana et al. (2009) | 12 |
| Kircher et al. (2009) | 12 |
| Krach et al. (2009) | 12 |
| Lombardo et al. (2010) | 33 |
| Malhi et al. (2008) | 20 |
| Marjoram et al. (2006) | 13 |
| Mitchell et al. (2005a) | 18 |
| Ochsner et al. (2005) | 16 |
| Platek et al. (2004) | 5 |
| Rabin et al. (2010) | 20 |
| Rilling et al. (2004) | 19 |
| Rilling et al. (2008) | 20 |
| Russell et al. (2000) | 7 |
| Samson et al. (2008) | 17 |
| Sommer et al. (2007) | 16 |
| Spiers & Maguire (2006) | 20 |
| Spreng & Grady (2010) | 16 |
| Sripada et al. (2009) | 26 |
| Vanderwal et al. (2008) | 17 |
| Vollm et al. (2006) | 13 |
| Walter et al. (2004) | 13 |
| ibid. | 12 |
| Wolf et al. (2010) | 18 |

Narrative Comprehension
| Author | N |
| --- | --- |
| Crinion & Price (2005) | 18 |
| Crinion et al. (2003) | 11 |
| ibid. | 6 |
| Dick et al. (2009) | 24 |
| Ferstl & von Cramon (2001) | 12 |
| Giraud et al. (2000) | 6 |
| Kansaku et al. (2000) | 22 |
| ibid. | 25 |
| Kuperberg et al. (2006) | 15 |
| Lindenberg & Scheef (2007) | 19 |
| Maguire et al. (1999) | 13 |
| Miura et al. (2005) | 30 |
| Papathanassiou et al. (2000) | 8 |
| Perani et al. (1996) | 9 |
| Perani et al. (1998) | 9 |
| ibid. | 12 |
| Robertson et al. (2000) | 8 |
| Siebörger et al. (2007) | 14 |
| Tzourio et al. (1998) | 10 |
| ibid. | 5 |

Moral Cognition
| Author | N |
| --- | --- |
| Avram et al. (2013) | 16 |
| Bahnemann et al. (2010) | 25 |
| Borg et al. (2006) | 24 |
| Borg et al. (2011) | 26 |
| Chiong et al. (2013) | 16 |
| FeldmannHall et al. (2012) | 14 |
| FeldmannHall et al. (2013) | 35 |
| Harada et al. (2009) | 18 |
| Harenski et al. (2008) | 28 |
| Harrison et al. (2008) | 22 |
| Hayashi et al. (2010) | 12 |
| Heekeren et al. (2003) | 8 |
| Heekeren et al. (2005) | 12 |
| Moll et al. (2001) | 10 |
| Moll et al. (2002a) | 7 |
| Parkinson et al. (2012) | 38 |
| Prehn et al. (2008) | 23 |
| Reniers et al. (2012) | 24 |
| Schleim et al. (2010) | 40 |
| Sinke et al. (2010) | 14 |
| Sommer et al. (2010) | 12 |
| Takahashi et al. (2008) | 15 |
| Akitsuki et al. (2009) | 26 |
| Basile et al. (2011) | 22 |
| Berhotz et al. (2002) | 12 |
| Borg et al. (2008) | 15 |
| Decety et al. (2011) | 22 |
| Finger et al. (2006) | 16 |
| Harenski et al. (2006) | 30 |
| Harenski et al. (2010) | 14 |
| ImmordinoYang et al. (2009) | 13 |
| Kedia et al. (2008) | 35 |
| Luo et al. (2006) | 20 |
| Mercadillo et al. (2011) | 24 |
| Michl et al. (2012) | 14 |
| Moll et al. (2002b) | 7 |
| Moll et al. (2005) | 13 |
| Robertson et al. (2007) | 16 |
| Takahashi et al. (2004) | 19 |
| Wagner et al. (2011) | 15 |

(Reproduced from Mar et al., 2011)

## 2.8 (S8) Inferior Frontal Junction (IFJ) studies

### 2.8.1 Summary table of IFJ studies

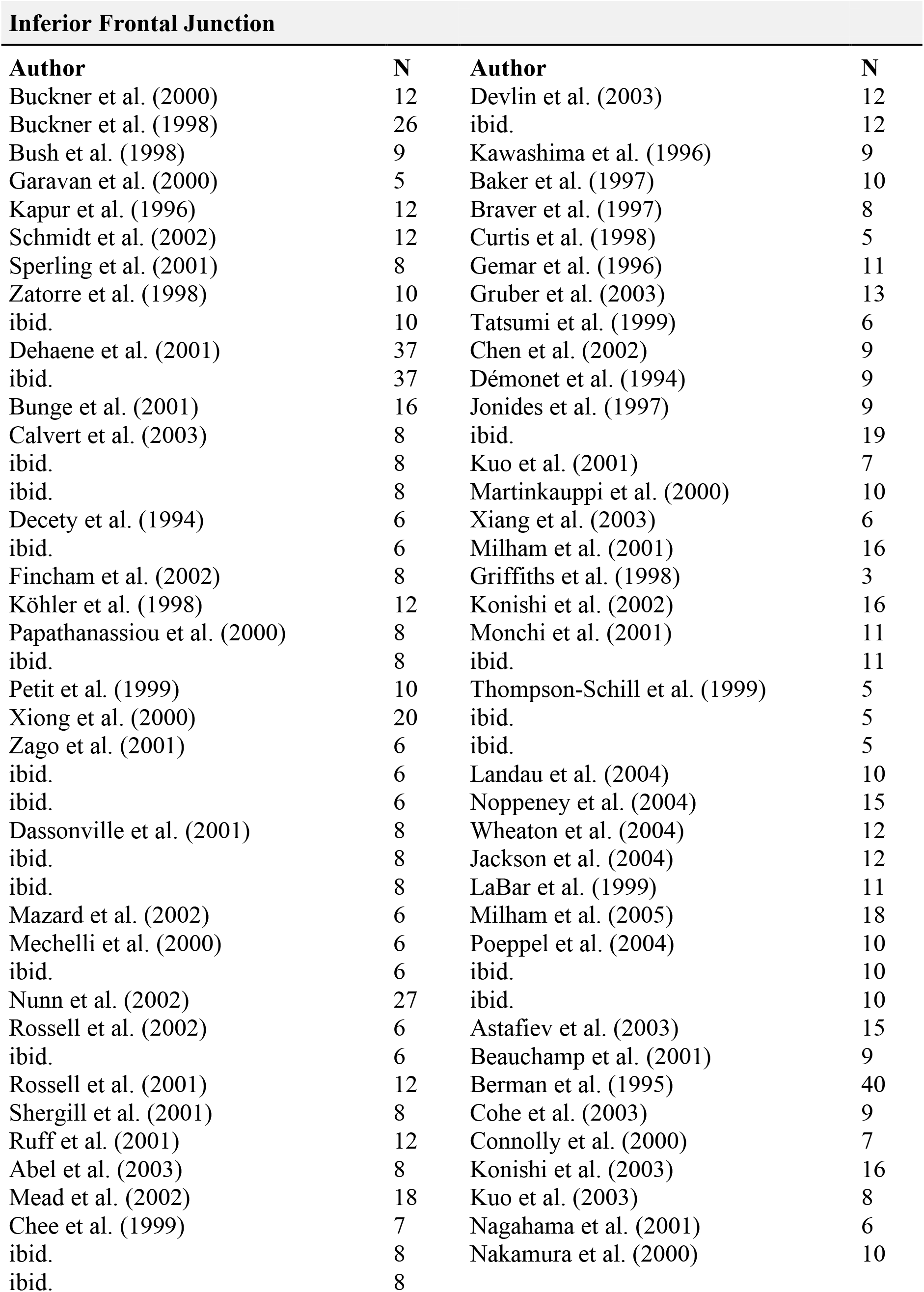

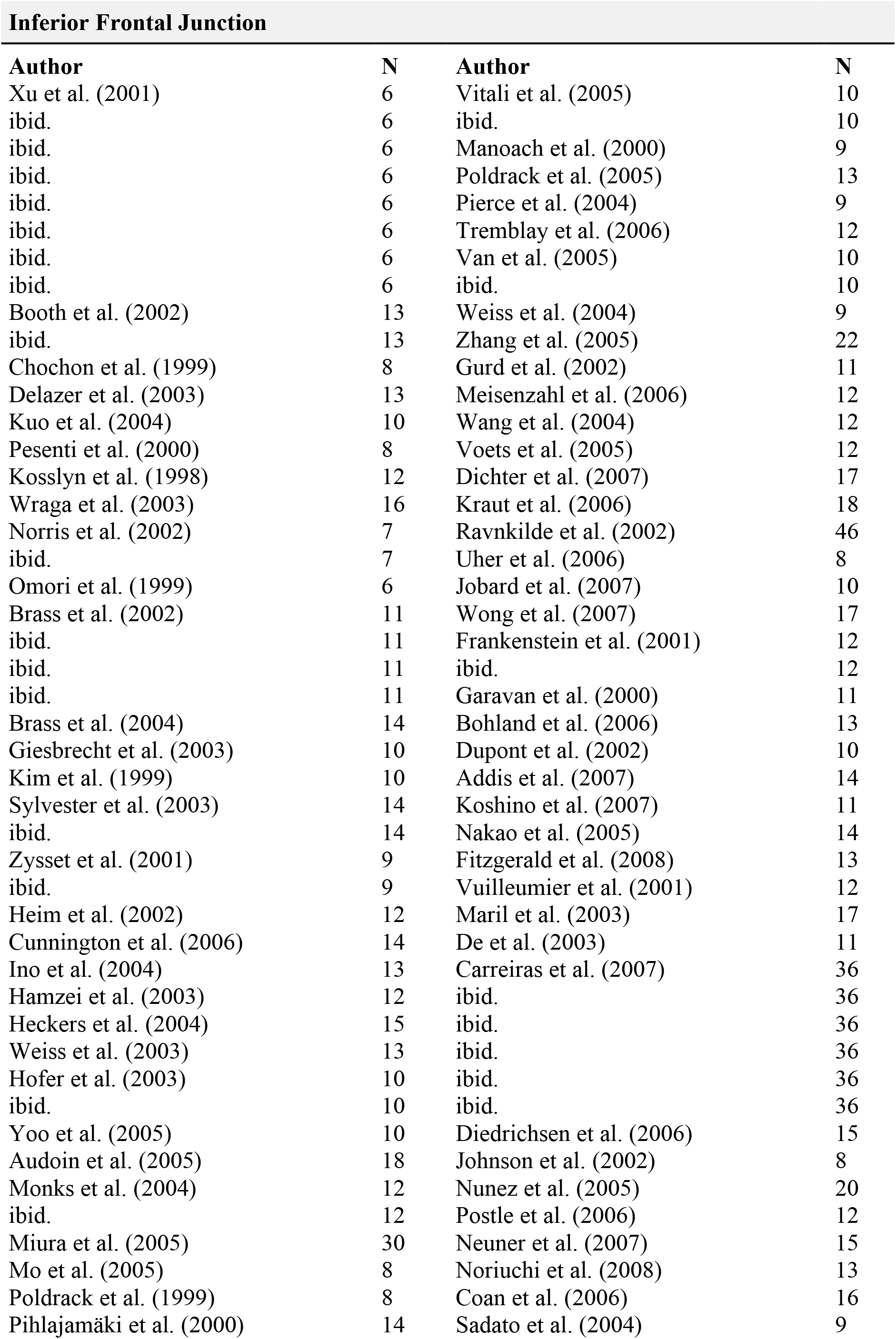

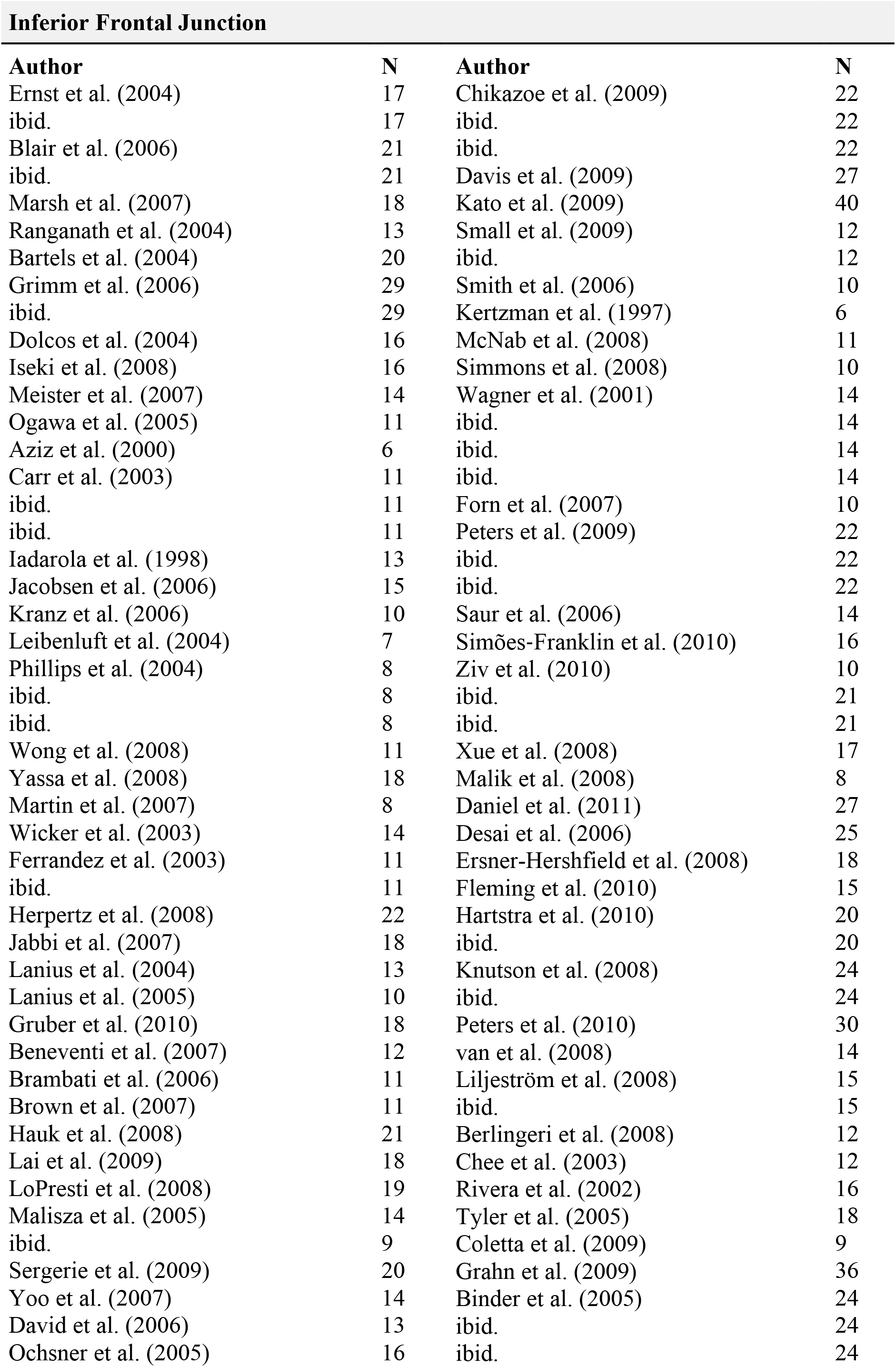

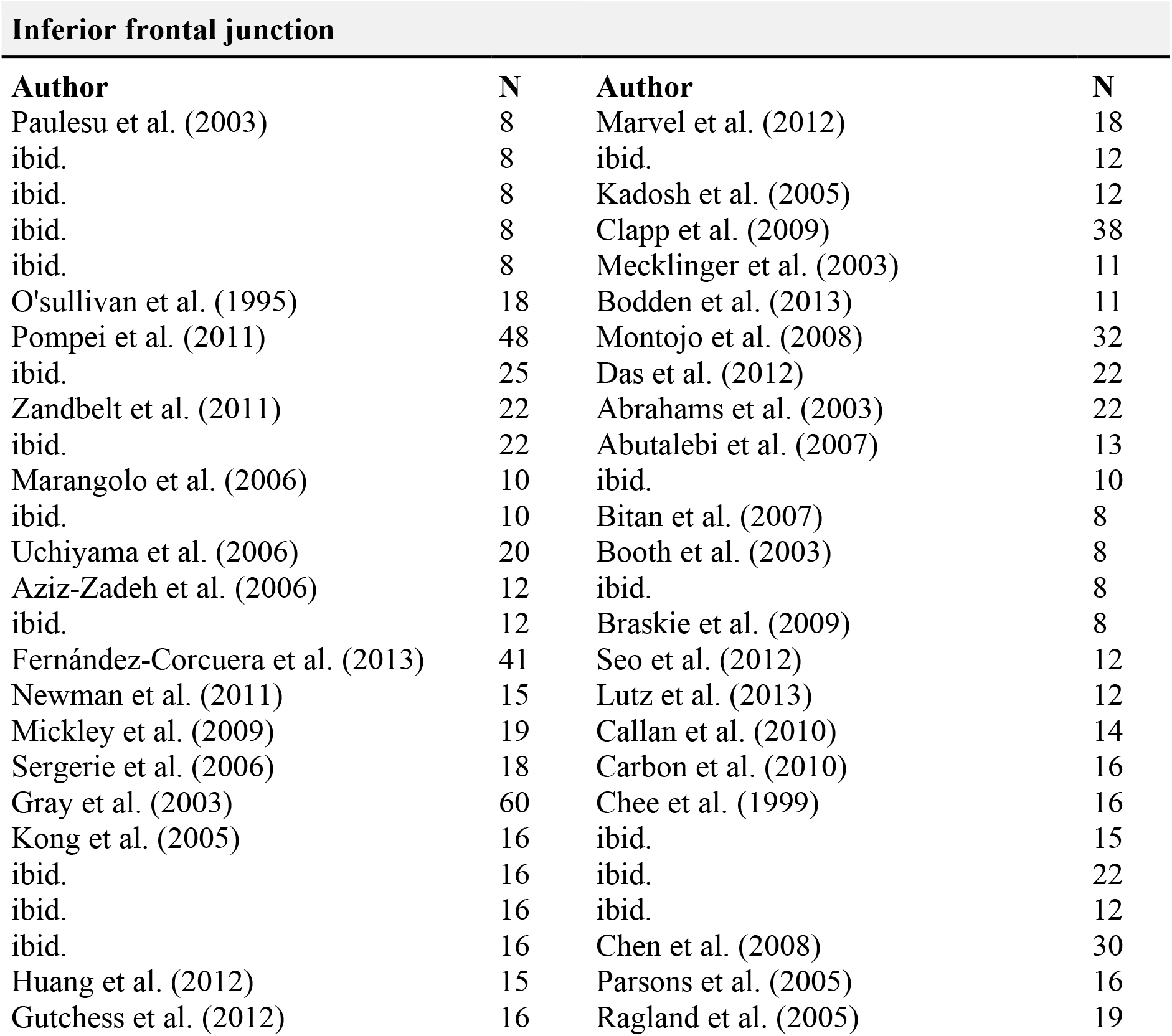

## Footnotes

1 Recall that simulation runs were discarded if ICA yielded negative weights.

2 Note that this is after adding back the mean signal removed during the ICA de-meaning step.

